# Painting Peptides with Antimicrobial Potency through Deep Reinforcement Learning

**DOI:** 10.1101/2025.01.13.632671

**Authors:** Ruihan Dong, Qiushi Cao, Chen Song

## Abstract

In the post-antibiotic era, antimicrobial peptides (AMPs) serve as ideal drug candidates for their lower likelihood of inducing resistance. Computational models offer an efficient way to design novel AMPs. However, current optimization and generation approaches are tailored for different application scenarios. To address this challenge, we propose a novel AMP design model named AMPainter. Based on deep reinforcement learning, AMPainter integrates both optimization and generation tasks in a unified framework. We apply AM-Painter to three types of peptides, including known AMPs, signal peptides (SPs), and random sequences. AMPainter outperforms ten related models in enhancing the activity of known AMPs, and evolves effective AMPs from membrane-active SPs with a success rate of 80%. Furthermore, several *de novo* designed AMPs from random sequences are validated along with their evolutionary paths. Therefore, AMPainter contributes to *paint* antimicrobial potency to diverse peptides, assisting in expanding the AMP sequence space and discovering novel antimicrobial agents.

## Introduction

Microbe-caused diseases threaten human health in recent years. According to the World Health Organization, drug-resistant infections result in over 700,000 deaths annually and this number may reach ten million in 2050. ^1^ There is an urgent need to develop new antimicrobial drugs, and one of the most attractive candidates is antimicrobial peptide (AMP). These peptides are well-known for their membrane-disrupting mechanisms and broad-spectrum antimicrobial properties, which makes them less likely to induce resistance in comparison with conventional antibiotics. ^2,3^ Even though there are more than 30,000 AMPs documented in existing databases, ^4^ they remain sparsely distributed across the vast sequence space and warrant further exploration.

Computational models provide an efficient way to discover novel AMPs and enhance their properties. Generally, AMP design works can be categorized into two types, optimization and generation. ^5^ Optimization strives to improve the antimicrobial potency of a known AMP when perturbing its sequence, while generation does not depend on available sequences and creates AMPs from scratch. Common optimization approaches include rational design ^6,7^ and evolutionary methods, ^8,9^ heavily relying on expert knowledge and exploring sequences within a limited space. ^10^ Data-driven generative models are facilitated by deep learning techniques such as the variational autoencoder (VAE), ^11–13^ generative adversarial network (GAN), ^14,15^ diffusion model^16^ and so on. These models can efficiently generate hundreds of new potential AMPs within seconds. Nevertheless, external filtering criteria are adopted to select the most promising sequences, only some of which will undergo experimental validation. ^17,18^ Particularly, some generative models can perform optimization tasks in an ‘analog generation mode’ to improve the antimicrobial potency of input sequences. ^5^ For instance, HydrAMP and deep-AMP generate analogs of specific AMPs based on VAE architecture. However, they cannot perform unconstrained and analog generation under the same training settings. ^19,20^

Classic cases of directed evolution inspired us to achieve the above two design tasks in a unified strategy. ^21^ By applying appropriate evolutionary pressure as guidance, proteins can be pushed to enhance their original functions or acquire new capacities through point mutation. ^22^ Hence, a possible solution for integrating the optimization and generation of AMPs is to develop a design strategy based on virtual directed evolution, for which reinforcement learning can be an effective way, especially for the identification of the mutation sites. Notably, a few studies have used reinforcement learning in the field of protein engineering to explore a wide search space (20*^L^*, where *L* is the length of the input sequence). ^23,24^ However, its application in *de novo* sequence design remains limited, as the starting position in the fitness landscape is often far from the ideal destination, necessitating an exhaustive search.

In this study, we propose an AMP design model named AMPainter. We split the mutation process into two steps, selecting the mutation site and assigning the residue type of the mutant. AMPainter paints antimicrobial potency to different sequences in an efficient manner with the aid of reinforcement learning and a protein language model. We utilized AMPainter to tackle three AMP design tasks, including enhancing the activity of known AMPs, evolving novel AMPs from signal peptides (SPs), and generating *de novo* AMPs from random sequences. In the optimization task, AMPainter outperformed 10 related models on a set of known AMPs. Several peptides demonstrated a 128-fold increase in antimicrobial activity *in vitro* after evolution. Experimental tests of the top 10 sequences evolved from membrane-active SPs exhibited a success rate of 80%. As for generation, a *de novo* designed AMP R04 had an average minimal inhibitory concentration (MIC) value of less than 3 µM against four bacteria. Meanwhile, AM-Painter can provide the evolutionary paths of evolved AMPs, benefiting in-depth investigations in sequence-activity relationships.

## Results

### Overview of AMPainter

Our model was named *AMPainter* for painting antimicrobial potency to any given peptide sequences. As illustrated in Fig. 1, AMPainter comprised three modules. First, we implemented a policy network to assign the mutation sites of the input sequences. This two-layer neural network transformed amino acid letters into probabilities, from which we sampled to designate a mutation site and substituted it with a masking character. Subsequently, we fine-tuned a protein language model Ankh ^25^ with AMP sequences and utilized it to decode the masked residues. Finally, we trained a novel and independent antimicrobial activity predictor Hyper-AMP to evaluate the mutated sequences. This evaluation score was then used as the reward for updating the policy network.

**Figure 1:**
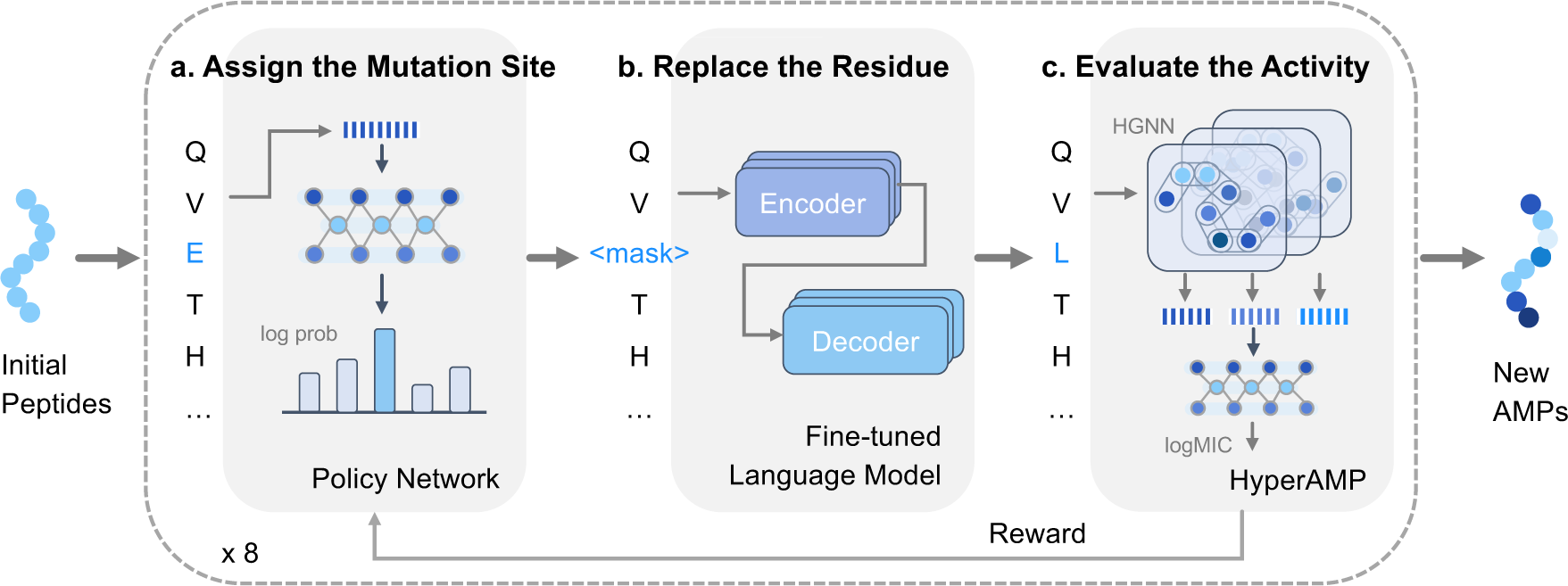
The framework of AMPainter. With a set of peptide sequences as inputs, AMPainter can evolve them into new AMPs by iteratively performing three steps: assigning the mutation site with a policy network (**a**), replacing the residue with a fine-tuned language model (**b**), and evaluating the antimicrobial activity with a predictor named HyperAMP (**c**). HyperAMP was a hypergraph neural network model, which was trained with known AMPs and their real antimicrobial activity labels before being incorporated into AMPainter. The predicted antimicrobial scores were used as rewards to update the policy network by reinforcement learning. Each peptide was processed with eight iterations.

We developed the regression model HyperAMP using multi-level hypergraph neural networks (framework in Fig. S1). In a peptide hypergraph, we treat residues as nodes. Unlike graph, a hyperedge in a hypergraph can connect more than two nodes, which is advantageous for capturing higher-order connections within the hypergraph. Therefore, we truncated sliding fragments of peptides and encoded them as hyperedges. We extracted the residual embeddings from the Ankh model as node features since it demonstrated better performance than other protein language models (Table S1). During the training of HyperAMP, we transformed the MIC labels of AMPs into reward scores (Fig. S3a). HyperAMP achieved a Pearson correlation coefficient of 0.923 and an RMSE of 0.163 on the test set, surpassing the performance of other baseline models (Fig. S2), ensuring its accurate guidance in AMPainter.

We trained AMPainter for 40 episodes, with 966 random sequences whose lengths are in the range between 10 and 40. The average reward score improved significantly, increasing from about 0.26 to 0.78. The ablation study further validated the role of bridging fine-tuned language model (Fig. S3b). AMPainter processed eight iterations for each input sequence, i.e. mutating eight steps in total and conducting one mutation for each step. We set this iteration number to achieve a balance between the antimicrobial score and diversity, as a higher number would increase the antimicrobial score but reduce the diversity (Fig. S4).

### AMPainter outperforms other methods on AMP optimization

After finishing training AMPainter, we compared its performance with 10 related methods on the task of enhancing the activity of known AMPs. The input data was 200 AMPs from the test set of the HyperAMP predictor, ensuring that these sequences had not been encountered by the AMPainter model. For the HydrAMP method, We selected its analog generation mode. ^19^ For other evolutionary approaches, we utilized HyperAMP as a surrogate model to guide the optimization process. To maintain consistency with AMPainter, we set the maximum number of mutations per sequence as eight for all methods. The top 200 sequences sorted by HyperAMP were retained for each method.

We used three types of evaluation metrics to compare the results. First, we re-scored the antimicrobial activity of the top 200 evolved sequences by the MBC-Attention model. ^26^ Since our HyperAMP had been used in optimization, we chose another scoring model here. The results showed that all 11 methods could improve the initial AMPs, and AMPainter achieved the best performance in terms of both the highest value and overall distribution (Fig. 2a). Second, we adopted six widely-used AMP classifiers (AMPScannerV2, ^27^ CAMPR4-RF, CAMPR4-SVM, CAMPR4-ANN, ^28^ MACREL, ^29^ and ampir^30^) to evaluate these evolved sequences. We calculated the average predicted probabilities for each sequence and plotted their distribution in Fig. 2b, instead of the binary labels. AMPainter outperformed other methods with overall higher probability values. Most sequences exhibited an average antimicrobial probability exceeding 0.9.

**Figure 2:**
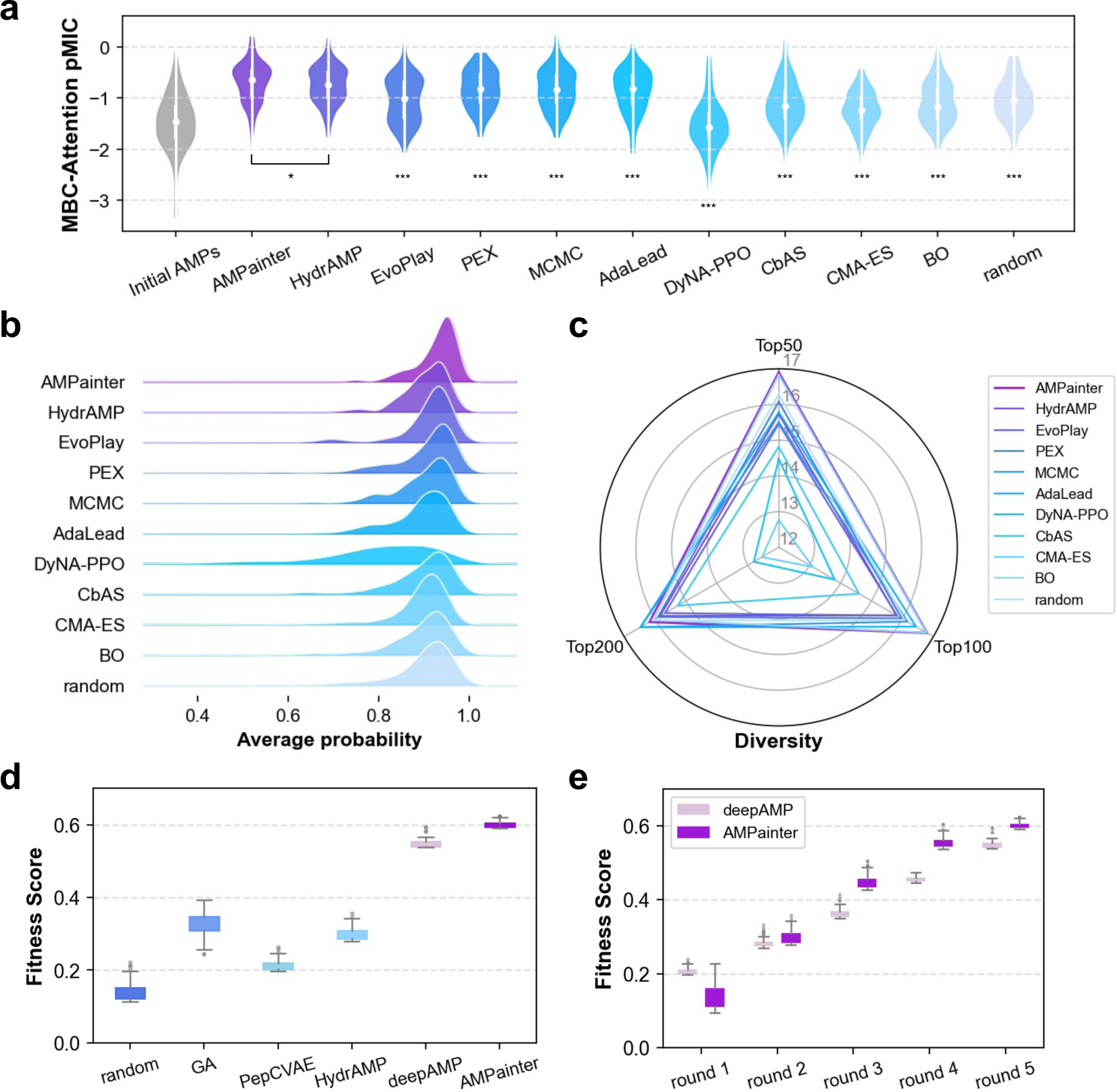
Results of AMPainter in comparison with other related methods on enhancing the activity of known AMPs. **a**, Antimicrobial evaluation of top 200 sequences optimized by various methods, scored by MBC-Attention model. One-sided Mann-Whitney test was used for statistical analysis. \**p* < 0.05. \*\*\**p* < 0.001. **b**, Distribution of average antimicrobial probability of top 200 sequences, as predicted by six AMP classifiers. **c**, Sequence diversity of top 50, 100, and 200 evolved sequences. **d**, Optimization of four Pg-AMP1 fragments with the guidance of fitness score. **e**, Comparison of AMPainter and deepAMP models on five rounds of Pg-AMP1 optimization task, with the fitness score as reward.

In addition to antimicrobial evaluations, we also compared the sequence diversity (Eq. (11)) of the top 50, 100, and 200 sequences evolved by each method in Fig. 2c. AMPainter ranked first for both the top 50 and top 100 sequences, and it secured third place for the top 200. Therefore, AMPainter can obtain a variety of optimized sequences across a range of inputs, rather than being trapped in the local optima of certain sequences.

Inspired by deepAMP, ^20^ we also investigated the capability of AMPainter in optimizing amphipathic helical peptides by using a fitness score as the reward. Starting with four fragments of Pg-AMP1 peptide, AMPainter contributed to the highest fitness score of 0.624 after five rounds of evolution, outperforming random optimization, genetic algorithm (GA, reported top 100 sequences ^9^), and three VAE models^31^ (Fig. 2d and Fig. S5a). Fig. 2e showed that the top 100 sequences evolved by AMPainter surpassed those of deepAMP from rounds 2 to 5. Evolution of AMPainter converged at significantly amphipathic sequences full of alternating leucine (L) and arginine (R) residues following the fitness score (Fig. S5b-d). These results demonstrate that AMPainter excels in optimizing AMPs.

### AMPainter enables to evolve diverse sequences into AMPs

We applied AMPainter to evolve three different sets of initial sequences, known AMPs, signal peptides (SPs), and random sequences (Fig. 3a). Enhancing the activity of known AMPs was an optimization task, for which we compared AMPainter with other approaches in the previous section. The second task was to evolve AMPs from bacterial signal peptides since their membrane-active feature may be related to the membrane-disrupting mechanisms of AMPs. The third task focused on designing AMPs from random sequences, which could be regarded as *de novo* generation. We used 200 known AMPs, 708 SPs, and 200 random sequences as input to AMPainter, respectively (for details, see the Methods section).

**Figure 3:**
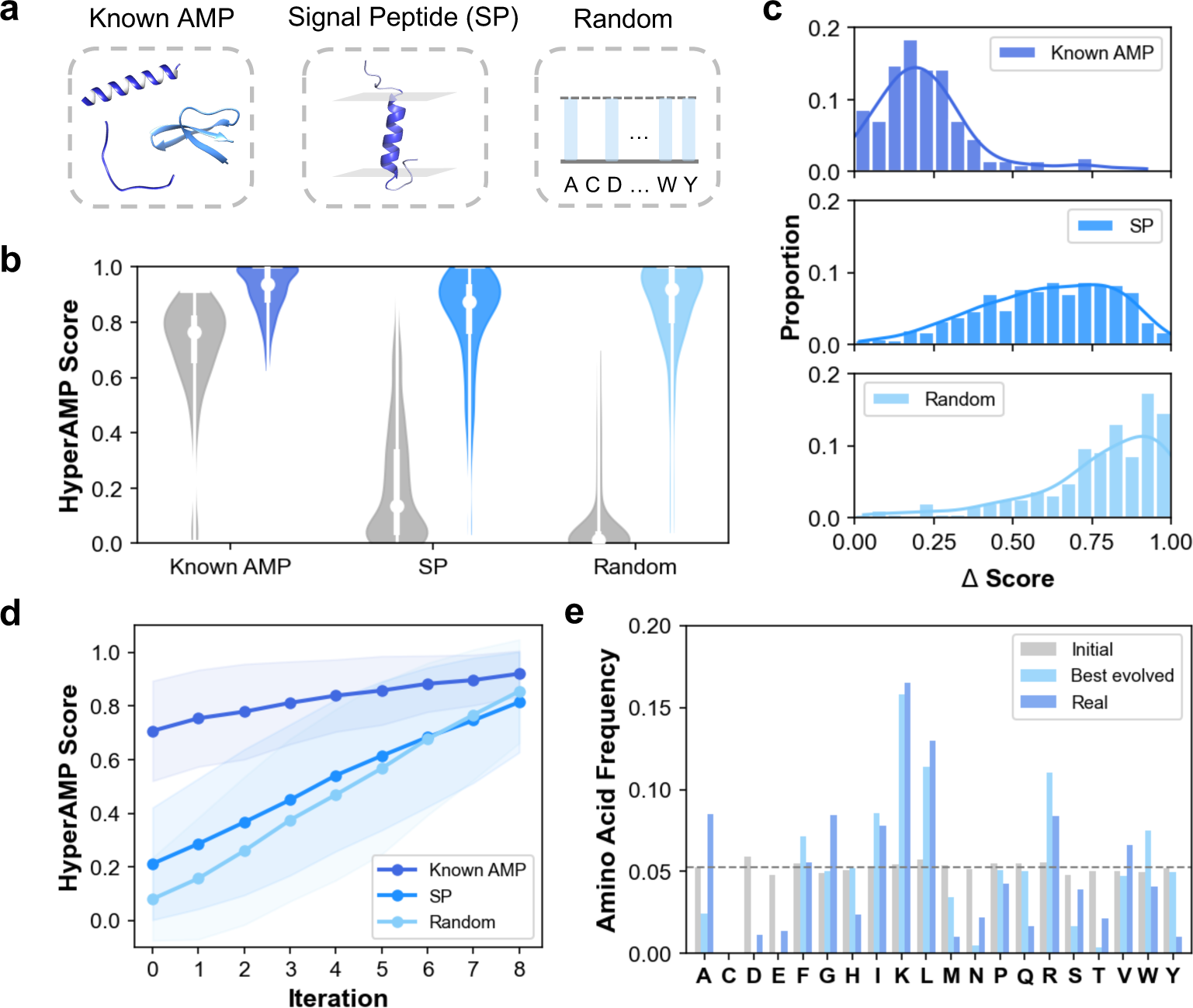
Applications of AMPainter. **a**, Three initial sets for AMPainter are known AMPs, bacterial signal peptides (SP), and random sequences. **b**, Overall evolutionary results of three sets by AMPainter. The score distributions of initial sequences are shown in grey, while the best-evolved ones are in blue. **c**, Distribution of the increasing scores of each initial sequence in three sets. **d**, Average scores of sequences at each evolutionary iteration. **e**, The amino acid frequency of initial random sequences, their best-evolved sequences, and real AMPs.

After eight iterations of evolution, we selected the best-evolved sequence of each initial sequence and investigated their antimicrobial scores. Fig. 3b displayed the overall score distributions of both the initial and best-evolved sequences in gray and blue, respectively. Scores of initial known AMPs were relatively higher than the other two sets, while some signal peptides also exhibited high scores. Almost all of the random sequences were scored around zero, indicating no antimicrobial potency. The scores of best-evolved sets increased significantly compared to the initial sets. Meanwhile, we calculated the increasing amount (Δ score) of each peptide (Fig. 3c). All the sequences had improvement with a Δ score > 0, which verified the ‘painting’ ability of AMPainter. Since the initial known AMPs already had high scores, their Δ scores were relatively smaller than the other two sets. About 60% of the sequences evolved from SPs obtained Δ scores exceeding 0.5, as did the random sequences. We also divided the increasing scores into each iteration. In Fig. 3d, the average scores of sequences in three sets grew consistently over eight iterations. For each sequence, the Δ score varied at each iteration, but the majority showed an increase, with random sequences improving at a faster rate (Fig. S6a).

Fig. 3e illustrated the amino acid frequency distribution of initial random sequences, best-evolved sequences after AMPainter’s evolution, and real AMPs. The initial sequences were generated randomly and thus exhibited a uniform distribution excluding cysteine (C), i.e., the 19 amino acids appeared at essentially the same frequency (dashed line in Fig. 3e). After our virtual evolution, the amino acid distribution of best-evolved sequences was similar to that of real AMPs, which showed a preference for positively charged lysine (K) and arginine (R). Negatively charged aspartic acid (D) and glutamic acid (E) were almost entirely replaced. Some hydrophobic amino acids like leucine (L) and isoleucine (I) were more prevalent in these two sets as well. Likewise, evolved sequences from known AMPs or SPs featured similar distributions (Fig. S6b). These results demonstrated that AMPainter could evolve different types of peptides to reach high antimicrobial potency, regardless of their initial amino acid probability distribution.

### Evolved AMPs obtain enhanced activities in vitro

To verify the antimicrobial activity of evolved AMPs, we measured their minimum inhibitory concentration (MIC) values *in vitro*. We selected 30 AMP candidates for chemical synthesis and MIC measurement, involving the top 10 peptides from the three evolved AMP sets discussed above. We didn’t include any additional metrics to filter the sequences for entering the experimental validation stage. According to their initial sequences and antimicrobial scores, we named them A01 to A10 (from known AMPs), S01 to S10 (from signal peptides), and R01 to R10 (from random sequences). All the peptides were positively charged and some featured a high hydrophobic ratio (Table S2). Meanwhile, we checked the similarity of these candidates using BLAST against the largest AMP database, DRAMP. ^4^ Except for the peptides that evolved from known AMPs, most of them had E-values greater than 1 or could not find any matching sequences (labeled ‘/’ in Table S3), highlighting the novelty of the selected peptides.

We chose four standard bacteria for the MIC test, including two Gram-positive strains (*B.subtilis* ATCC6633 and *S.aureus* ATCC6538) and two Gram-negative strains (*E.coli* ATCC25922 and *P.aeruginosa* ATCC9027). Fig. 4 displayed all the test results. We regarded a peptide as an AMP if it exhibited a MIC of less than or equal to 128 µM against at least one bacterial strain. ^32,33^ All peptides evolved from known AMPs met this criterion and retained their antimicrobial potency (Fig. 4a). As for peptides evolved from SPs (Fig. 4b), eight out of ten were identified as AMPs, except for S09 and S10, indicating a success rate of 80%. For peptides evolved from random sequences, the success rate was 60%, with R01-R05 and R09 demonstrating antimicrobial activity (Fig. 4c). Among them, R04 had an outstanding performance with an average MIC of less than 3 µM, which was superior to most of *de novo* generated AMPs. ^13^ Overall, the AMPs got lower MICs against Gram-positive strains compared to Gram-negative strains. This might be attributed to the structural variations in their membranes, as Gram-positive bacteria lack the outer membrane.

**Figure 4:**
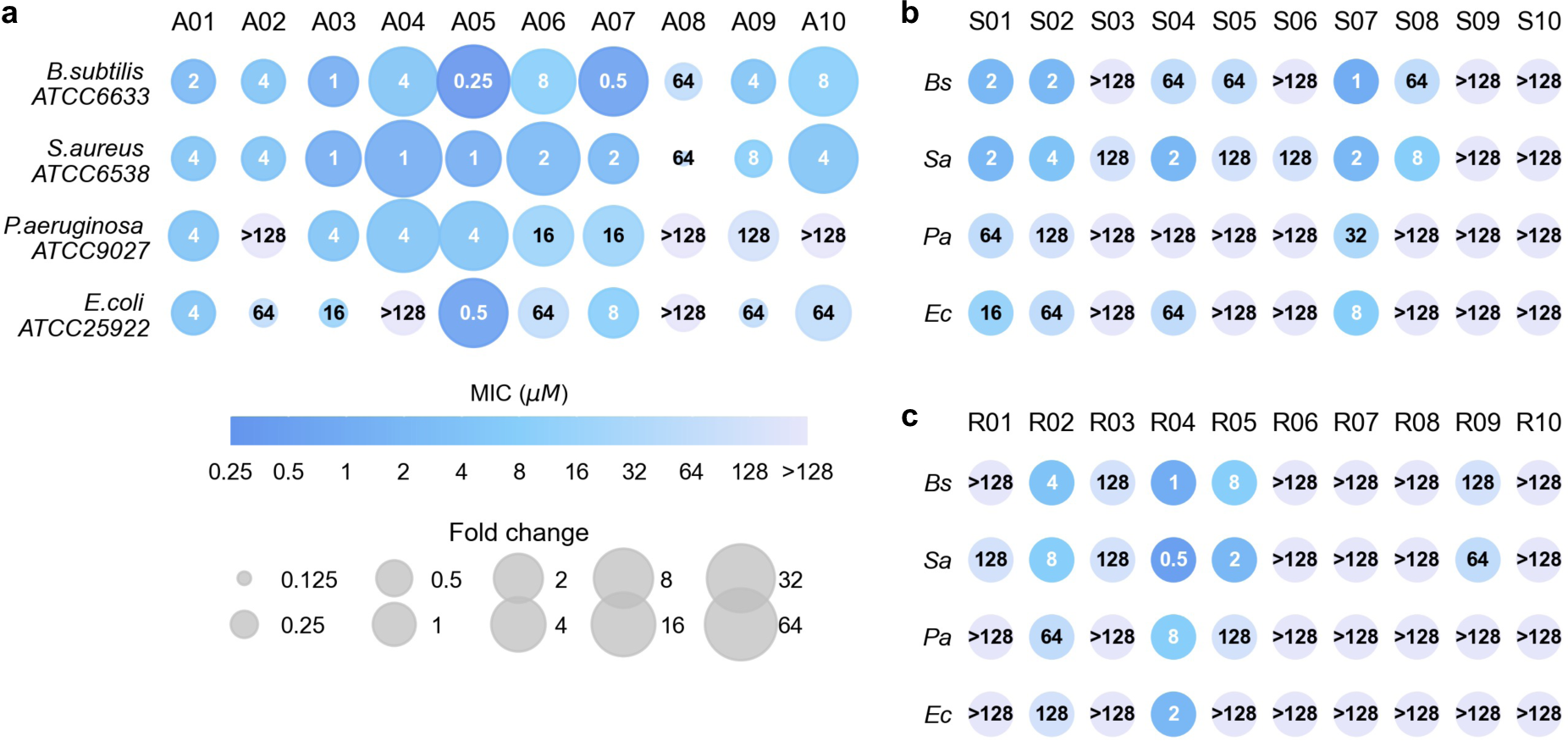
Minimal inhibitory concentrations (MIC) of evolved peptides against four bacterial strains. **a**, Top 10 sequences (A01-A10) evolved from known AMPs. The area of scatter shows the fold changes of evolved sequences compared to their corresponding initial sequences. **b**, Top 10 sequences (S01-S10) evolved from bacterial signal peptides. **c**, Top 10 sequences (R01-R10) evolved from random sequences. *Bs*, *B.subtilis* ATCC6633. *Sa*, *S.aureus* ATCC6538. *Pa*, *P.aeruginosa* ATCC9027. *Ec*, *E.coli* ATCC25922.

For A01-A10, we also tested the MICs of their corresponding initial AMPs (results in Table S4) and calculated the fold changes after being optimized by AMPainter (Fig. 4a). Here we measured the MICs of initial AMPs under the same conditions with A01-A10 to ensure the values were comparable. The MICs of eight peptides improved against at least one bacteria, and nine MICs of five peptides increased more than 16 times. For example, A05 peptide obtained an MIC of 0.25 µM against *B.subtilis*, and this value of its original AMP Ascaphin-5 was 16 µM, which indicated that the antimicrobial potency of A05 increased by 64 times after the optimization. A05 performed remarkably on the other three strains as well, with MICs of 1, 4, and 0.5 µM, corresponding to fold changes of 32, 32, and 4. We also checked the predicted scores of initial sequences of S01-S10 and R01-R10 by HyperAMP, and all scores were below 0.1, showing no potential for these peptides to be AMPs. Therefore, AMPainter accomplished the tasks of evolving new AMPs from a diverse range of sequences, enabling both the optimization and *de novo* generation of highly active AMPs.

### Evolved AMPs preferentially target bacteria membranes

Hemolysis and cytotoxicity are important factors to check in AMP design, as they can influence the potential of AMPs as drug candidates. Due to the membrane-disrupting mechanism of most AMPs, they can often damage the plasma membrane of human cells. We investigated the concentration causing 25% hemolysis of rat erythrocytes (HC_25_) and the concentration reducing the viability of human embryonic kidney 293T (HEK293T) cells by 50% (CC_50_) of the 24 AMPs evolved by AMPainter (Fig. 5a, details in Table S5). We found that several evolved AMPs from known AMPs showed hemolytic and cytotoxic tendencies. Notably, these attributes were also observed in some initial AMPs as previously reported (Table S4). Meanwhile, we harvested some AMPs with ideal hemolysis and cytotoxicity, particularly those evolved from signal peptides. All HC_25_ values of S01-S08 exceeded 128 µM, indicating that bacterial signal peptides may preferentially interact with bacterial membrane rather than human membrane.

**Figure 5:**
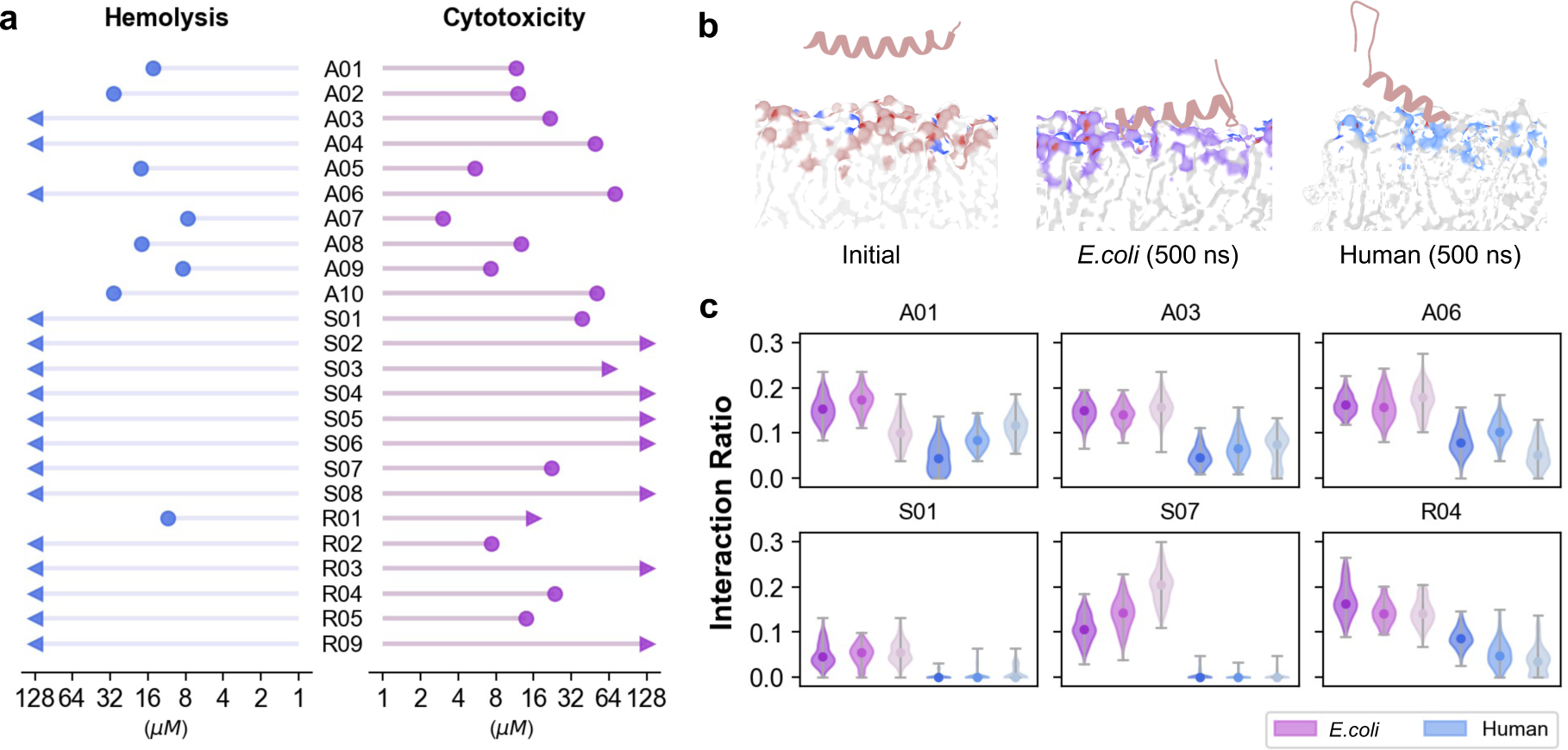
Hemolysis, cytotoxicity and membrane preferences of evolved peptides. **a**, Hemolysis (HC_25_) on rat blood and cytotoxicity (CC_50_) on HEK293T cells of AMPs evolved by AM-Painter. **b**, Representative snapshots of MD simulations, with *E.coli* inner membrane or human plasma membrane models. **c**, Interaction ratios of AMP heavy atoms with membrane heavy atoms (with a cutoff of 3.5 Å).

We also calculated the selectivity index (SI) of the 24 AMPs by dividing the mean MIC by HC_25_ or CC_50_ (Table S5). An AMP was likely to function as an antimicrobial agent with reduced side effects when its SI exceeded one. Hence, some of our AMPs held the capability to distinguish different membranes such as A03 (SI > 23.27), S07 (SI > 11.91), and R04 (SI > 44.44). To further verify their membrane-targeting preferences, we performed molecular dynamics (MD) simulations on six peptides with relatively high SI and CC_50_. We built two systems with distinguished compositions of lipid bilayers ^13^ for each peptide, mimicking the inner membrane of *E.coli* and human plasma membrane (Table S6), respectively. The peptides were placed 2 nm above and parallel to the membranes at the beginning of the simulations, and showed different interactions with the membrane surface after 500 ns of simulation, as shown in Fig. 5b. The significant difference was that these peptides were completely adsorbed to *E.coli* membrane, while they only contacted with human membranes with one terminus or even no contact at all (S01 and S07). We counted the ratio of heavy atoms in peptides interacting with membranes (cutoff 3.5 Å) in the last 100 ns (Fig. 5c). These six peptides exhibited a higher interaction ratio with the *E.coli* membrane than the human membrane, particularly for S01 and S07. The interaction ratio calculated by frame counting showed similar results (Fig. S7). This preference appears to be caused by electrostatic interactions, since bacterial membranes had more negative charges than human membranes and tended to interact with positively charged AMPs. A similar phenomenon was observed by previous studies as well. ^13^ The specific mechanisms warranted further exploration through larger-scale simulation and wet-lab experiments.

### Evolutionary paths of a de novo generated AMP

To investigate how AMPainter evolved the peptides, we analyzed the improvement in antimicrobial potency over the eight steps of evolution. Specifically, we focused on the task of evolving AMPs from random sequences and regarded it as *de novo* generation. After validating *de novo* designed AMPs *in vitro* as above, we found that R04 presented attractive antimicrobial activity and selectivity. The 26-residue R04 is an amphipathic *α*-helix peptide, representing a successful case of generation with typical AMP characteristics (Fig. 6a, b).

**Figure 6:**
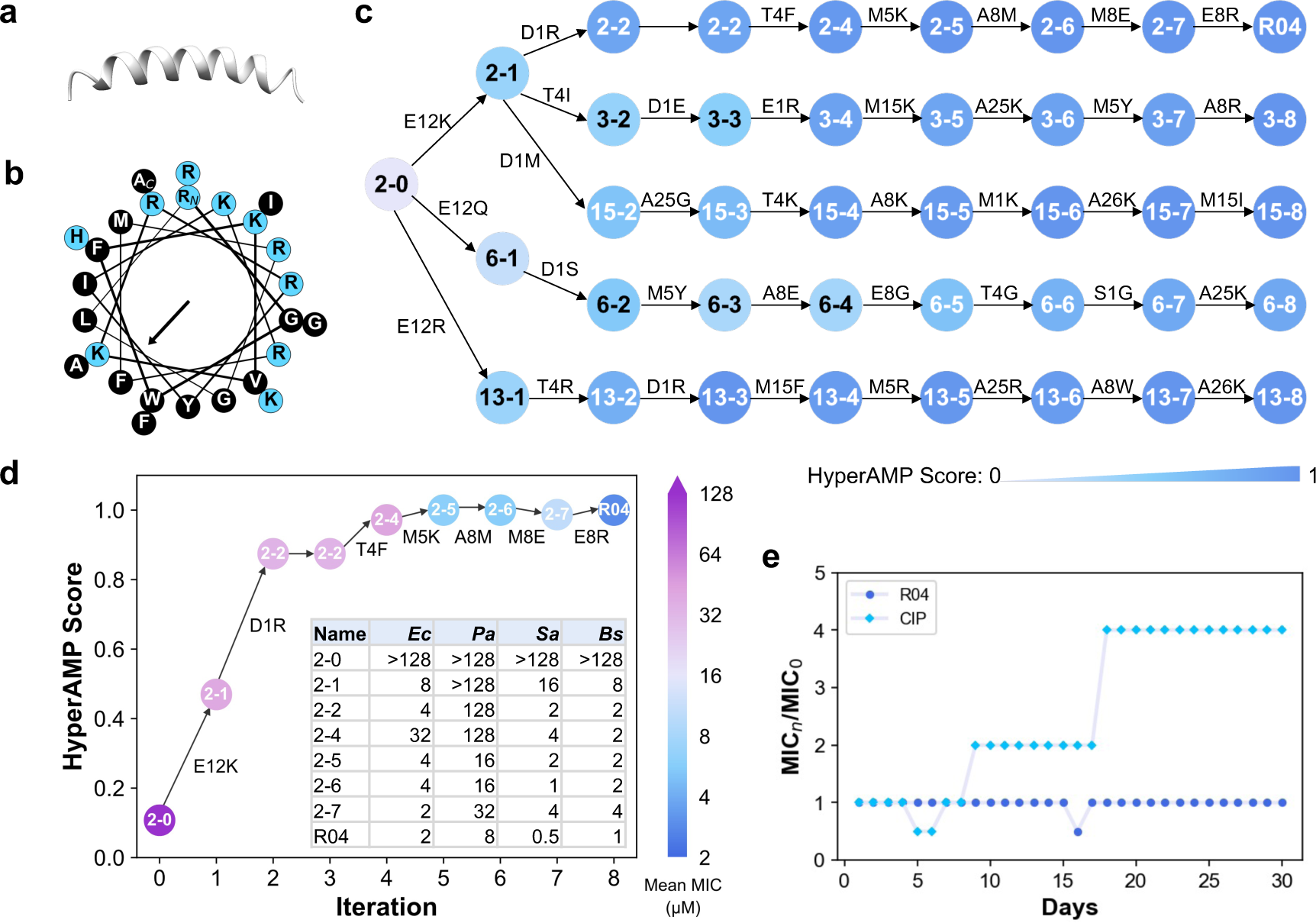
Characterizations of a *de novo* designed AMP R04 from AMPainter. **a**, Structure of R04 predicted by AlphaFold 2. **b**, Helical wheel of R04. Blue and black stand for polar and non-polar residues, respectively. **c**, Five parallel *in silico* evolutionary paths of R04 from AMPainter. Peptide names are labeled as *path-iteration*, and are in black with HyperAMP score of less than 0.6 for clarity. **d**, The predicted HyperAMP scores of sequences in the R04 path match experimentally measured mean MIC values. *Ec*, *E.coli* ATCC25922. *Pa*, *P.aeruginosa* ATCC9027. *Sa*, *S.aureus* ATCC6538. *Bs*, *B.subtilis* ATCC6633. **e**, The resistance test of R04 in 30 days. CIP, ciprofloxacin.

We took a deeper insight into R04 by collecting its evolutionary paths from AMPainter. Fig. 6c displayed five parallel paths starting from the same sequence 2-0. AMPainter produced 20 paths for 2-0 in total and the top five, whose ending sequences had higher activity, were presented here. In other words, all the peptides shown in Fig. 6c were the homologous sequences of R04. As the number of mutations increased, the activity score elevated along all five paths, and rose rapidly during the first four iterations. In terms of mutation types, K and R appeared multiple times to increase positive charges, which was consistent with the common features of AMPs. When aligning different paths, we found some conserved mutation sites. For instance, the negatively charged E12 of 2-0 was mutated in all paths at the first iteration. The replacement of M5, A8, and A25 also occurred multiple times, although their mutation orders were not the same.

Furthermore, we concentrated on the evolutionary path of R04 from 2-0 (Fig. 6d). According to their predicted antimicrobial score, 2-0 was a non-AMP, and transformed to 2-1 and 2-2 by replacing two negatively charged residues with positively charged residues. Then AM-Painter mutated T4F of 2-2 and M5K of 2-4 with scores increasing slightly. There were consecutive mutations of A8 during the last three iterations, including stepwise mutations to M, E, and R. This indicated that AMPainter did not always add positive charges to generate AMPs, and sometimes took detours, as the score slightly decreased at 2-7. We also experimentally tested the MICs of all eight sequences in this path. Impressively, the mean MICs against four bacteria were consistent with the predicted scores, exhibiting a decreasing trend (higher antimicrobial activity) from 2-0 to R04, except for 2-7. MICs of 2-7 against three strains were doubled or quadrupled compared to 2-6, presenting a negative effect of M8E. After the E8R mutation, R04 obtained the best MICs against all four strains. The *in silico* evolutionary paths from AMPainter assisted in exploring the sequence-activity relationships of AMPs, eliminating the need for time-consuming mutational scanning.

Furthermore, we conducted a 30-day resistance test of R04 in comparison with the antibiotic ciprofloxacin (CIP) on *E.coli* ATCC25922 (Fig. 6e). CIP began to induce resistance on the eighth passage day, whereas the continuous use of R04 did not lead to resistance. In a word, R04 is a *de novo* designed AMP with superior properties and has great potential to serve as an antimicrobial agent.

## Discussion

In the post-antibiotic era, severe drug-resistant infections necessitate the development of novel therapeutic compounds, with AMPs serving as promising candidates. Compared to the time-consuming identification of AMPs from natural resources *in vitro*, artificial intelligence significantly accelerates their discovery. For instance, hundreds of AMPs have been discovered by mining from metaproteomes. ^33,34^ Meanwhile, numerous design methods have emerged to optimize existing AMPs or generate *de novo* AMPs with various model architectures. ^5^ To integrate these two design modes and further improve the model performance, we presented AM-Painter, a deep reinforcement learning-based framework. We successfully applied AMPainter to three types of initial peptides (known AMPs, signal peptides, and random sequences), and obtained a series of novel AMPs with the guidance of an antimicrobial activity predictor HyperAMP. *In vitro* experiments demonstrated a notable success rate (80% for SP, and 60% for random sequences) for the top 10 novel AMPs.

Without restricting the types of initial sequences, AMPainter effectively broadens the AMP sequence space. Although we cannot guarantee that every initial sequence can be transformed into a highly active AMP within limited mutation steps, the optimization directions are aligned with increasing reward scores, as demonstrated by the ‘painting’ capability of AM-Painter. During the evolution process, we did not incorporate complex rewards beyond the antimicrobial potency of peptides. Also, we only took this data-driven predictive metric into account when selecting evolved peptides for experimental validation, thereby reducing reliance on expert knowledge.

Another advantage of AMPainter is that it can provide a set of parallel evolutionary paths when iteratively adding mutations to an initial sequence. In particular, we have verified the consistency between the actual and predicted antimicrobial potency along the evolutionary path of the *de novo* designed AMP R04. Typical charge-based mutations were observed along this path. We propose that these sequences derived from parallel paths can function as artificial evolutionary profiles, potentially serving complementary roles to actual multiple sequence alignments (MSA) in related tasks.

There are also some limitations of AMPainter. The current model only supports one-point substitution at each iteration and excludes multi-point substitution, insertion, and deletion operations. This limitation restricts the evolution space of each sequence to some extent. Additionally, hemolysis and cytotoxicity are not considered during our virtual evolution, which is a challenging task that needs to be addressed. Also, it is still required to conduct thorough studies on the function mechanisms of the generated AMPs. Recent studies have investigated combining data from simulations or wet-lab assays with machine learning tools to assess the membrane selectivity of peptides. ^35,36^ How AI techniques can push the boundaries of mechanism-driven design of AMPs remains to be explored. We anticipate developing such design strategies in the future.

## Methods

### Data curation

#### HyperAMP dataset

The training, validation, and testing datasets for HyperAMP contained AMPs and non-AMPs. The AMP dataset was collected from PepVAE ^11^ with their minimal inhibitory concentration (MIC) labels against *E.coli*. Peptides with chemical modifications and disulfide bonds were excluded to avoid their effects on activity. Since HyperAMP encoded a sequence via sliding windows of 2-, 3-, and 4-gram fragments, peptides whose lengths are less than five were also filtered out. The dataset included 3,265 AMPs in total. Non-AMPs with the same amount were selected from short peptides in Uniprot without antimicrobial-related annotations, as processed by SenseXAMP, ^37^ and were labeled with zero. The entire dataset contained 6,530 peptides with a length distribution of [5,40] and was split as the training, validation, and testing sets in a ratio of 8:1:1 randomly. The output score of HyperAMP was transformed from logMIC values following the Eq. (1) below (Fig. S3a).

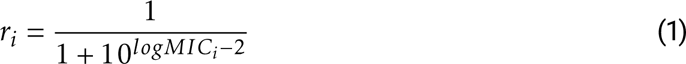

where *r_i_*is the reward score of peptide *i*.

#### Fine-tuning dataset

To fine-tune the Ankh-base model on sufficient AMPs, 9,149 non-overlapping AMPs were collected from 11 AMP databases, including APD3, ^38^ DRAMP, ^4^ and DBAASP, ^39^ etc. ^40^ 7,316 sequences remained after removing redundancy via CD-HIT ^41^ with a cutoff of 0.9. The length of this fine-tuning set ranged from 10 to 100 amino acids, with no further restrictions. Following the training settings of the Ankh-base model during its pretraining stage, the masking ratio was set as 20%, meaning that 20% of the residues in each sequence were replaced with masking tokens.

#### Policy network training dataset

Random sequences were used to train the policy network of AMPainter. A thousand random sequences were produced with length in [10, 40] and ensured no overlap existed between these sequences and the other two mentioned AMP datasets (HyperAMP and fine-tuning datasets). Then CD-HIT ^41^ was also employed to remove redundancy with a stricter cutoff of 0.7 to maintain the sequence diversity. Finally, this dataset comprised 966 random sequences for training the policy network.

#### Initial evolving datasets

We chose three sets of initial sequences and evolved them by AMPainter, which included known AMPs, signal peptides, and random sequences.

##### Known AMPs

Part of the AMPs from the test set of HyperAMP were used as initial sequences for optimization, as they were not seen by AMPainter during the training process. To compare AMPainter with the analog generation mode of HydrAMP, ^19^ those AMPs with lengths of less than 25 residues were kept to meet HydrAMP’s limit. Then the remaining AMPs were ranked by their actual MIC labels and the top 200 AMPs with MIC values exceeding 10 µM were selected.

##### Signal peptides

Signal peptides are membrane-active protein fragments. 4,664 signal peptides from the training set of SignalP 6.0^42^ were considered. The signal peptides derived from bacteria (including both Gram-positive and Gram-negative) with lengths in [10,30] were kept, considering the membrane selectivity of evolved AMPs. Then peptides containing cysteine (C) or three identical amino acids in a row were excluded due to synthesis difficulty. ^19^ The first methionine (M) of each peptide was also removed, as it was translated from the start codon during expression and had no relation with membrane-active function. As a result, 708 bacterial signal peptides were kept for modification.

##### Random sequences

The number of random peptides to be evolved was set as 200, with lengths ranging in [10, 30] as well. To prevent the formation of disulfide bonds and its effect on peptides, cysteine was excluded from the amino acid composition and the remaining 19 types of amino acids were used with a balanced frequency distribution. No sequences contained three identical amino acids in succession.

### AMPainter model

#### HyperAMP predictor

HyperAMP is a regressor which predicts the antimicrobial score of the inputting peptide sequence. A peptide was split into sliding fragments with different lengths (2-, 3-, 4-gram). Each peptide was encoded as a hypergraph *H* =*< V, E >*, where residues were nodes (*V*) and fragments were hyperedges (*E*). Residue-level node features were 768-dimension embeddings from pretrained language model Ankh-base, ^25^ and hyperedge weights were term frequency-inverse document frequency (TF-IDF) values of corresponding fragments. Hypergraphs of each level were fed into a hypergraph neural network (HGNN), ^43^ an expansion of the graph convolution network. The hyperedge convolution includes two message-passing operations, from nodes to hyperedges and from hyperedges to nodes, following Eq. (2) below:

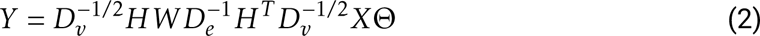

where *X* is the matrix of node features and *W* is the diagonal matrix of hyperedge weights. *H* is the incidence matrix of the hypergraph. *D_v_*and *D_e_* denote the diagonal matrices of the vertex and edge degrees, respectively. Θ is the learnable parameter during training. Output embedding *Y* of three levels were concatenated and used for regression with a 2-layer fully-connected neural network. The framework of HyperAMP was shown in Fig. S1.

TF-IDF is a commonly used encoding approach in the field of natural language processing. ^44^ TF is the emerging frequency of each word in a sentence, representing their importance. IDF is used to prevent the effects of some meaningless conjunctive words with high TF in the entire document. A peptide sequence *P_i_*was defined as a sentence consisting of *n* continuous amino acid fragments *f_i_* (Eq. (3)). The entire training set was defined as the document consisting of *N* peptides. Hence, the TF-IDF value of a single fragment *f_i_* can be calculated as Eq. (4).

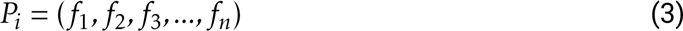

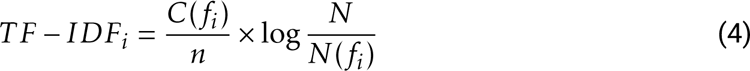

where *C*(*f_i_*) is the counts of *f_i_*in peptide *P_i_*, and *N* (*f_i_*) is the counts of peptides containing *f_i_* in the set of all peptides.

HyperAMP was developed based on PyTorch and Deep Hypergraph (DHG) libraries. ^43^ The optimizer was Adam and the loss function was the mean squared error function. The learning rate was 0.0001 and the batch size was 128. The training epoch was set to be 20 and early stopping was adopted to avoid overfitting.

#### Fine-tuned language model

The Ankh-base model was fine-tuned on AMP sequences to ensure its decoding preference. To keep up with the pretraining stage of Ankh-base, we used the same hyperparameters including a learning rate of 0.0003, a weight decay of 0.0005, and a batch size of 16. ^25^ The number of epochs was two since we observed fast convergence on this data scale. The entire process of fine-tuning was completed within 20 minutes on an NVIDIA A40 GPU. When using this fine-tuned language model to decode in the AMPainter framework, we set the temperature as two and the beam number as ten to maintain the diversity of output sequences.

#### Policy network

A 2-layer plain neural network was used as the policy network. When inputting a peptide sequence with length *L*, this policy network would output a *L* × 1 vector as their probability to be assigned as the mutation site. From the perspective of reinforcement learning, the current state is the peptide sequence, the action is to select where to mutate (and to decode the mutated amino acid by the fine-tuned language model to finish a mutation step), and the reward score is the antimicrobial score predicted by HyperAMP. The policy gradient-based REINFORCE algorithm^45^ was utilized to train AMPainter. Within the timestep *T* − 1, a training trajectory is a sequential connection of the state, action, and reward (*s*_1_*, a*_1_*, r*_1_*, …, s_T_* _−1_*, a_T_* _−1_*, r_T_* _−1_). A policy *π_θ_*(*a_t_*|*s_t_*) is the output with parameter *θ* at step *t*. For *t* in *T* − 1 timesteps, the sum of rewards *r_t_* is calculated as *G_t_* without the discount factor:

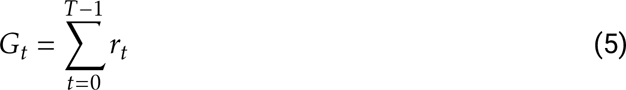

And the objective function of REINFORCE is used to update the parameter *θ*:

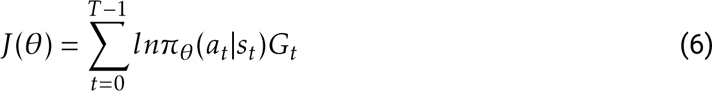

Every sequence was mutated once during each iteration, and this process was repeated eight times in an episode. The maximum number of training episodes was 40. If there was no improvement in the average reward score within 15 episodes, the training would be halted. The total training time of AMPainter on an NVIDIA A40 GPU was about 36 wall-clock hours.

### Comparative experiments

#### Comparisons of HyperAMP

Due to the lack of trainable antimicrobial activity regressors, we compared HyperAMP with six baseline models in Fig. S2. Transformer, CNN-LSTM, CNN-GRU, MLP, and CNN models were built with DeepPurpose. ^46^ Transformer took in the peptide fragments, MLP used amino acid composition (AAC), and others used one-hot encoding. As for GCN, we transformed the hyper-graph in HGNN to graph by clique expansion and kept the same node features. Hyperparameter settings of baseline models were the same as HyperAMP.

As for the embedding comparison, five pretrained language models were used (Ankh-base, Ankh-large, ^25^ ESM-1b, ^47^ ESM-2, ^48^ and ProtT5-XL-UniRef50^49^). Their embedding dimensions were 768, 1536, 1280, 1280, and 1024, respectively. The dimension of the first layer in HGNN was set in consistency of the embedding. Other model settings were kept the same.

Regression evaluation metrics including mean square error (MSE), root mean square error (RMSE), Pearson correlation coefficient (Pearson), and Spearman’s rank correlation coefficient (Spearman) are calculated as following:

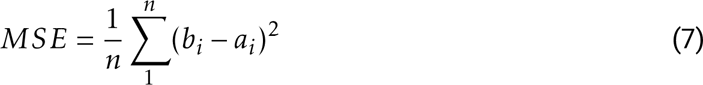

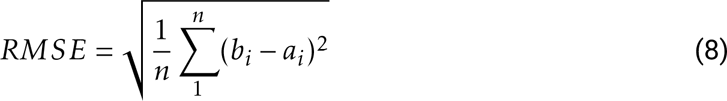

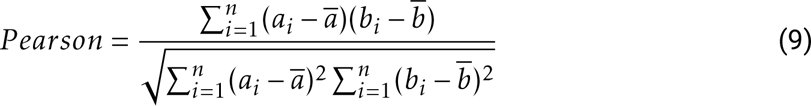

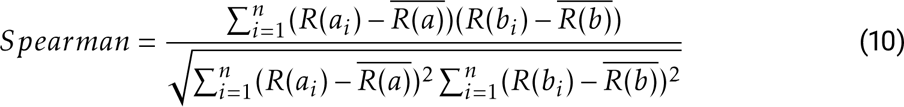

where *a_i_* and *b_i_* denote the actual and predicted values of sequence *i*, respectively. *R*(•) calculates the ranked number.

#### Ablation study of fine-tuned language model

In the ablation study, two ablating versions of the fine-tuned language model part were adopted to train the policy network (Fig. S3b). One was to directly use pretrained Ankh-base model without fine-tuning with AMPs and the other was to choose amino acid type randomly instead of incorporating the language model. All other settings were the same as AMPainter.

#### Comparing with other evolutionary methods

AMPainter was compared with other 10 related methods for AMP improvement, among which HydrAMP was an AMP generation model and others were evolutionary approaches. EvoPlay, PEX, and MCMC were explored with their original codes, while others were all implemented through the FLEXS library. ^50^ HyperAMP was added as the surrogate model for all evolutionary methods.

1. HydrAMP: ^19^ a conditional VAE model for AMP generation. Here its analog generation mode was adopted and filters were the default setting.
2. EvoPlay: ^24^ a protein engineering model based on Monte Carlo Tree Search (MCTS). It explored the mutations under the guidance of fitness functions.
3. PEX: ^51^ a proximal exploration approach. It assumed that local optima existed near the wild-type sequence and tried to minimize the number of mutations. Fitness was scored via a mutation factorization network (MuFacNet).
4. MCMC: ^52^ a protein engineering model based on the Markov Chain Monte Carlo algorithm which decided to accept mutations on the Metropolis-Hasting rule.
5. AdaLead: ^50^ a model-guided evolutionary approach that optimized query sequence via batches with a high-climbing search strategy.
6. DyNA-PPO: ^23^ a sequential decision-making model for sequence design based on model-based reinforcement learning, which used proximal policy optimization (PPO).
7. CbAS: ^53^ a method called conditioning by adaptive sampling which restricted sampling distribution in labeled dataset.
8. CMA-ES: ^54^ a statistical method called covariance matrix adaptation evolution strategy. It estimated the covariance matrix to adjust the search.
9. BO: ^55^ the traditional Bayesian optimization.
10. random: mutate randomly.

The top 200 sequences ranked by HyperAMP scores from all methods were collected for analysis. MBC-Attention^26^ was used as a validation predictor. Its predictive score was represented as the negative logarithmic value of MIC (pMIC) so a higher score indicated better antimicrobial activity (Fig. 2a). The sequence diversity in Fig. 2c was calculated as Eq. (11).

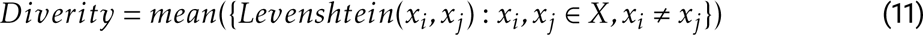

where *X* is the output set of sequences, and *x_i_*and *x_j_* are two different sequences.

#### Optimizing by fitness score

The fitness score (Eq. (12)) ^9 20^ was used as the reward to train AMPainter, instead of the HyperAMP score. Four fragments of Pg-AMP1 were evolved for five rounds using this new version of AMPainter. In each round, ten parallel evolutions and eight iterations of mutation were performed. The top 100 sequences were collected and used as initial sequences for the next round.

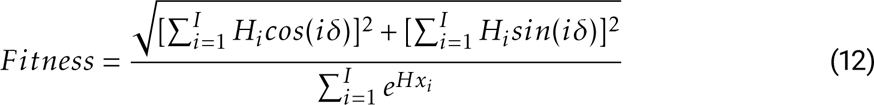

where *δ* equals to 100°, *H_i_* is the Eisenberg’s hydrophobicity of residue *i*, and *Hx_i_*is the Pace-Schols’ helix propensity of residue *i* in a sequence with length *I*.

### MD simulations

3D structures of designed peptides were predicted by AlphaFold 2^56^ (ColabFold v1.5.5^57^) with single sequence mode. CHARMM-GUI ^58^ was used to build the lipid bilayer systems with peptides parallel to the membrane surface above 2 nm. Two systems were built with different lipid compositions for each peptide to mimic both the *E.coli* inner membrane and human plasma membrane (Table S6). Ions of 150 mM NaCl were added to neutralize the system. Simulations were run with GROMACS 2018^59^ and CHARMM36m force field. ^60^ The timestep was set as 2 fs. The temperature was kept at 310 K by Nose-Hoover method and pressure at 1.0 bar with Parinello-Rahman method. The cutoff value of Lennard-Jones potential was 1.2 nm. The NVT and NPT equilibriums were conducted following the default procedures of CHARMM-GUI. All runs were produced for 500 ns with three independent repeats. The interaction ratio of peptide heavy atoms and membrane heavy atoms was calculated using MDAnalysis. ^61^

### Wet-lab validations

#### Peptide synthesis

All peptides were synthesized via solid-phase peptide synthesis by DGpeptide Co., ltd. Their molecular weights were verified by mass spectrometry and their purity (>95%) was verified by HPLC (results shown in Fig. S8 - Fig. S51).

#### Minimal inhibitory concentration measurement

Four standard bacteria used for MIC test were *Escherichia coli* ATCC25922, *Pseudomonas aeruginosa* ATCC9027, *Staphylococcus aureus* ATCC6538, and *Bacillus subtilis* ATCC6633. The testing procedures followed the Hancock methods. ^62^ Peptides were diluted in sterilized PBS to 512 µM as storage solutions. Then two-fold gradient dilution was performed to make different concentrations (128, 64, 32, 16, 8, 4, 2, 1, 0.5, 0.25, 0.125 µM) of AMPs. Each bacteria was incubated in LB broth at 37°C until its absorbance at 625 nm reached 0.08-0.10, then diluted 10,000 times and added into a sterile 96-well polypropylene plate (Grenier, #655201). A co-incubation system included 100 µL peptide solution and 100 µL bacterial suspension. After growing at 37°C for 18 hours, the lowest peptide concentration causing no observed bacteria grown was determined to be the MIC. All tests were in triplicate.

#### Resistance test

A 30-day resistance test was conducted to observe the resistance-causing trend of both AMPs and antibiotics. Ciprofloxacin (Aladdin, #C129896) was used as the control. Each day, 1 µL bacterial suspension from the wells containing the highest concentration of AMPs which enabled the growth of bacteria (1/2 current MIC) was extracted into 10 mL LB broth to dilute 10,000 times. Other procedures were the same as above to detect MICs. The concentration gradients of R04 peptide were 16, 8, 4, 2, 1, 0.5 µM. All tests were in triplicate.

#### Hemolysis assay

*In vitro* hemolysis and cytotoxicity were measured by WuXi AppTec Co., ltd. The rat blood was collected and mixed in PBS, then centrifuged at 500 g for 5 mins to make the red blood cell solution (RBC, 10%). Two-fold gradient dilution was performed to make different concentrations (128, 64, 32, 16, 8, 4, 2, and 1 µM) of AMP solutions. Then AMP solutions were added into 100 µL RBC to make assay solution and incubated at 37°C for 1 min. The assay solution was centrifuged at 2500 g for 6 mins. After centrifuging, 75 µL of supernatant was placed into a 96-well plate. PBS was used as the vehicle control and 0.1% Triton X-100 was the positive control. The absorbance of supernatant in each well was detected at 450 nm using EnVison (PerkinElmer) and the percentage of hemolysis was calculated as Eq. (13). The HC_25_ value was fitted using GraphPad Prism 8. All tests were in triplicate.

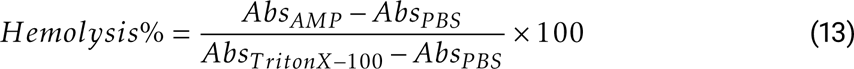

#### Cytotoxicity assay

HEK293T cells were cultured in DMEM supplemented with 10% FBS and incubated at 37°C with 5% CO_2_. After planting the cells into a 96-well plate and incubated overnight, AMP solutions were added. The gradient dilution of AMPs followed the same setting as mentioned in the hemolysis assay. Staurosporine (STS) was used as positive control and PBS as vehicle control. STS was diluted to 60, 12, 2.4, 0.48, 0.1, 0.02, 0.0038, and 0.0008 µM. After incubating for 72 hours, CellTiter-Glo® Luminescent assay was performed to evaluate the cell viability. After equilibration to room temperature, the 96-well plate containing treated cells was loaded with 100 µL CellTiter Glo (CTG, Promega) reagent each well. The plate was incubated for 30 mins, then was detected via the EnVision system at the wavelength of 570 nm. Data was fitted and analyzed with GraphPad Prism 8. All tests were in triplicate.

## Data and code availability

All datasets and codes will be available after peer review at https://github.com/ComputBiophys/AMPainter.

## Author contributions

C.S. and R.D. conceived the project. R.D. conducted the computational works and analyzed the results. Q.C. and R.D. performed the experiments. R.D. and C.S. wrote the original manuscript. All authors participated in the preparation of the manuscript. C.S. supervised the work.

## Acknowledgement

The authors thank Dr. Lei Wang, Dr. Jiaxuan Li, and Zefeng Zhu for their valuable suggestions. Part of the computation was performed on the computing platform of the Center for Life Sciences, Peking University. This work was supported by the National Key R&D Program of China (2024YFA0916800) and the Science Fund for Creative Research Groups of the National Natural Science Foundation of China (T2321001).

