## Supplementary Information for "Painting Peptides with Antimicrobial Potency through Deep Reinforcement Learning"

#### Supplementary Tables

**Table S1:** Comparison of different node embedding from pretrained language models in HyperAMP on independent test set.

| Node Embedding | Param. | Spearman | Pearson | $R^2$ | RMSE |
|---|---|---|---|---|---|
| Ankh-base | 450M | **0.8725** | **0.9227** | **0.8494** | **0.1630** |
| Ankh-large | 1.15B | 0.8568 | 0.9147 | 0.8361 | 0.1701 |
| ESM-1b | 650M | 0.8684 | 0.9174 | 0.8374 | 0.1694 |
| ESM-2 | 650M | 0.8672 | 0.9213 | 0.8482 | 0.1637 |
| ProtT5-XL-UniRef50 | 3B | 0.8666 | 0.9221 | 0.8492 | 0.1632 |

**Table S2:** Sequences and features of 30 designed peptides by AMPainter.

| Name | Sequence | Length | MW | Charge | pI | Aromaticity[a] | Boman | Hydrophobicity[b] |
|---|---|---|---|---|---|---|---|---|
| A01 | RWWRGWWRRKLKKLRKKLKKKLKGM | 25 | 3395.27 | 13.99 | 12.74 | 0.16 | 3.65 | 0.20 |
| A02 | IFHHIFRGIHHIFKGIHRLWKR | 22 | 2849.40 | 5.20 | 12.42 | 0.18 | 1.66 | 0.41 |
| A03 | WRWKGRWRRKLKKLRWKLKKKLKLG | 25 | 3377.23 | 13.99 | 12.74 | 0.16 | 3.55 | 0.20 |
| A04 | KWWKRRRWRWRWKKLWK | 17 | 2670.22 | 9.99 | 12.73 | 0.35 | 4.91 | 0.06 |
| A05 | GIKKWIKGWAKKLIKKVAKKIAKQ | 24 | 2792.55 | 9.98 | 11.63 | 0.08 | 0.85 | 0.38 |
| A06 | RWRKWRRWWWRKKKLGF | 17 | 2560.07 | 8.99 | 12.72 | 0.35 | 4.49 | 0.12 |
| A07 | GIKKWIKGAIKKLIKKVAKWIWKQ | 24 | 2892.67 | 8.99 | 11.58 | 0.13 | 0.39 | 0.38 |
| A08 | GRKKRRRRRRGGWWWKLGLRWF | 22 | 2998.55 | 10.99 | 12.95 | 0.23 | 5.01 | 0.14 |
| A09 | FFHHIFRKIHHVFKKIHRLFHH | 22 | 2962.52 | 5.28 | 12.18 | 0.23 | 1.84 | 0.45 |
| A10 | WKKWRWWWRWKKLWW | 15 | 2445.91 | 5.99 | 12.20 | 0.53 | 1.90 | 0.07 |
| S01 | KKQVKWLLKVWKKVGIKLGAKLPVWK | 26 | 3102.94 | 8.99 | 11.58 | 0.12 | 0.16 | 0.38 |
| S02 | RRLRRLRILLFLKRVLRR | 18 | 2434.08 | 8.99 | 12.95 | 0.06 | 4.64 | 0.50 |
| S03 | WVQKKVIKLILKGLFALKLLG | 21 | 2409.10 | 4.99 | 11.28 | 0.10 | -1.10 | 0.57 |
| S04 | RKRVVWGKIFFRRR | 14 | 1904.32 | 6.99 | 12.71 | 0.21 | 4.53 | 0.36 |
| S05 | GILFLKRLRILGIGKLLLLKR | 21 | 2434.16 | 5.99 | 12.44 | 0.05 | 0.07 | 0.57 |
| S06 | IKRRIRRRPIRRRQRR | 16 | 2272.76 | 10.99 | 13.06 | 0.00 | 9.10 | 0.19 |
| S07 | VKKRLLFRKPLLKLFLLFGRRLKA | 24 | 2954.78 | 8.99 | 12.61 | 0.13 | 1.35 | 0.54 |
| S08 | KQQRKRGRVRKALLGKVIGFGGLKLKG | 27 | 2992.67 | 9.99 | 12.62 | 0.04 | 2.26 | 0.33 |
| S09 | KLKIKFKFHLKLFGLF | 16 | 2007.56 | 5.03 | 11.28 | 0.25 | -0.32 | 0.56 |
| S10 | GYFKRVVLRIIVKVKKVKIII | 21 | 2513.25 | 6.99 | 11.64 | 0.10 | 0.19 | 0.57 |
| R01 | RLKIHLYIKHYRRPKIVIRLKIK | 23 | 2985.76 | 9.07 | 11.82 | 0.09 | 2.33 | 0.39 |
| R02 | KWFFKWFRKHFLKWMRQYFHK | 21 | 3075.69 | 7.07 | 11.64 | 0.43 | 2.07 | 0.33 |
| R03 | YWRRWYIWRWRRWRWLRYI | 19 | 3057.57 | 6.98 | 11.99 | 0.47 | 4.01 | 0.16 |
| R04 | RGWFKVKRRYIKRFMRGLRGFHIKAA | 26 | 3278.98 | 10.03 | 12.43 | 0.19 | 2.99 | 0.38 |
| R05 | WWWHLRIWLKKKHQKAQWKKFGWFMKQHH | 29 | 4014.77 | 8.15 | 11.90 | 0.28 | 1.70 | 0.24 |
| R06 | LWRWKLLGRISVIIGKIKAALRLWF | 25 | 3037.78 | 5.99 | 12.44 | 0.16 | 0.04 | 0.52 |
| R07 | WIPKRFGIVIKRFYYRPFYPLFQ | 23 | 3035.64 | 4.98 | 10.90 | 0.35 | 1.00 | 0.39 |
| R08 | YKGFWRKIIYLIMMHAMKFMVWFLHLPKQ | 29 | 3757.70 | 5.07 | 10.81 | 0.24 | -0.24 | 0.52 |
| R09 | KLYKYKYRKVYKKQRRR | 17 | 2404.91 | 9.98 | 11.26 | 0.24 | 5.30 | 0.12 |
| R10 | QPKKPYKPIYKKVRGVLIPFKRV | 23 | 2783.46 | 7.98 | 11.30 | 0.13 | 1.66 | 0.30 |

[a] Relative frequency of aromatic residues (F, W, Y).
[b] Relative frequency of hydrophobic residues (A, C, F, I, L, M, V).

**Table S3:** E-values and most similar fragments of 30 designed peptides searched in known AMPs by BLAST.

| Name | Length | Most similar fragment | E-value |
|---|---|---|---|
| A01 | 25 | WSGMWRRKLKKLRNALKKKLKG | 9.00E-05 |
| A02 | 22 | FHHIFRGIVHVGKTIHRL | 0.002 |
| A03 | 25 | SMWSGMWRRKLKKLRNALKKKLK | 3.00E-04 |
| A04 | 17 | EWFKCRRWQWRMKKL | 0.5 |
| A05 | 24 | GIKDWIKGAAKTLIKTVASHIANQ | 7.00E-05 |
| A06 | 17 | WFKCRRWQWRMKKLG | 0.67 |
| A07 | 24 | GIKDWIKGAAKTLIKTVASHIANQ | 2.00E-04 |
| A08 | 22 | / | / |
| A09 | 22 | FFHHIFRGIVHVGKTIHRL | 5.00E-04 |
| A10 | 15 | / | / |
| S01 | 26 | / | / |
| S02 | 18 | / | / |
| S03 | 21 | KGLFAL | 2 |
| S04 | 14 | / | / |
| S05 | 21 | LFKKILKYLGIGKFL | 5 |
| S06 | 16 | IRRRP | 7.5 |
| S07 | 24 | PL−FLLF | 4.3 |
| S08 | 27 | / | / |
| S09 | 16 | HLKLF | 3.3 |
| S10 | 21 | / | / |
| R01 | 23 | RRPKI | 9 |
| R02 | 21 | KSFFKSFRK | 1.8 |
| R03 | 19 | RRWY | 9 |
| R04 | 26 | RFMKG-KGFHI | 0.068 |
| R05 | 29 | / | / |
| R06 | 25 | KLLGRI | 1.3 |
| R07 | 23 | WIADQFGI | 7.9 |
| R08 | 29 | / | / |
| R09 | 17 | / | / |
| R10 | 23 | YKVP-YKKVESRAVL | 2.9 |

**Table S4:** Minimal inhibitory concentrations (MICs) of the initial sequences of A01-A10 against four bacteria.

| Initial AMP | Evolved AMP | MIC (μM) | | | | Hemolysis and cytotoxicity | Source |
|---|---|---|---|---|---|---|---|
| | | *E.coli* | *P.aeruginosa* | *S.aureus* | *B.subtilis* | | |
| Latarcin-1 | A01, A03 | 4 | 8 | 4 | 2 | 80 μM for 20% hemolysis on rabbit erythrocytes | DBAASP |
| Piscidin 1 [F1A] | A02 | 16 | >128 | 4 | 4 | 100 μM for 100% hemolysis on human erythrocytes | DBAASP |
| LAP5 | A04, A06 | >128 | >128 | 128 | 128 | 400 μM for 1% hemolysis on sheep erythrocytes | DBAASP |
| Ascaphin-5 | A05, A07 | 16 | 128 | 4 | 16 | $HC_{50}$ > 200 μM on human erythrocytes | DRAMP |
| TAT-Ras-GAP317-326 | A08 | 128 | >128 | 8 | 32 | Enhances genotoxin-induced cytotoxicity in tumor cells | DRACP |
| Piscidin 1 [G8A] | A09 | 16 | >128 | 4 | 4 | $HC_{50}$ = 4 μM on human erythrocytes | DBAASP |

**Table S5:** Hemolysis and cytotoxicity values and selectivity index (SI) of designed AMPs.

| **Name** | **Mean MIC** | **$HC_{25}$** | **$CC_{50}$** | **SI[a]** | **SI[b]** |
|---|---|---|---|---|---|
| A01 | 3.5 | 14.66 | 11.62 | 4.19 | 3.320 |
| A02 | >24 | 30.2 | 11.97 | / | / |
| A03 | 5.5 | >128 | 21.67 | >23.27 | 3.940 |
| A04 | >3 | >128 | 49.63 | / | / |
| A05 | 1.44 | 18.41 | 5.41 | 12.78 | 3.757 |
| A06 | 22.5 | >128 | 71.09 | >5.69 | 3.160 |
| A07 | 6.63 | 7.73 | 3 | 1.17 | 0.452 |
| A08 | >64 | 17.94 | 12.78 | / | / |
| A09 | 51 | 8.5 | 7.28 | 0.17 | 0.143 |
| A10 | >25.33 | 30.13 | 51.11 | / | / |
| S01 | 21 | >128 | 39.14 | >6.10 | 1.864 |
| S02 | 49.5 | >128 | >128 | >2.59 | >2.586 |
| S03 | >128 | >128 | >64 | / | / |
| S04 | >64.5 | >128 | >128 | / | / |
| S05 | >82 | >128 | >128 | / | / |
| S06 | >128 | >128 | >128 | / | / |
| S07 | 10.75 | >128 | 22.12 | >11.91 | 2.058 |
| S08 | >82 | >128 | >128 | / | / |
| R01 | >128 | 11.07 | >16 | / | / |
| R02 | 51 | >128 | 7.33 | >2.51 | 0.144 |
| R03 | >128 | >128 | >128 | / | / |
| R04 | 2.88 | >128 | 23.59 | >44.44 | 8.190 |
| R05 | >66.5 | >128 | 13.97 | / | / |
| R09 | >112 | >128 | >128 | / | / |

[a] SI = $HC_{25}$ / Mean MIC
[b] SI = $CC_{50}$ / Mean MIC

**Table S6:** Lipid bilayer compositions of the two molecular dynamics simulation systems.

| **Membrane** | **Lipid** | **Number** |
|---|---|---|
| *E.coli* inner membrane | PVPE | 75 |
| | PVPG | 20 |
| | PVCL2 | 5 |
| Human plasma membrane | Cholesterol | 36 |
| | PLPC | 16 |
| | SOPC | 8 |
| | PAPC | 6 |
| | SAPS | 1 |
| | PSM | 13 |
| | LSM | 8 |
| | NSM | 10 |
| | PLA20(PE) | 2 |

### Supplementary Figures

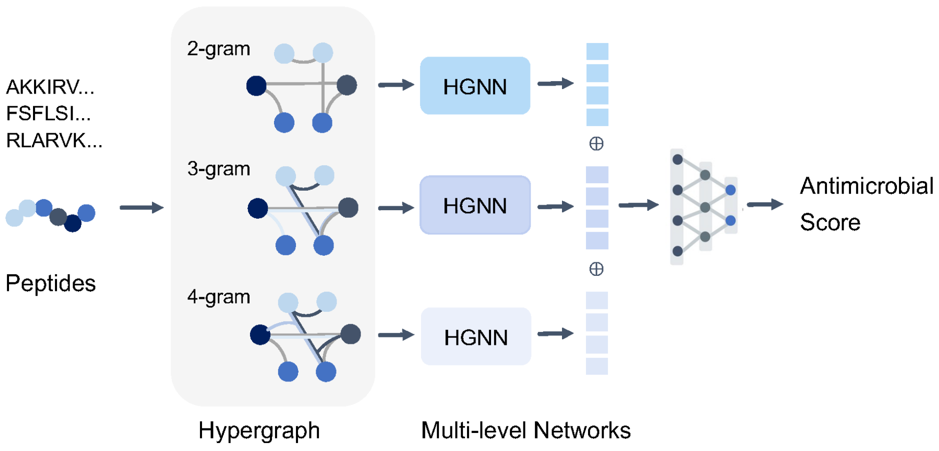


**Figure S1:** Framework of the multilevel hypergraph network predictor HyperAMP.

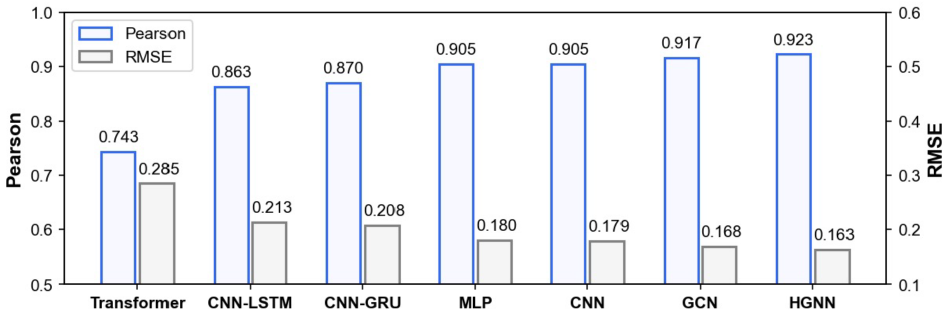


**Figure S2:** Comparison of baseline networks on antimicrobial regression with hypergraph neural network.

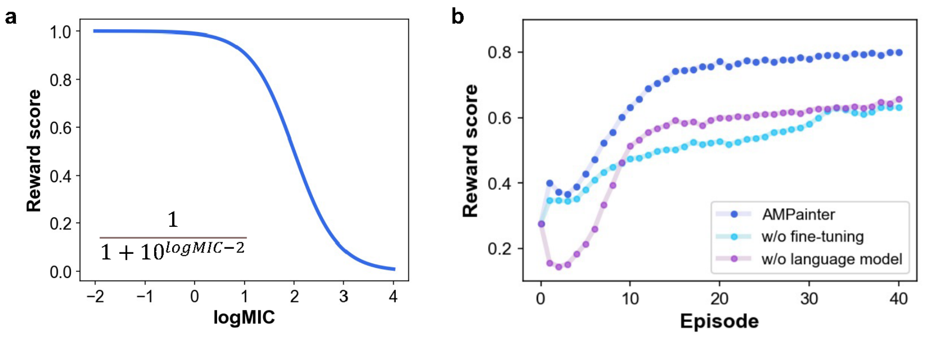


**Figure S3:** Training AMPainter. **a**, The reward function. **b**, Results of ablation study.

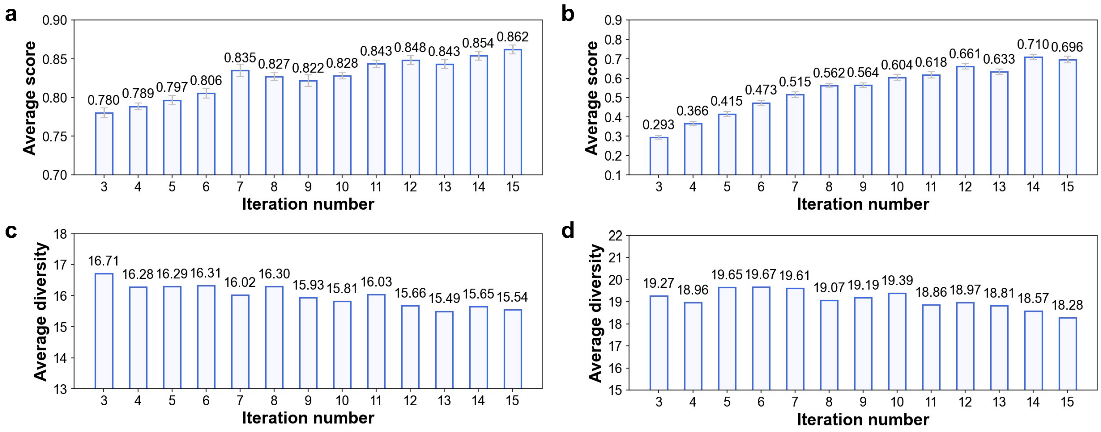


**Figure S4:** Comparison of setting different iteration numbers for AMPainter. **a**, Average scores for evolving known AMPs. **b**, Average scores for evolving random sequences. **c**, Diversity for evolving known AMPs. **d**, Diversity for evolving random sequences. Error bars show the standard error of 10 steps.

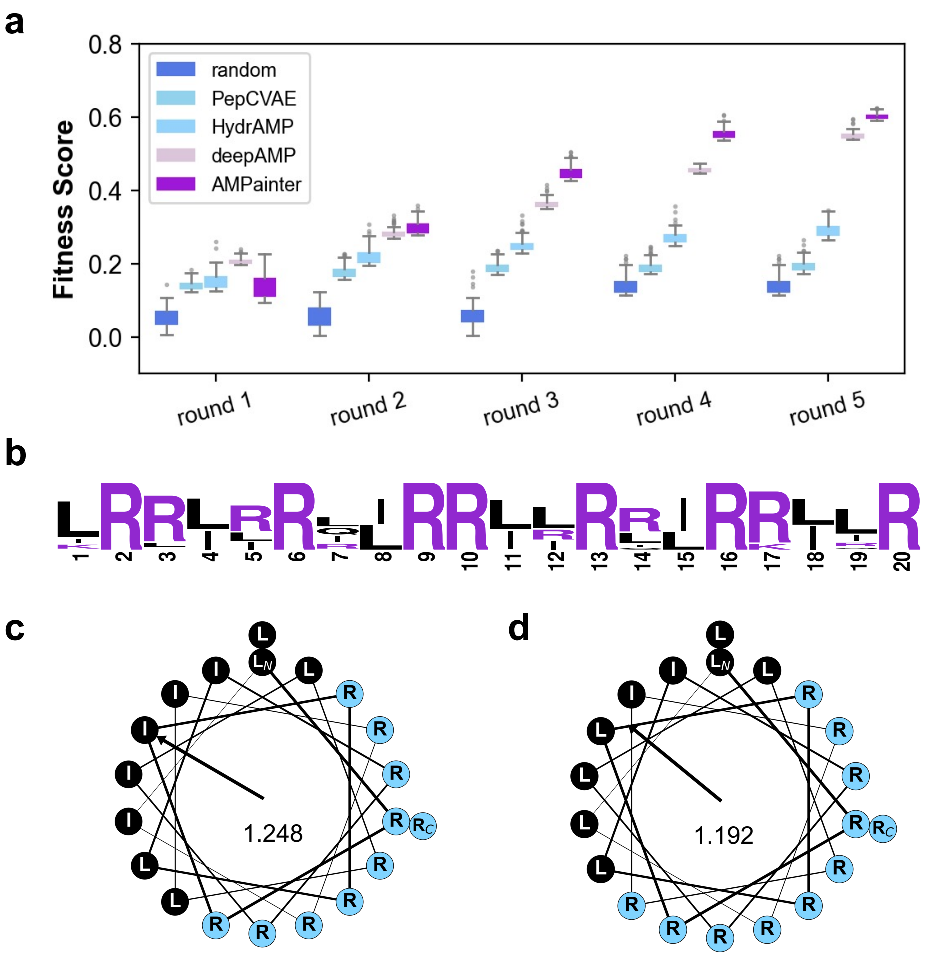


**Figure S5:** Evolution of Pg-AMP1 fragments guided by fitness score. **a**, Top 100 sequences of five rounds of optimization. **b**, Sequence logo of top 100 sequences evolved by AMPainter at round 5 (created with weblogo.berkeley.edu). **c**, Helical wheel of the top 1 sequence by AMPainter with fitness score 0.624. **d**, Helical wheel of the converged sequence of top 100 sequences by AMPainter (obtained from the sequence logo in **b**). Hydrophobic moments are shown in **c** and **d**.

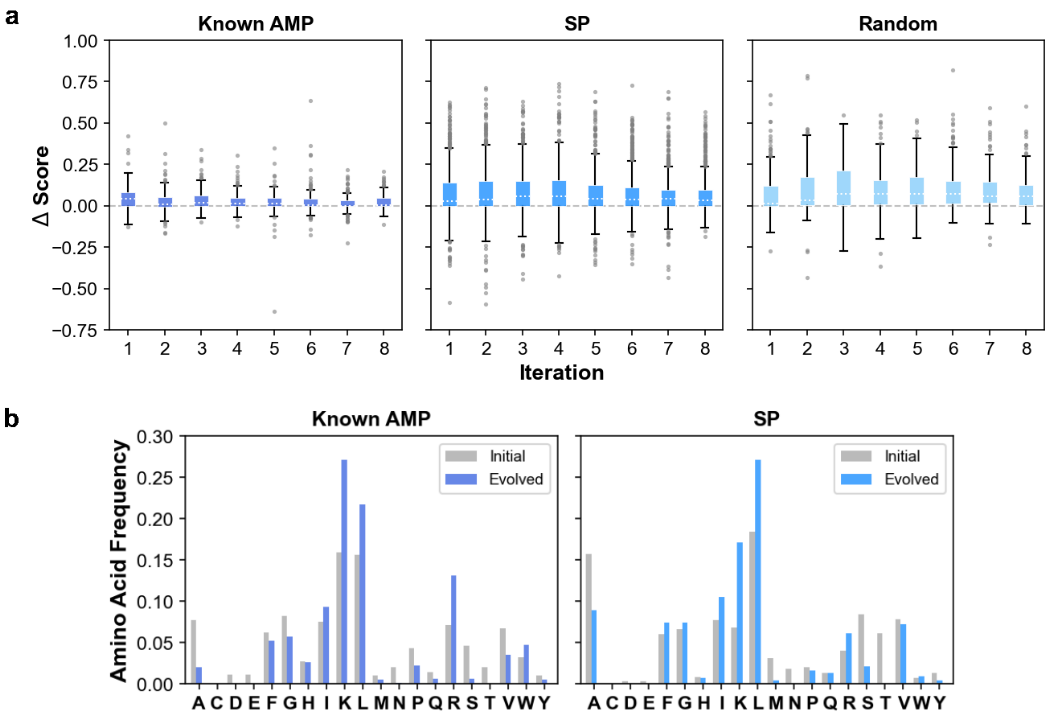


**Figure S6:** Evolving results of three sets of initial sequences. **a**, Increasing scores of each peptide along with iterations. **b**, Amino acid frequency of known AMP and random sequence before and after evolving.

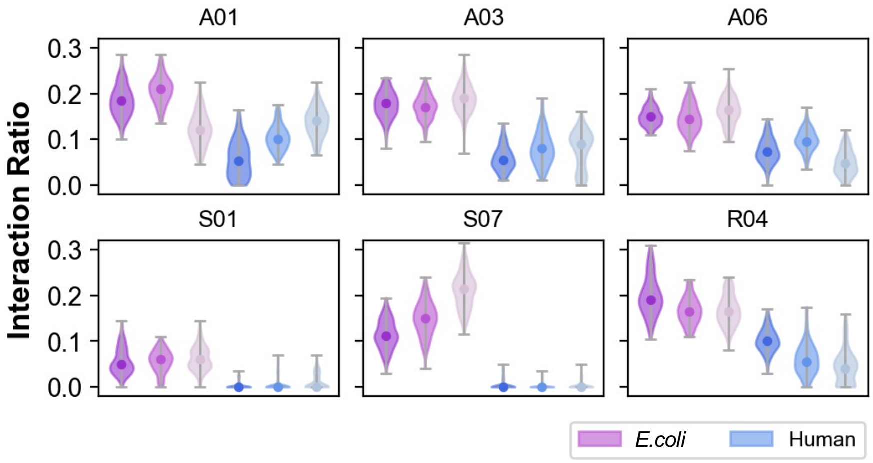


**Figure S7:** Interaction frame ratios of AMP heavy atoms with membrane heavy atoms in MD simulations. For each AMP heavy atom, this ratio is calculated as the number of interacted frames divided by the number of extracted frames. A frame is defined as *interacted* if the distance between this atom and any membrane heavy atom is greater than 3.5 Å.

**a**

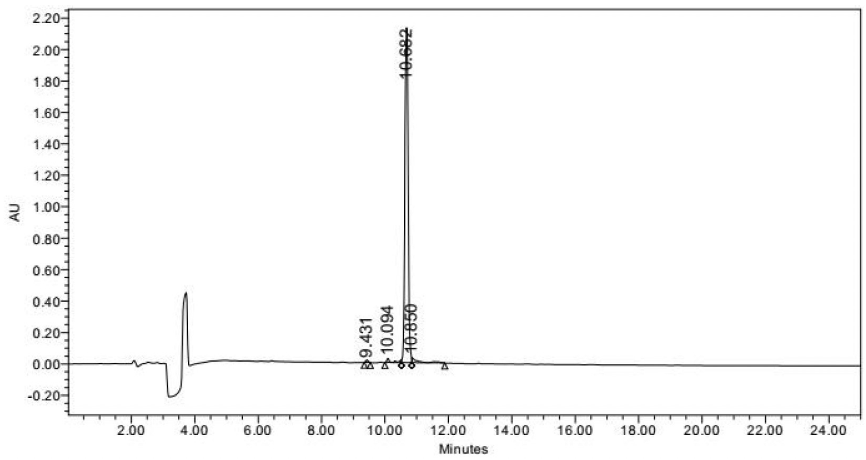


**b**

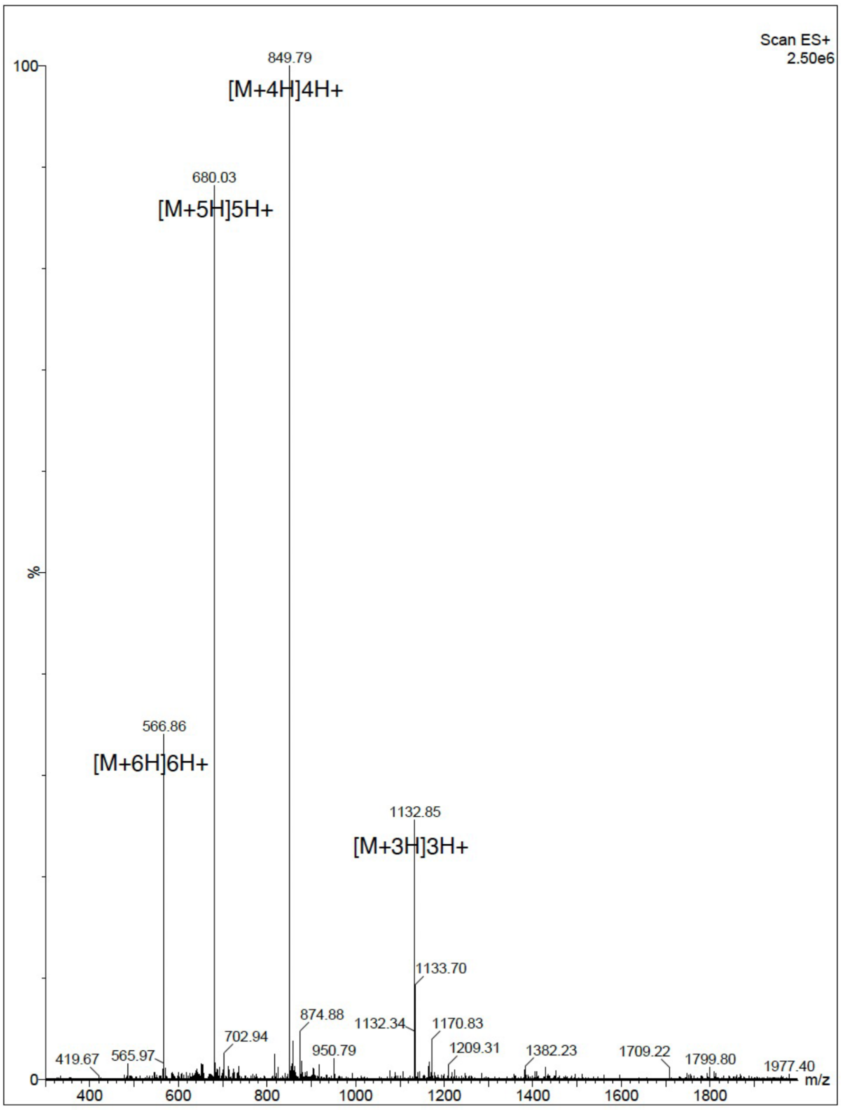


**Figure S8:** Validation of synthesized peptide A01. **a**, HPLC chromatography. **b**, Mass spectrometry.

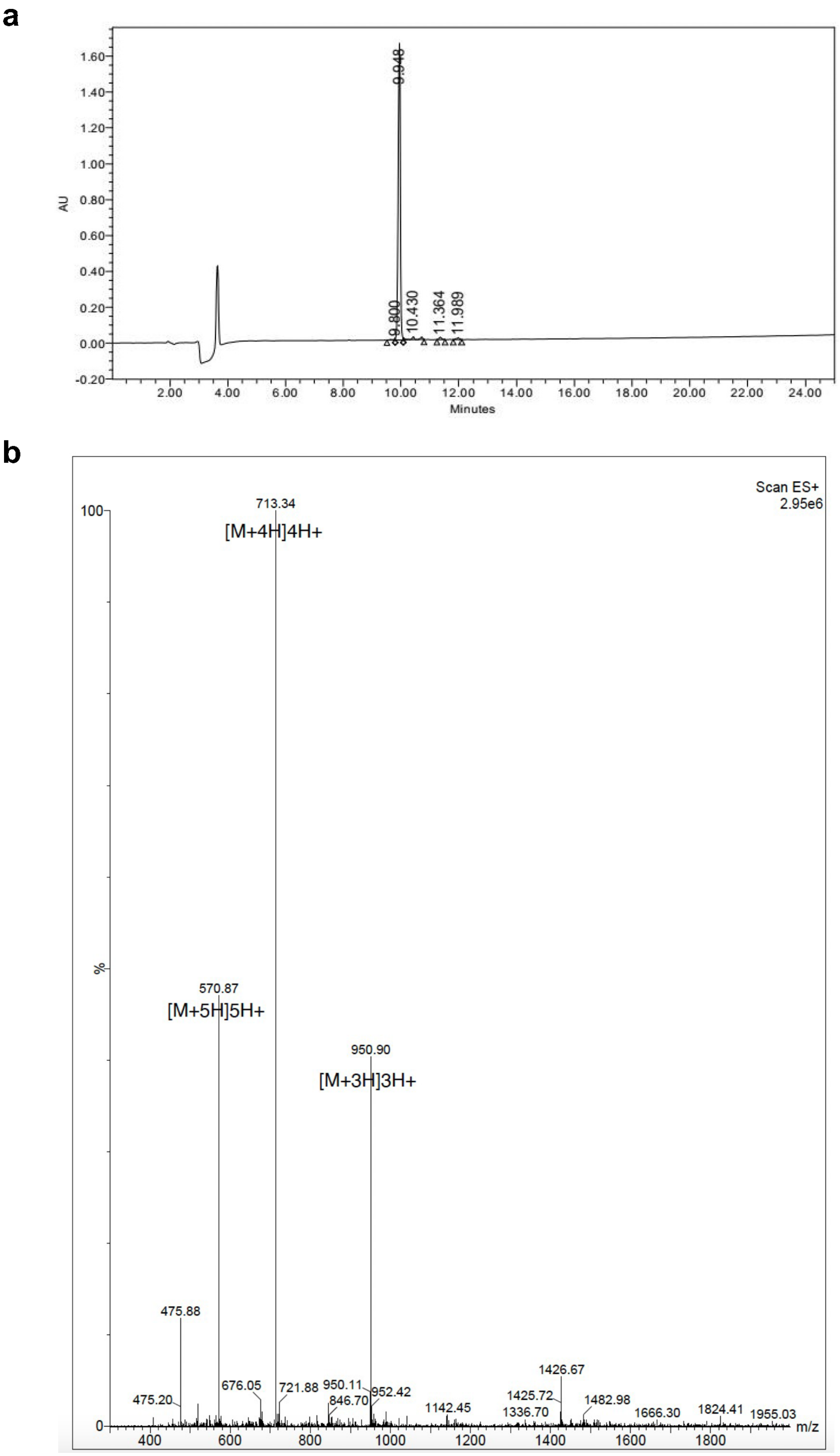


**Figure S9:** Validation of synthesized peptide A02. **a**, HPLC chromatography. **b**, Mass spectrometry.

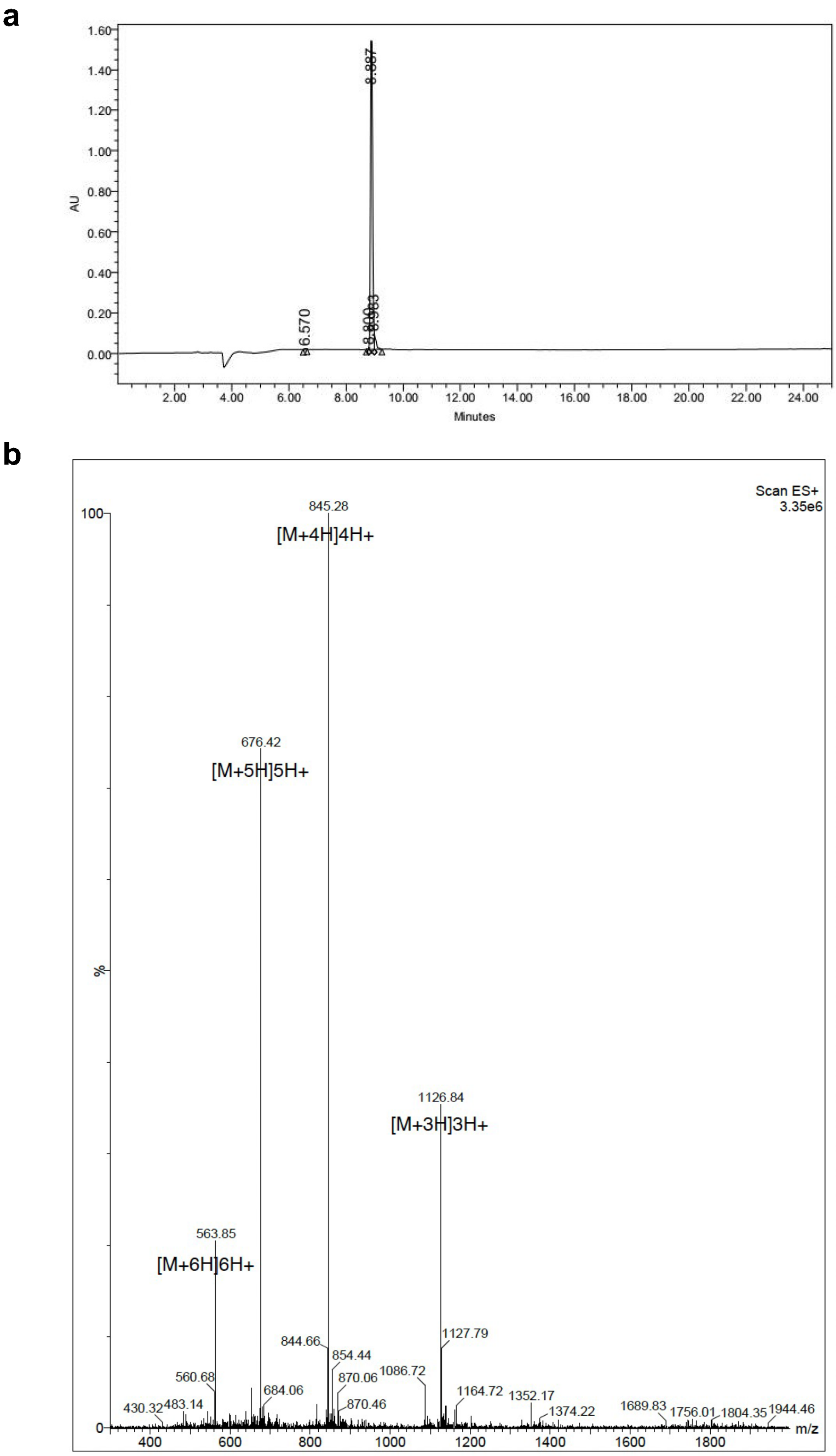


**Figure S10:** Validation of synthesized peptide A03. **a**, HPLC chromatography. **b**, Mass spectrometry.

**a**

**b**

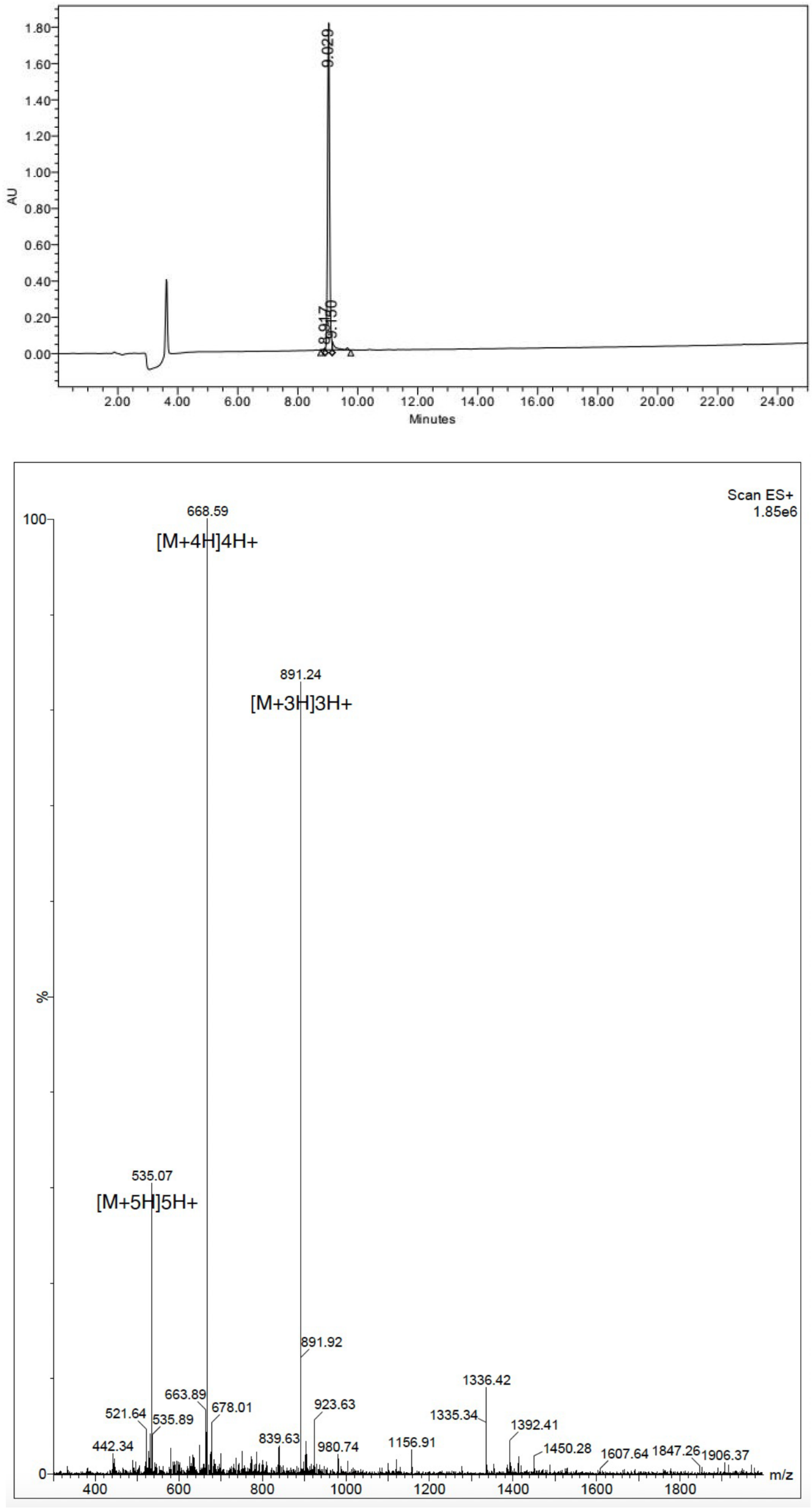


**Figure S11:** Validation of synthesized peptide A04. **a**, HPLC chromatography. **b**, Mass spectrometry.

**a**

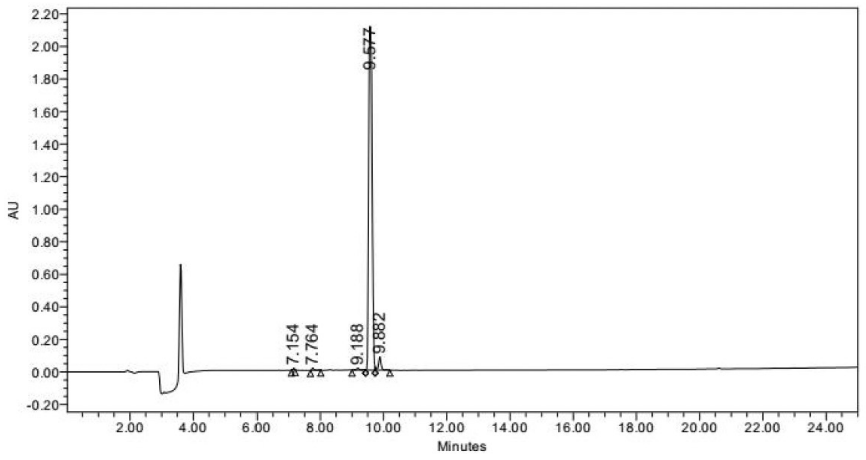


**b**

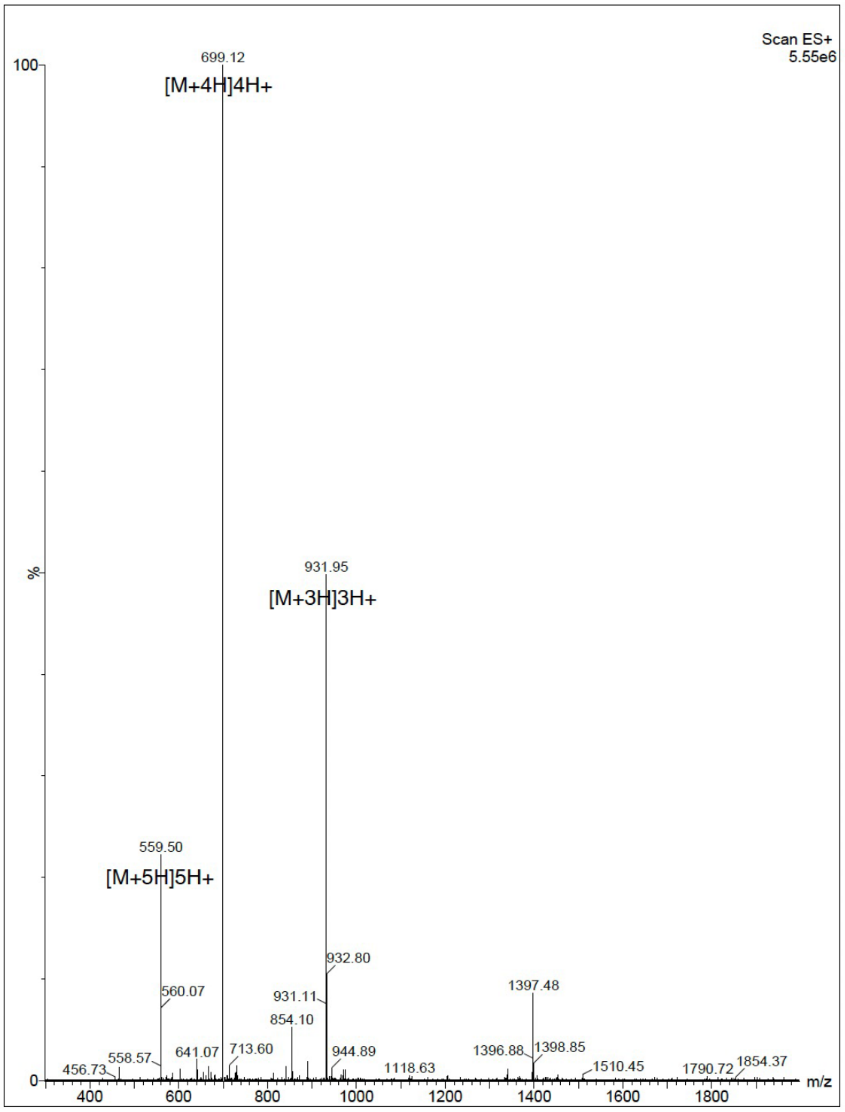


**Figure S12:** Validation of synthesized peptide A05. **a**, HPLC chromatography. **b**, Mass spectrometry.

a

b

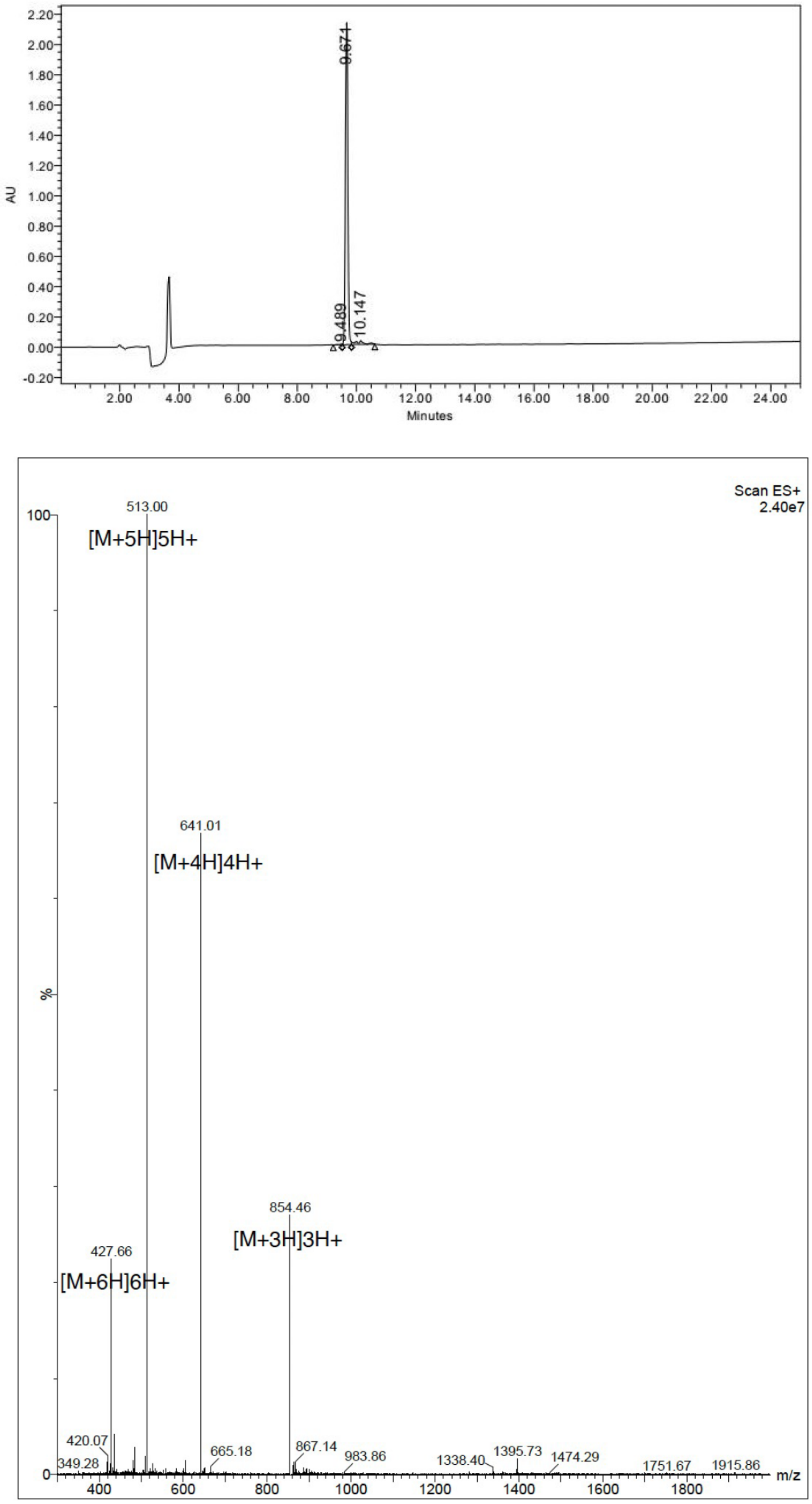


**Figure S13:** Validation of synthesized peptide A06. **a**, HPLC chromatography. **b**, Mass spectrometry.

**a**

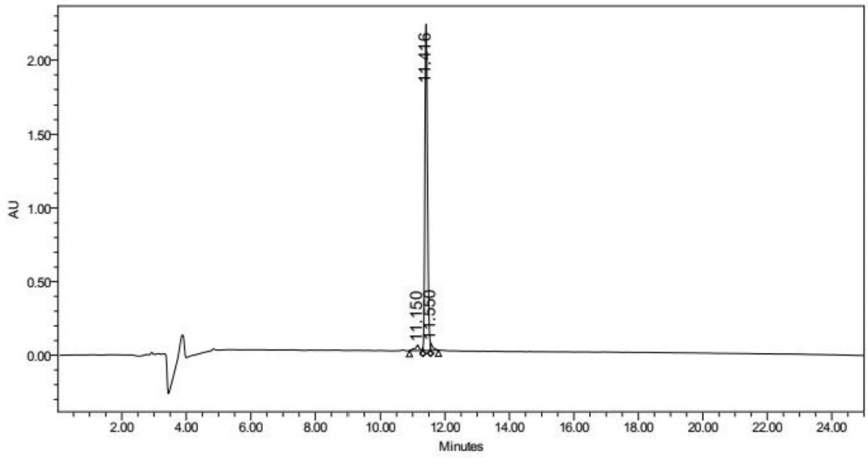


**b**

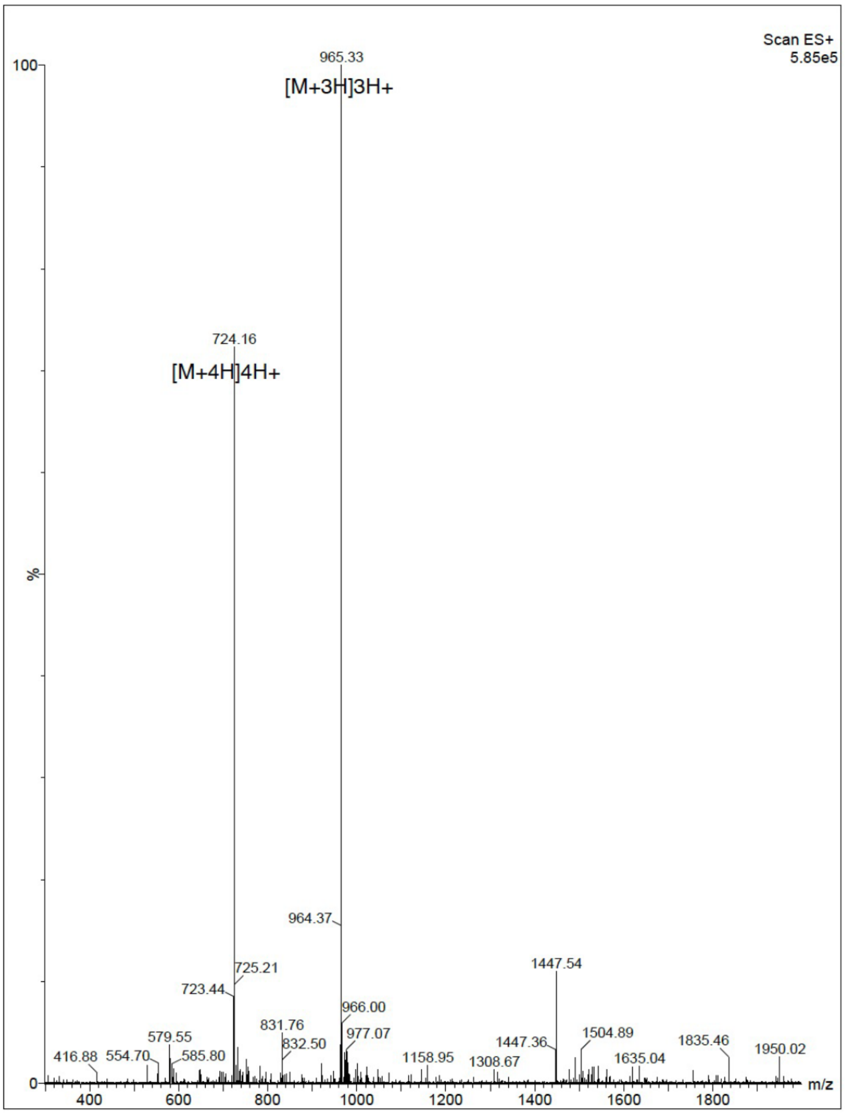


**Figure S14:** Validation of synthesized peptide A07. **a**, HPLC chromatography. **b**, Mass spectrometry.

a

b

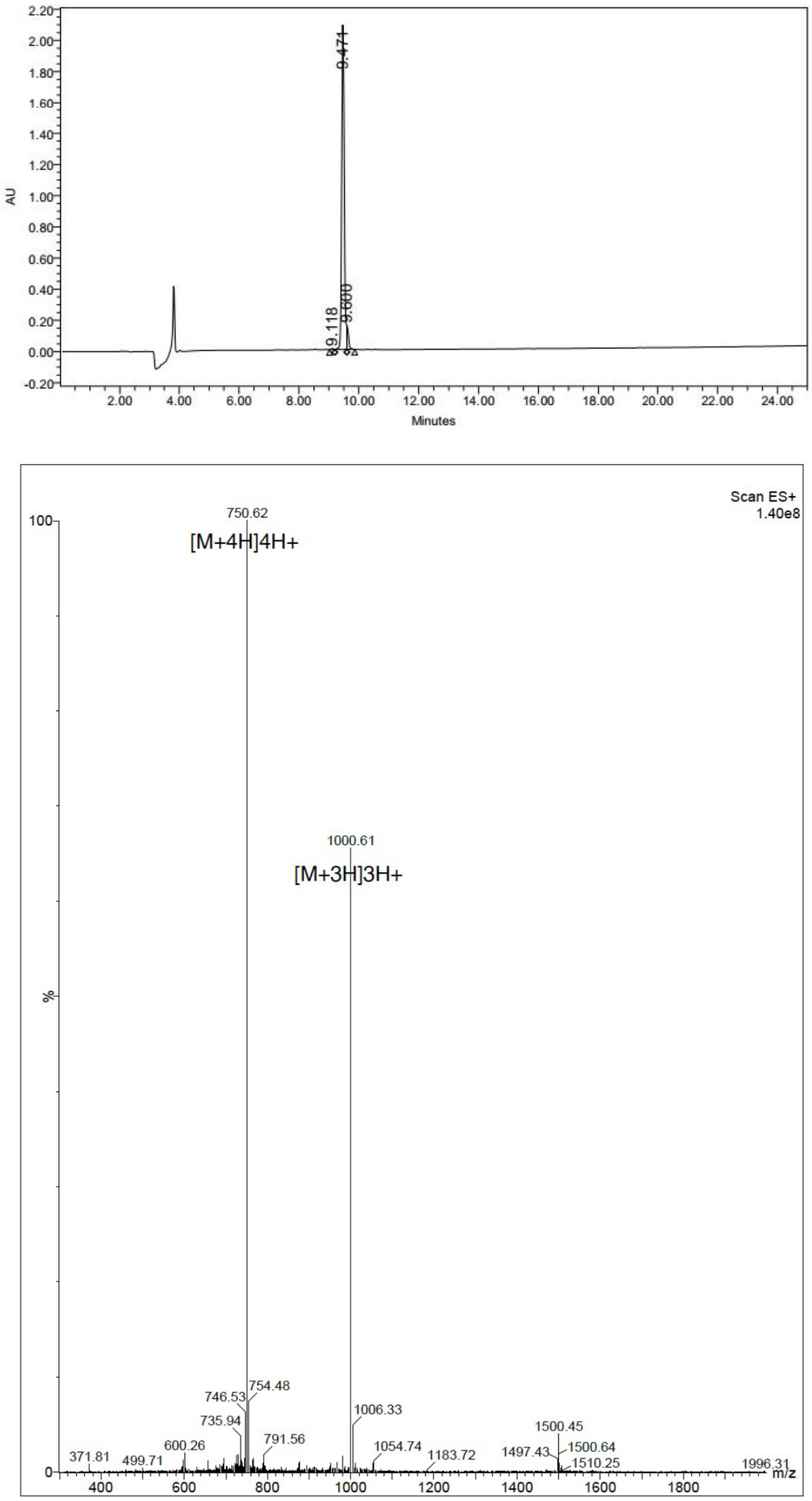


**Figure S15:** Validation of synthesized peptide A08. **a**, HPLC chromatography. **b**, Mass spectrometry.

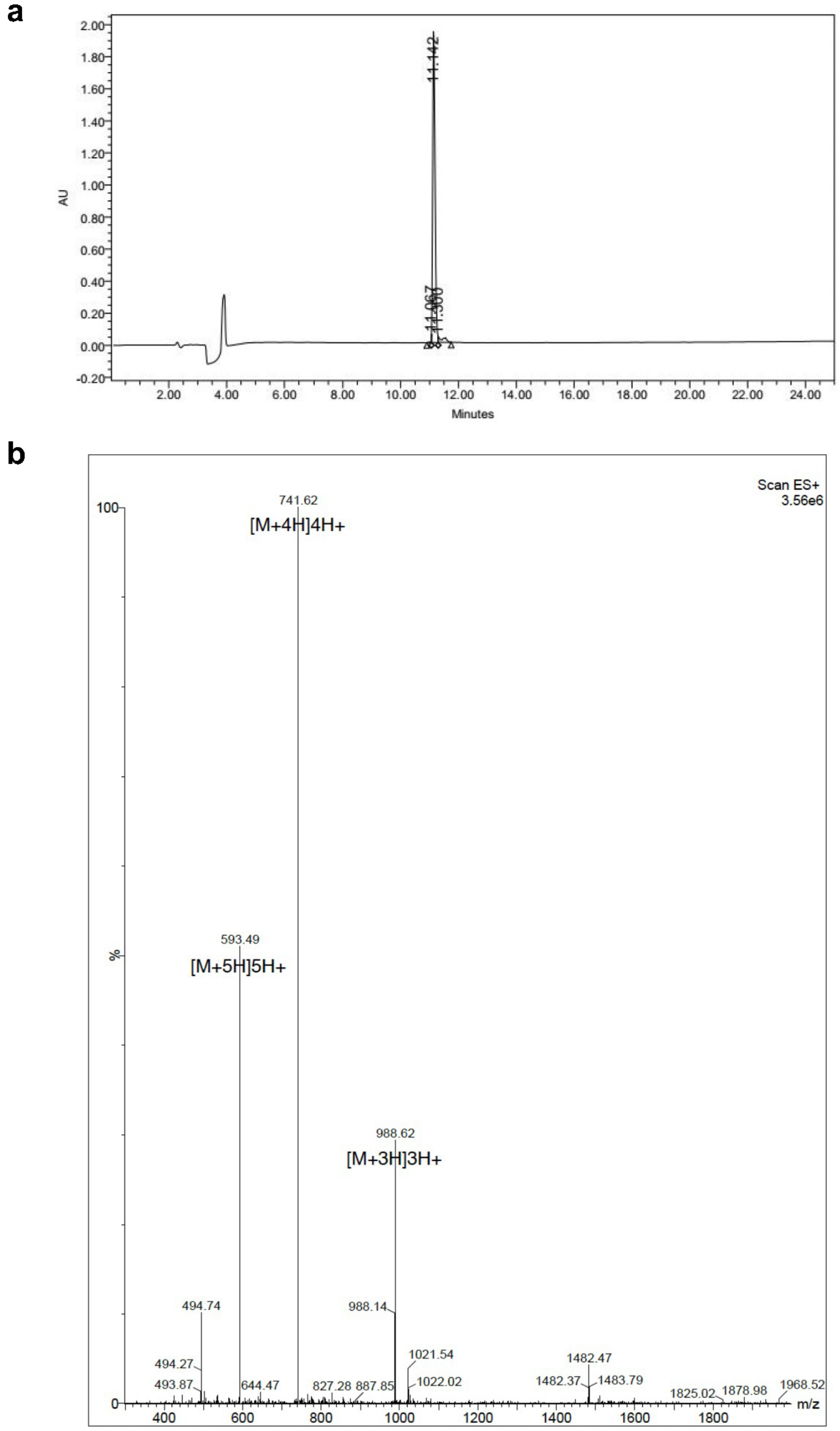


**Figure S16:** Validation of synthesized peptide A09. **a**, HPLC chromatography. **b**, Mass spectrometry.

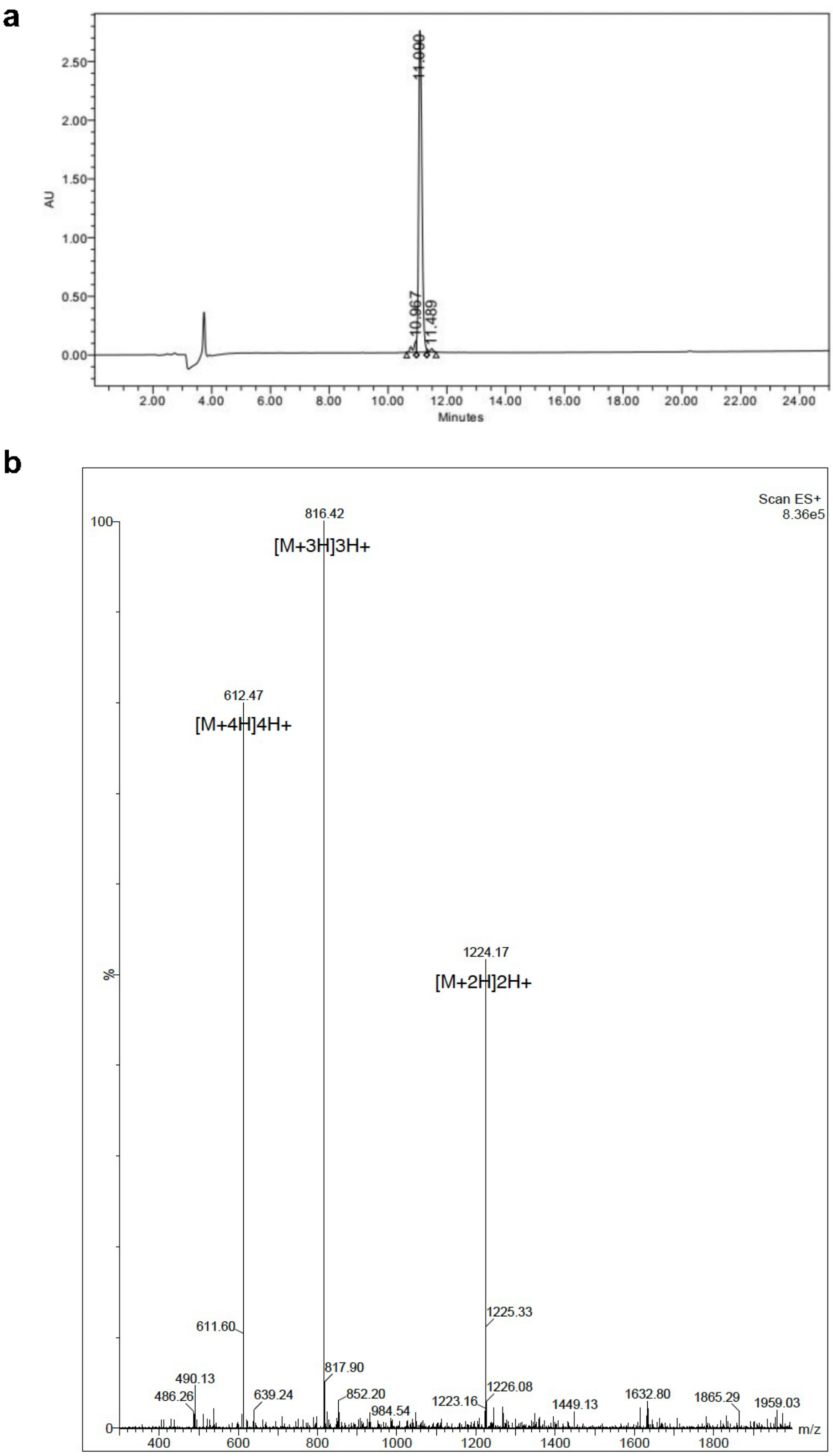


**Figure S17:** Validation of synthesized peptide A10. **a**, HPLC chromatography. **b**, Mass spectrometry.

**a**

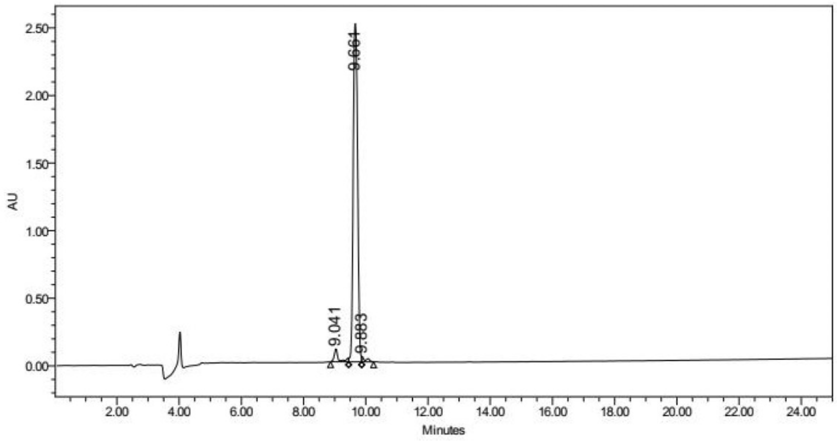


**b**

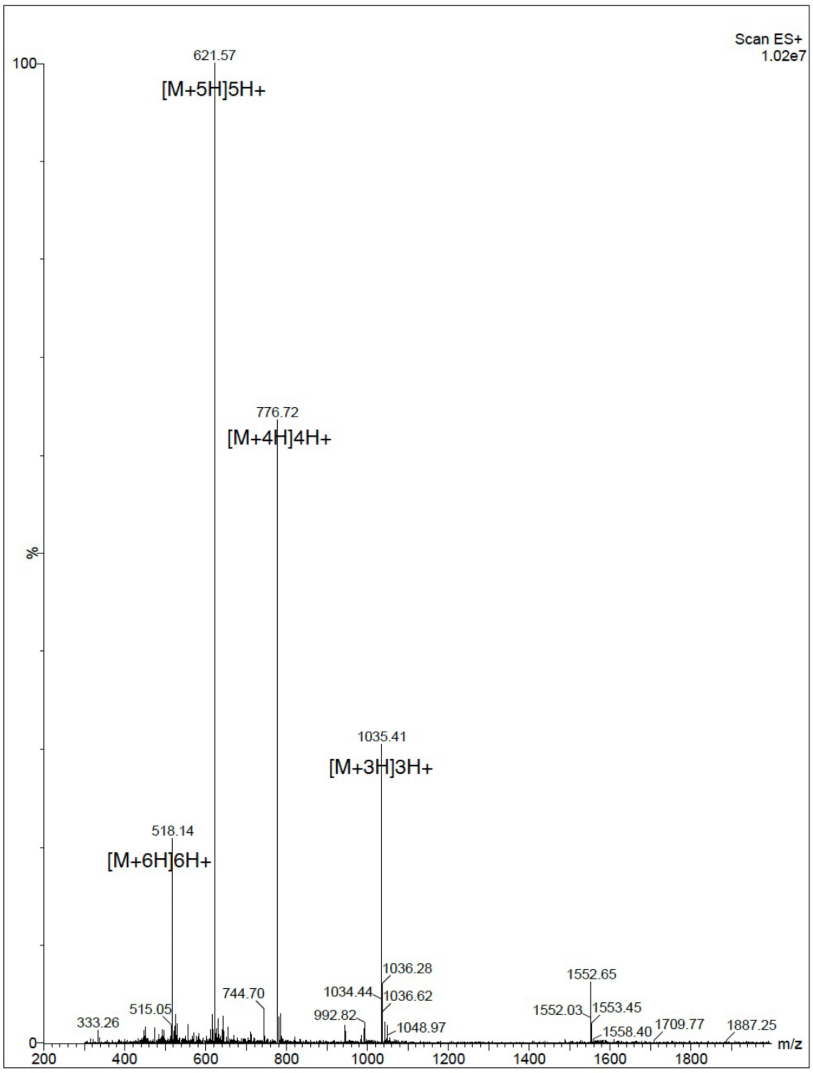


**Figure S18:** Validation of synthesized peptide S01. **a**, HPLC chromatography. **b**, Mass spectrometry.

a
b

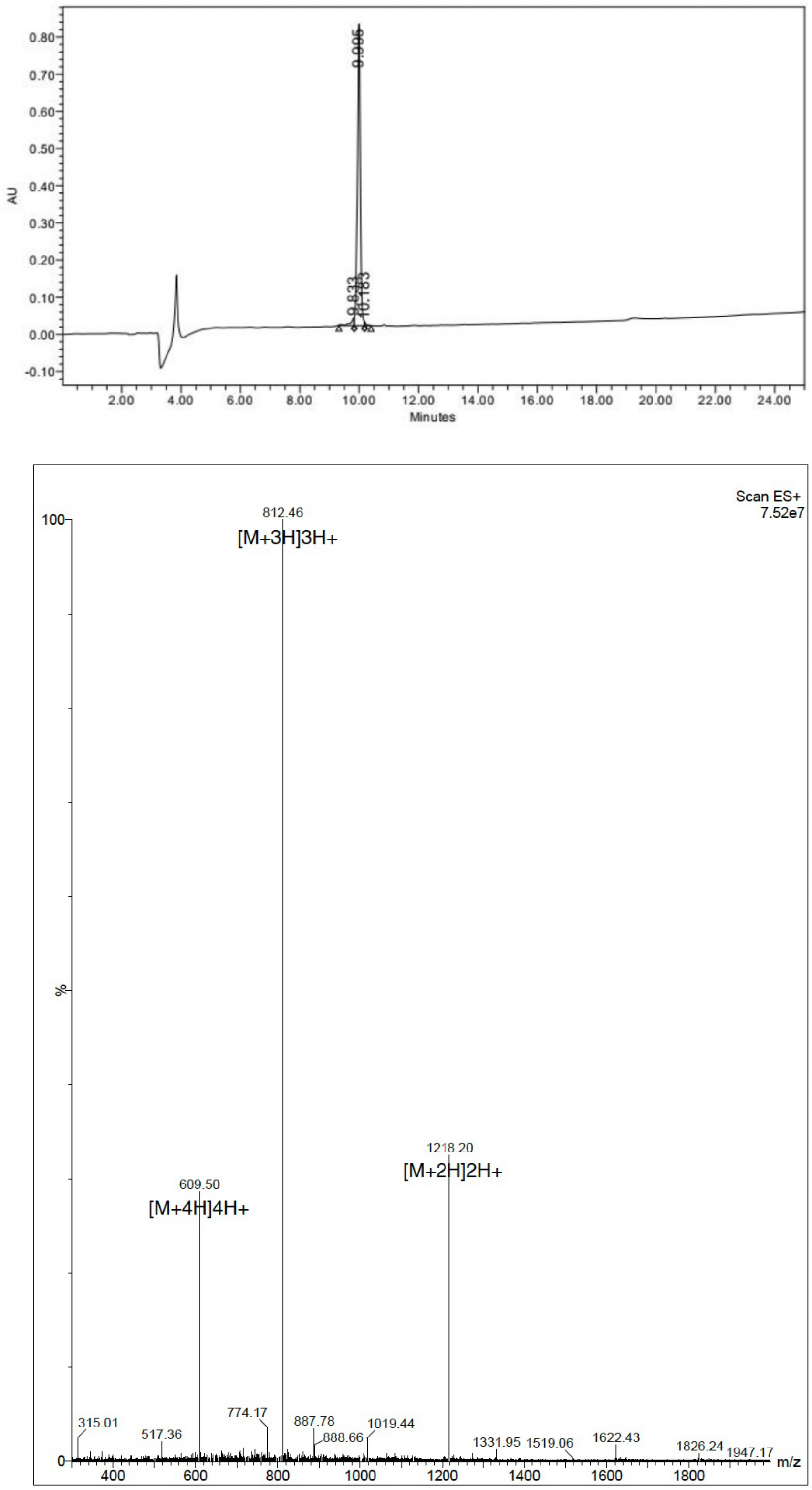


**Figure S19:** Validation of synthesized peptide S02. **a**, HPLC chromatography. **b**, Mass spectrometry.

a

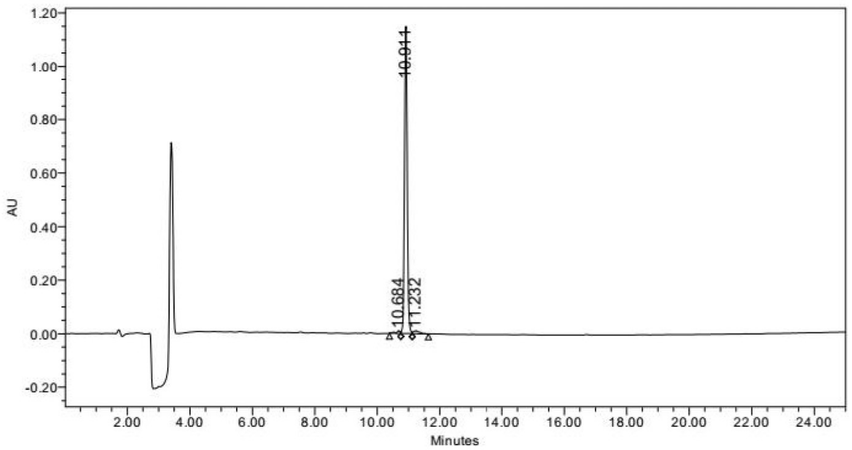


b

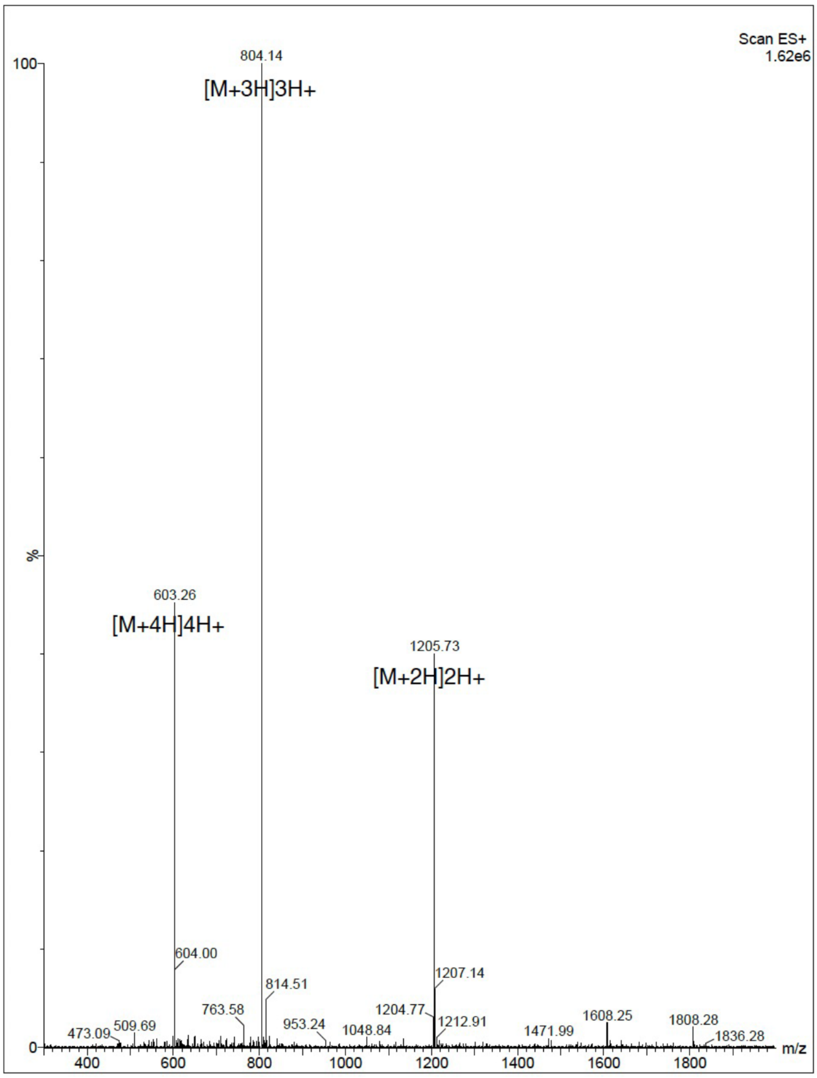


**Figure S20:** Validation of synthesized peptide S03. **a**, HPLC chromatography. **b**, Mass spectrometry.

**a**

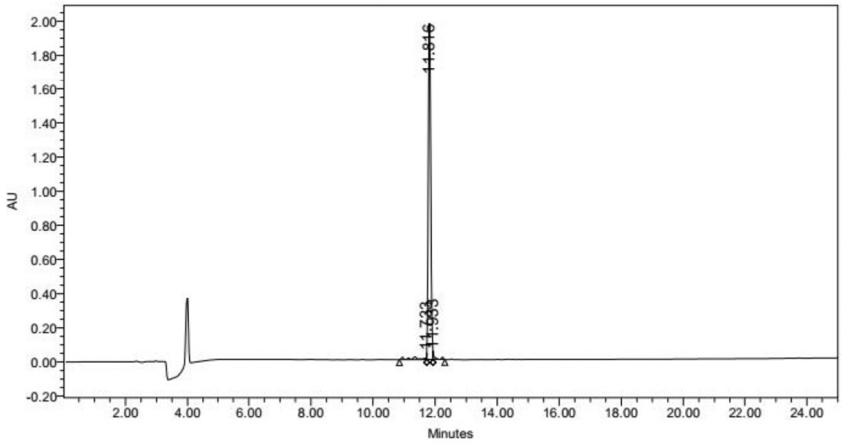


**b**

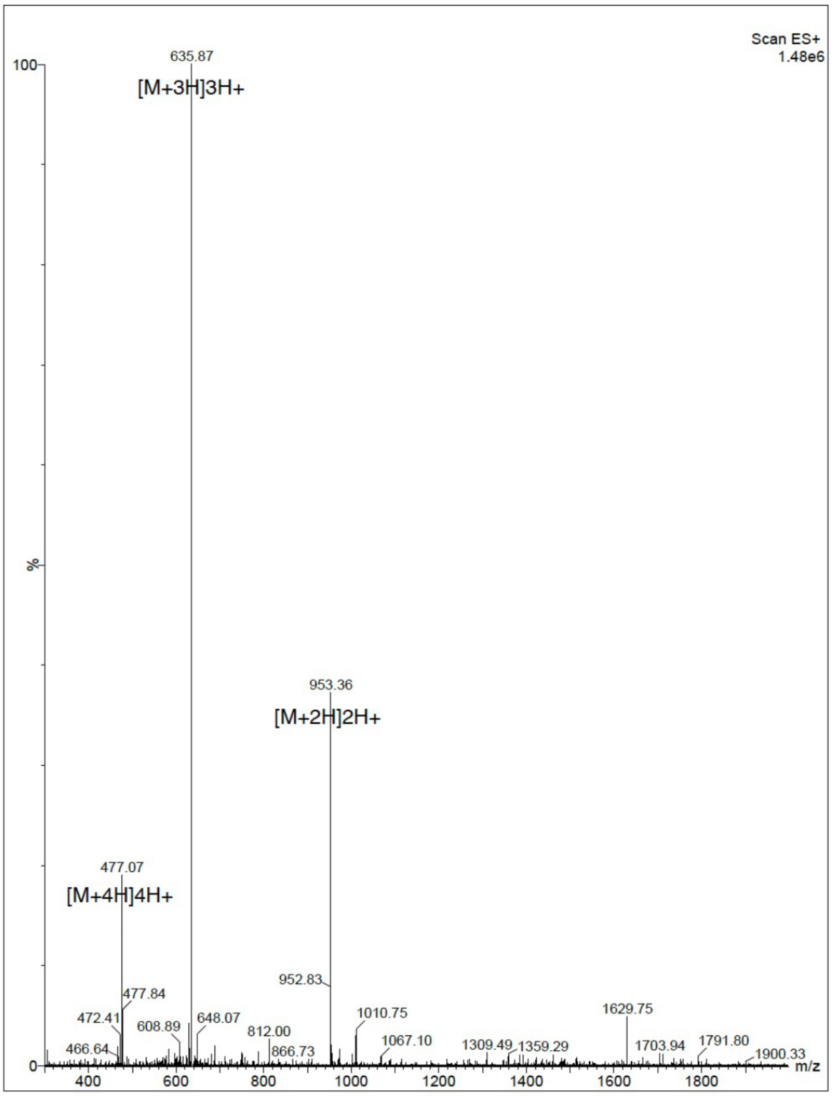


**Figure S21:** Validation of synthesized peptide S04. **a**, HPLC chromatography. **b**, Mass spectrometry.

a

b

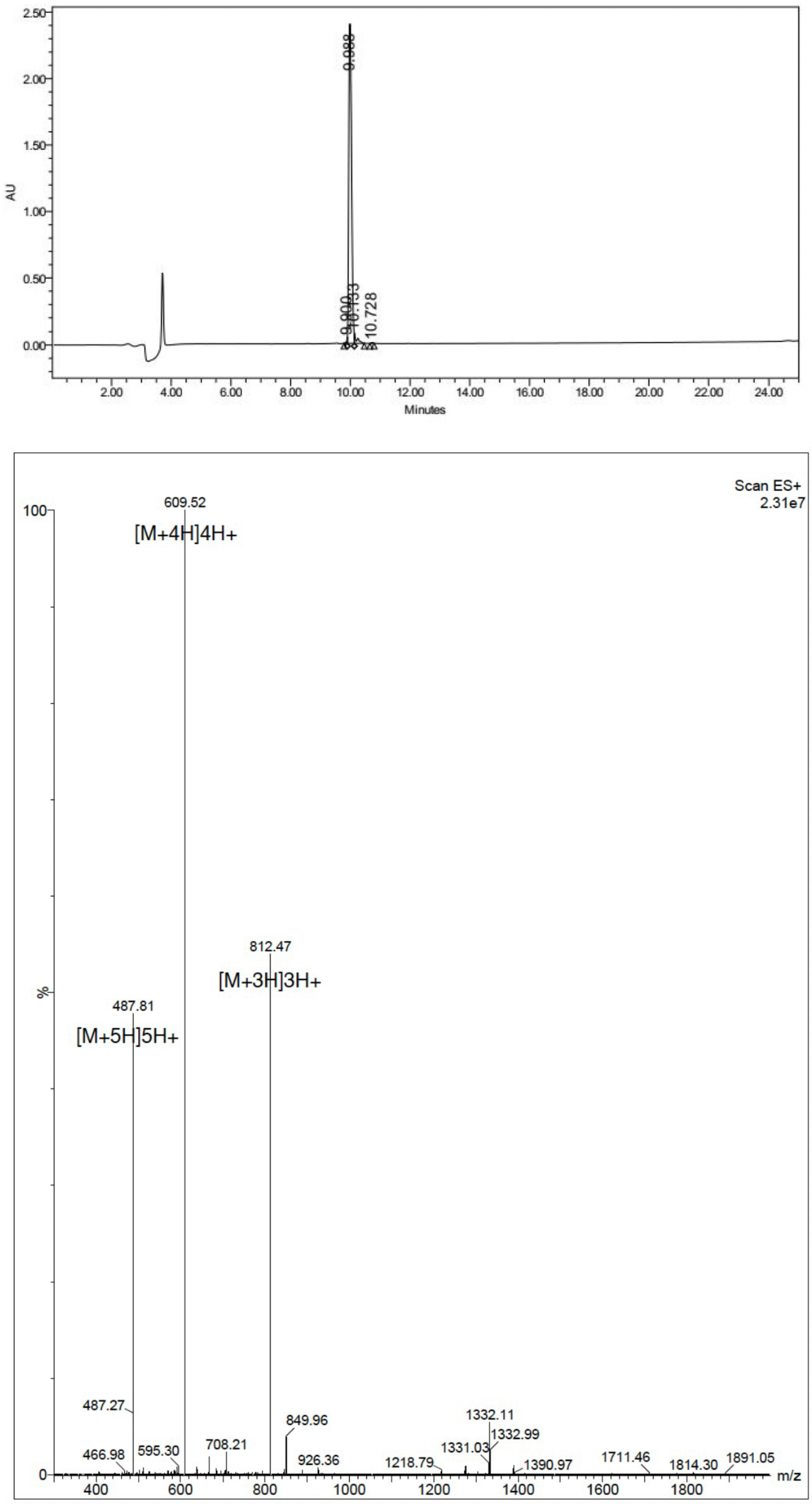


**Figure S22:** Validation of synthesized peptide S05. **a**, HPLC chromatography. **b**, Mass spectrometry.

**a**

**b**

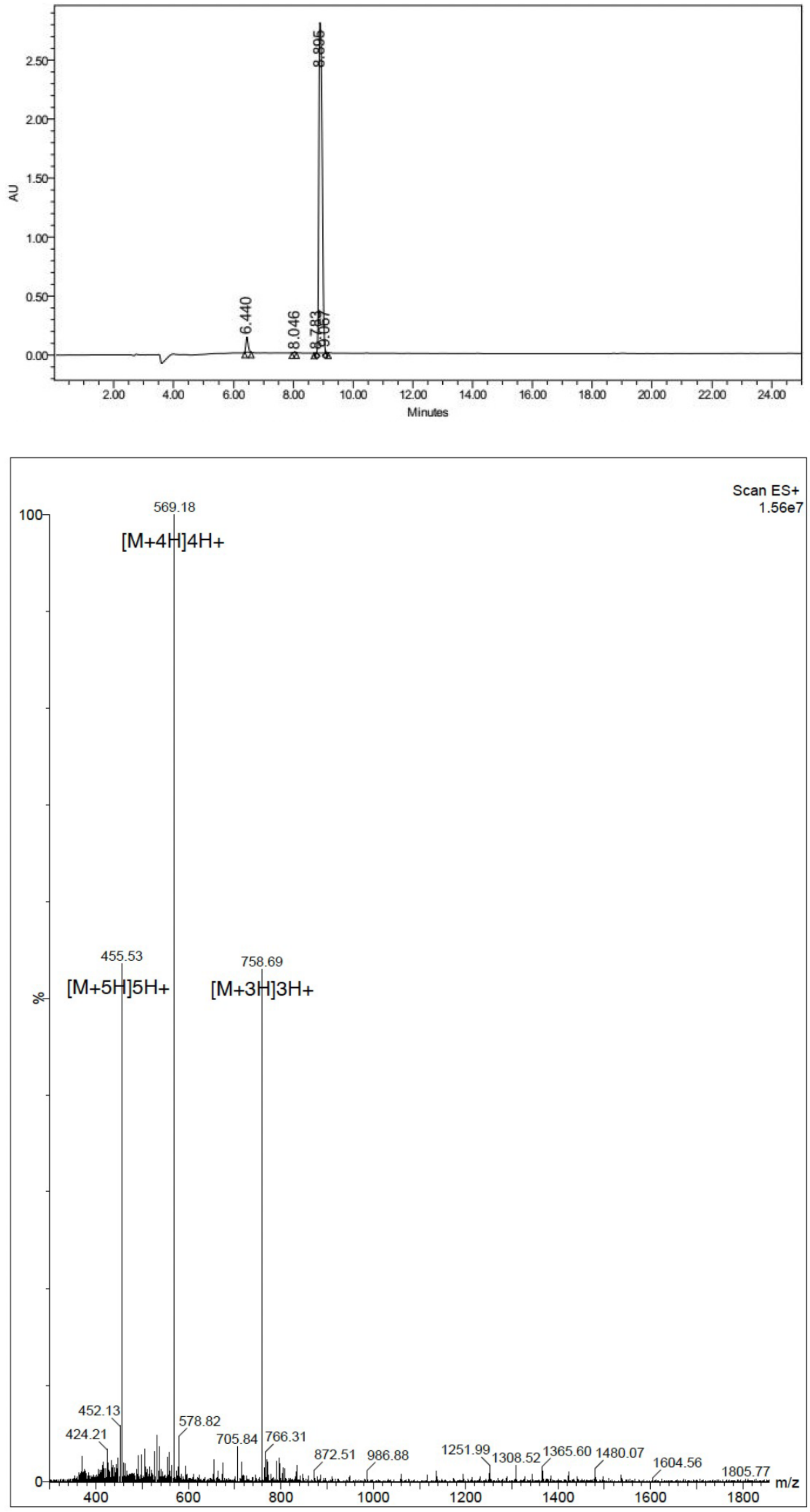


**Figure S23:** Validation of synthesized peptide S06. **a**, HPLC chromatography. **b**, Mass spectrometry.

**a**

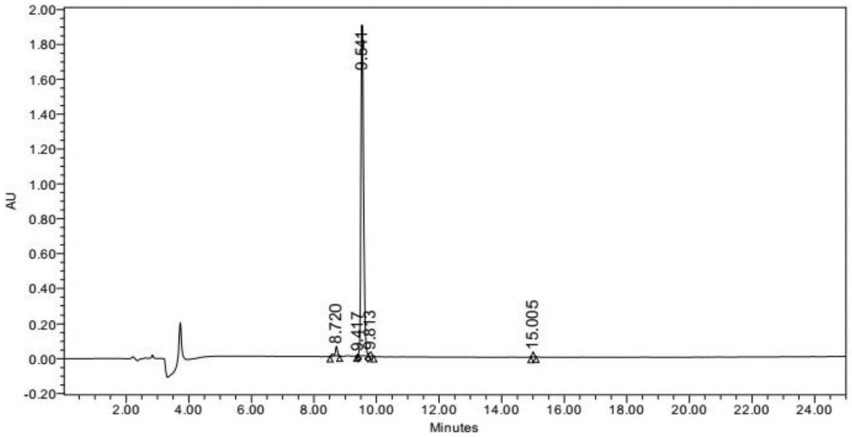


**b**

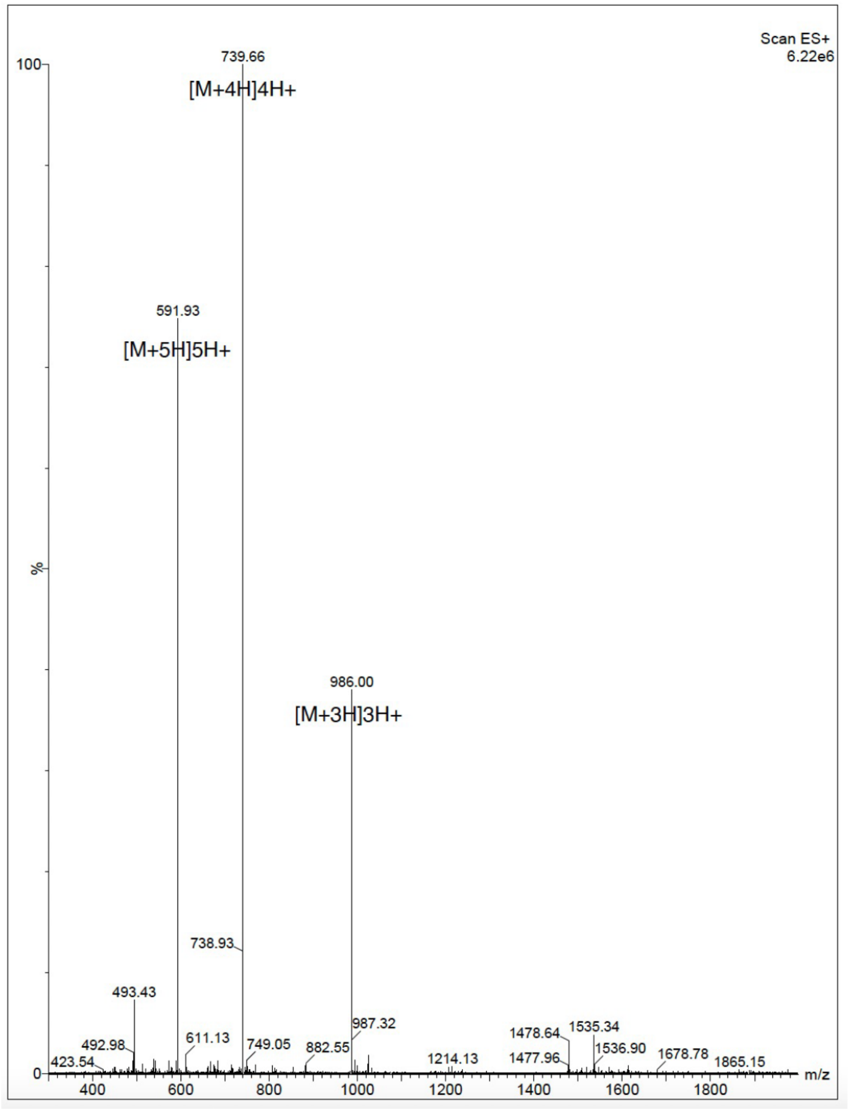


**Figure S24:** Validation of synthesized peptide S07. **a**, HPLC chromatography. **b**, Mass spectrometry.

a

b

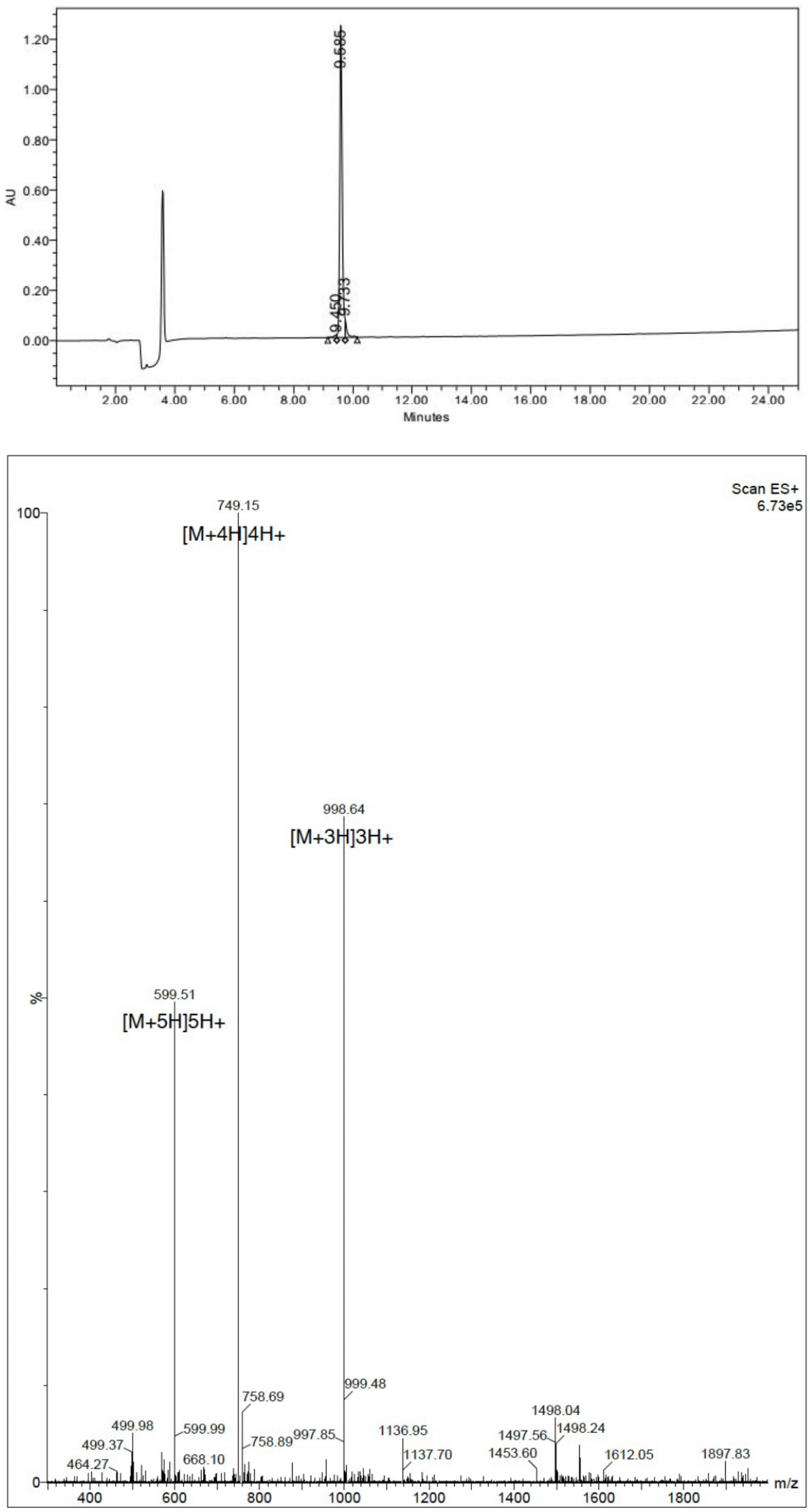


**Figure S25:** Validation of synthesized peptide S08. **a**, HPLC chromatography. **b**, Mass spectrometry.

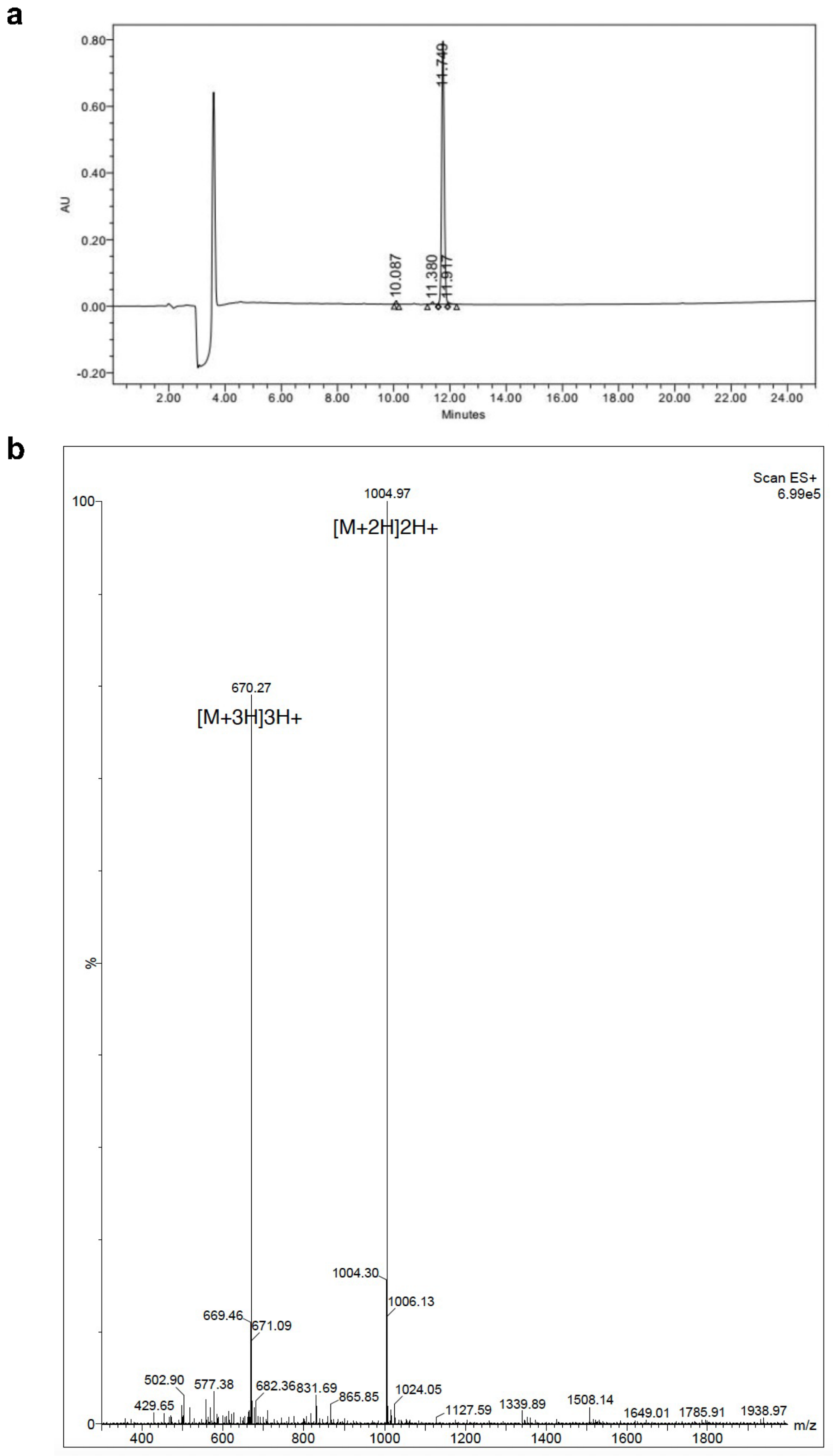


**Figure S26:** Validation of synthesized peptide S09. **a**, HPLC chromatography. **b**, Mass spectrometry.

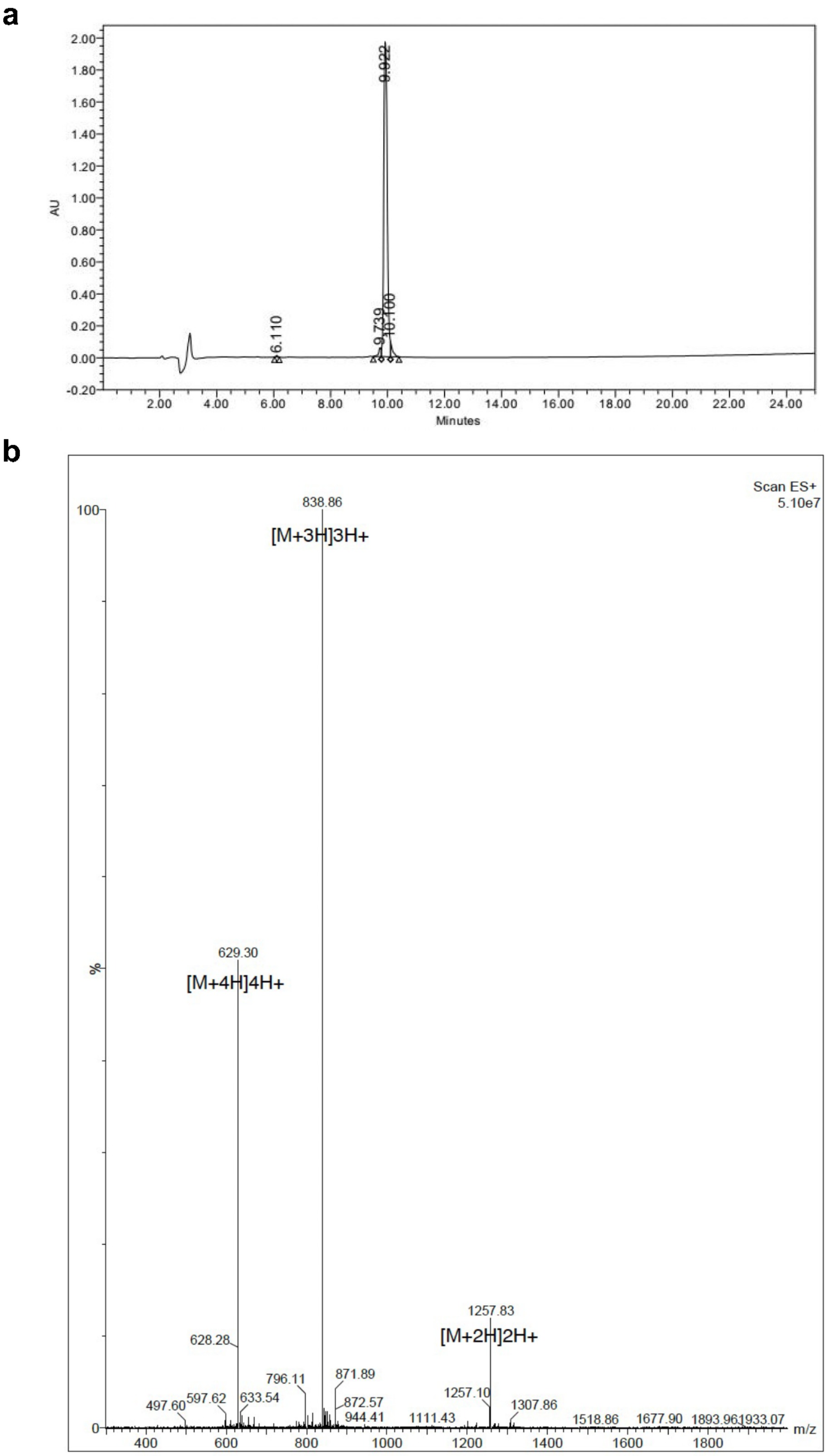


**Figure S27:** Validation of synthesized peptide S10. **a**, HPLC chromatography. **b**, Mass spectrometry.

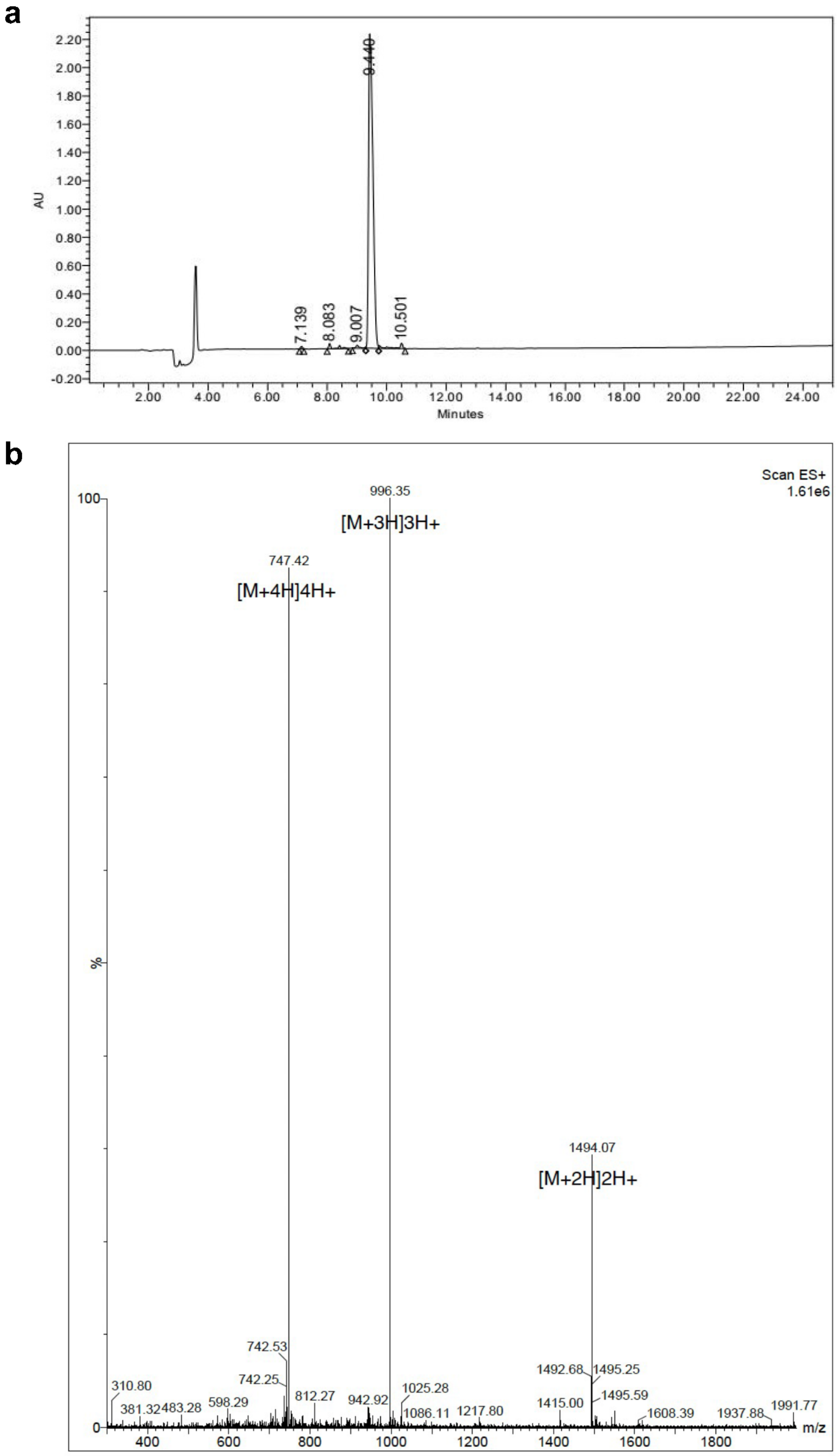


**Figure S28:** Validation of synthesized peptide R01. **a**, HPLC chromatography. **b**, Mass spectrometry.

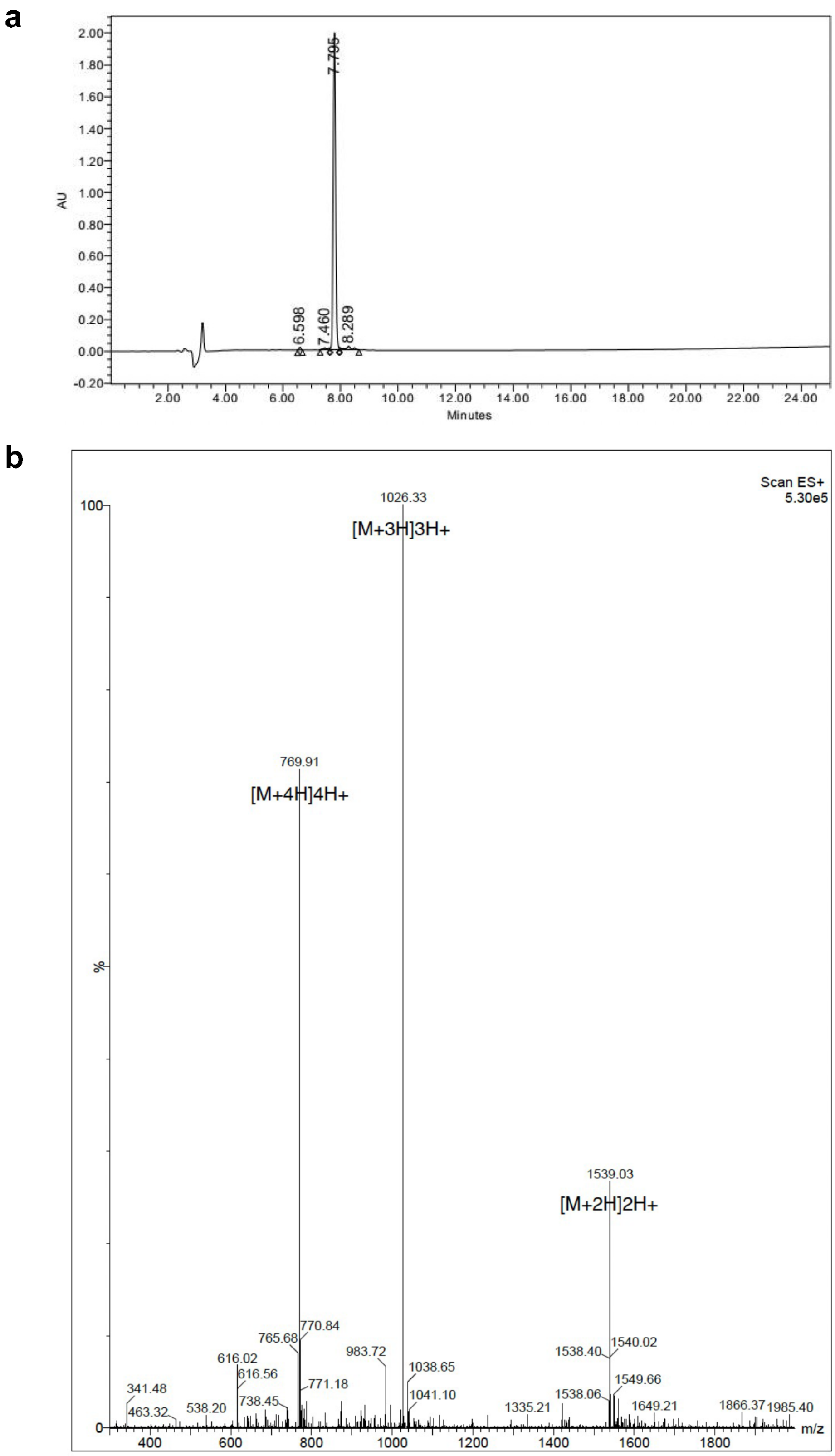


**Figure S29:** Validation of synthesized peptide R02. **a**, HPLC chromatography. **b**, Mass spectrometry.

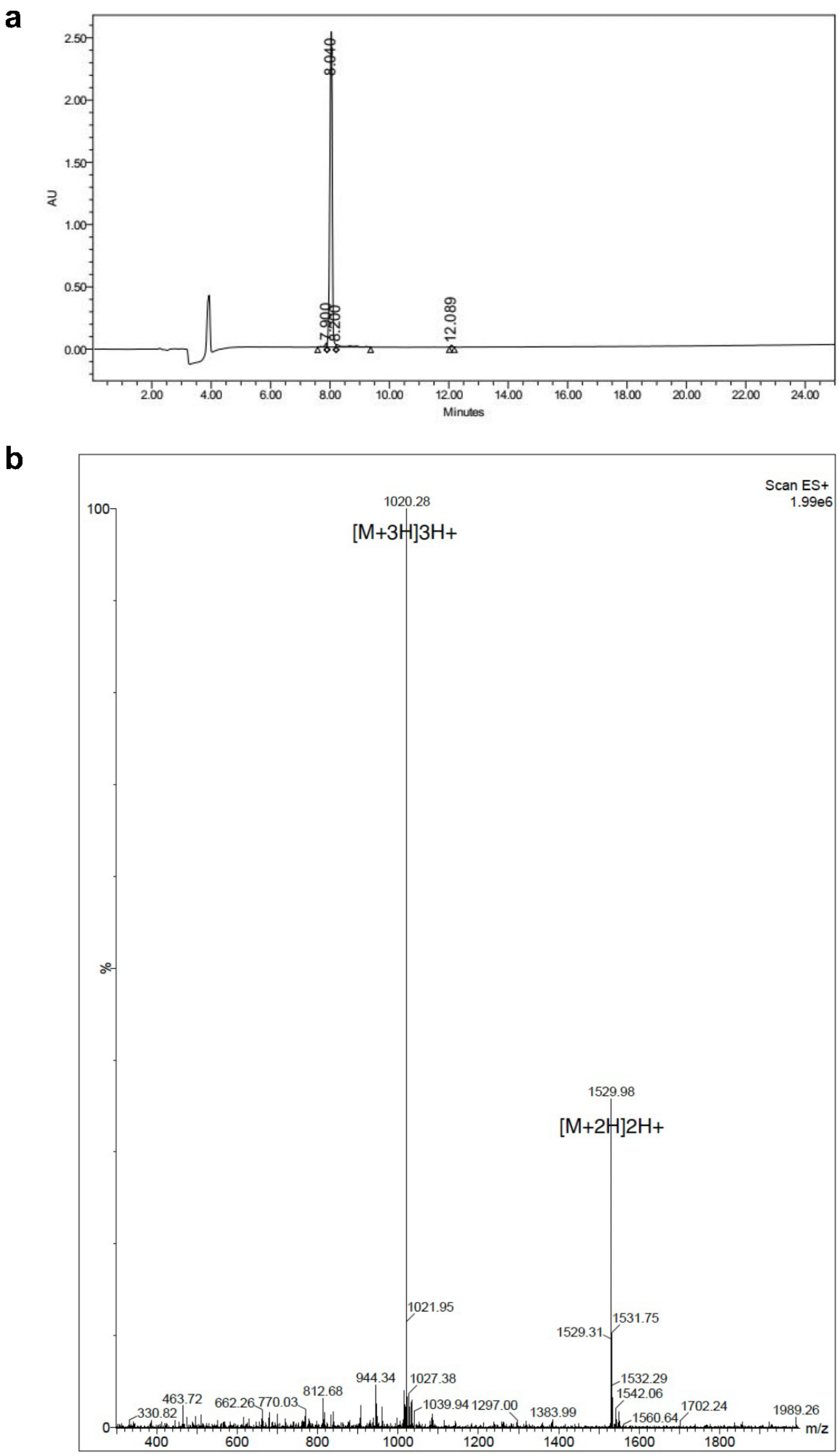


**Figure S30:** Validation of synthesized peptide R03. **a**, HPLC chromatography. **b**, Mass spectrometry.

a

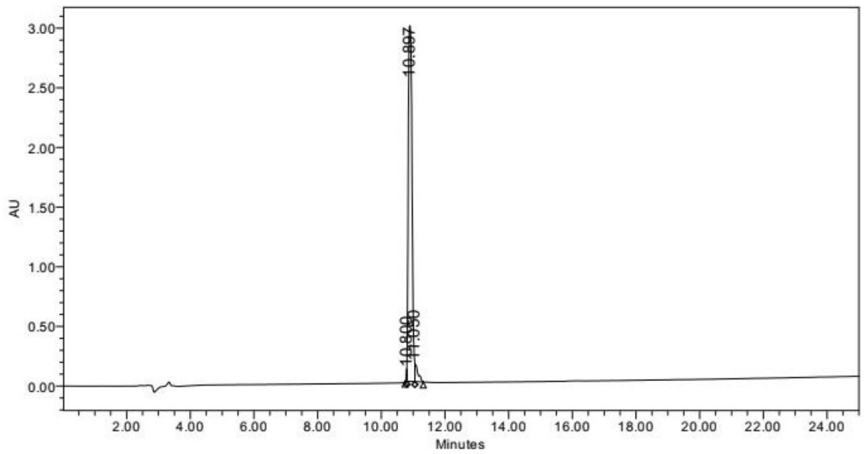


b

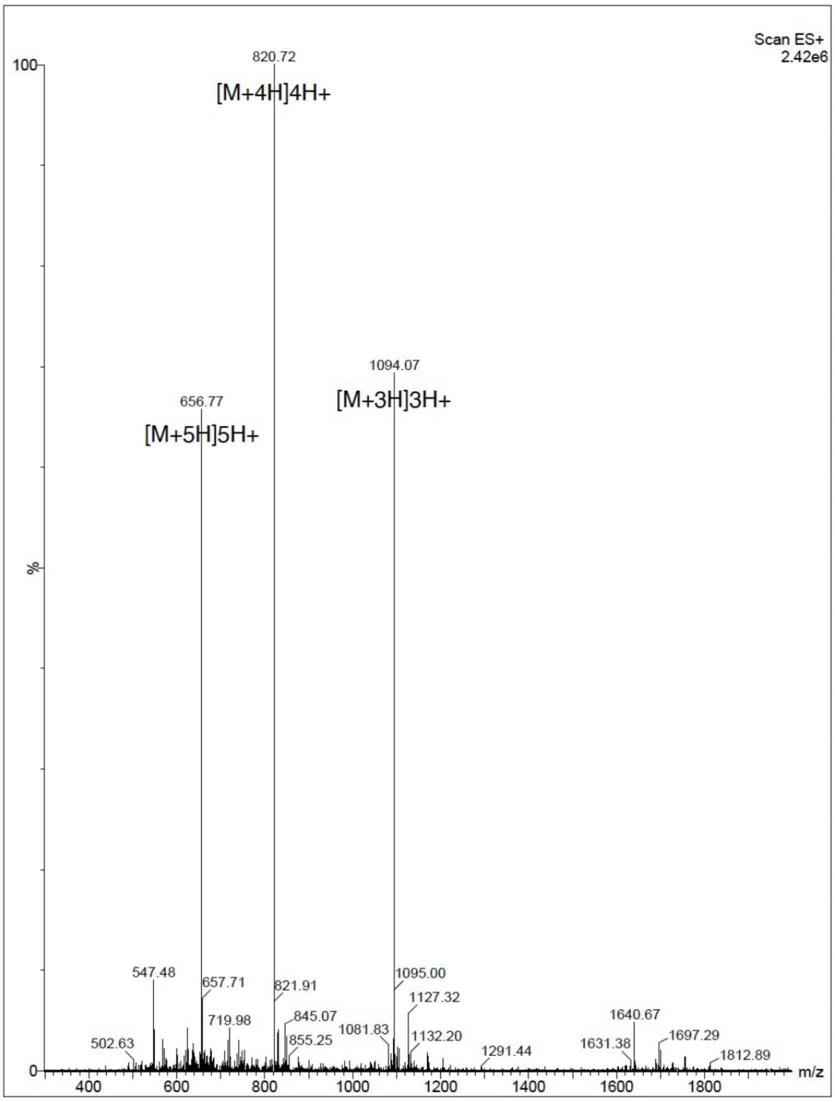


**Figure S31:** Validation of synthesized peptide R04. **a**, HPLC chromatography. **b**, Mass spectrometry.

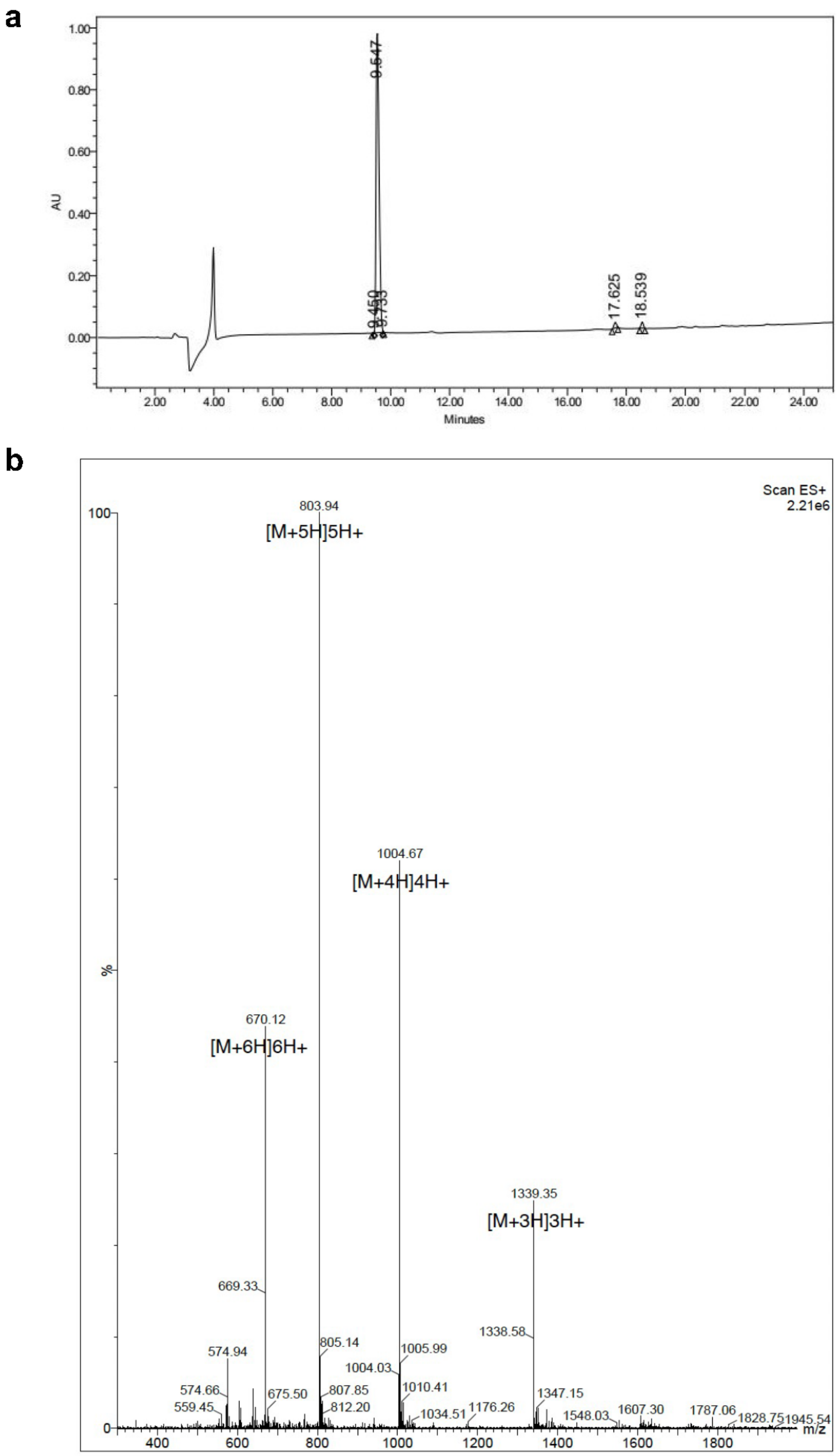


**Figure S32:** Validation of synthesized peptide R05. **a**, HPLC chromatography. **b**, Mass spectrometry.

**a**

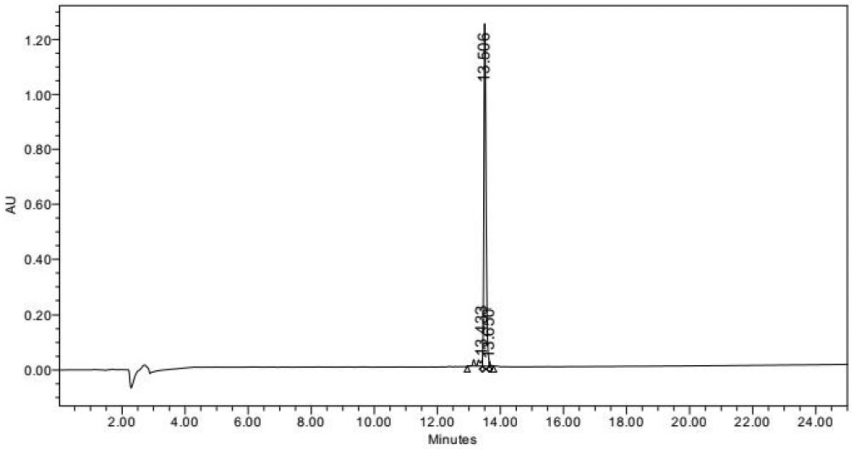


**b**

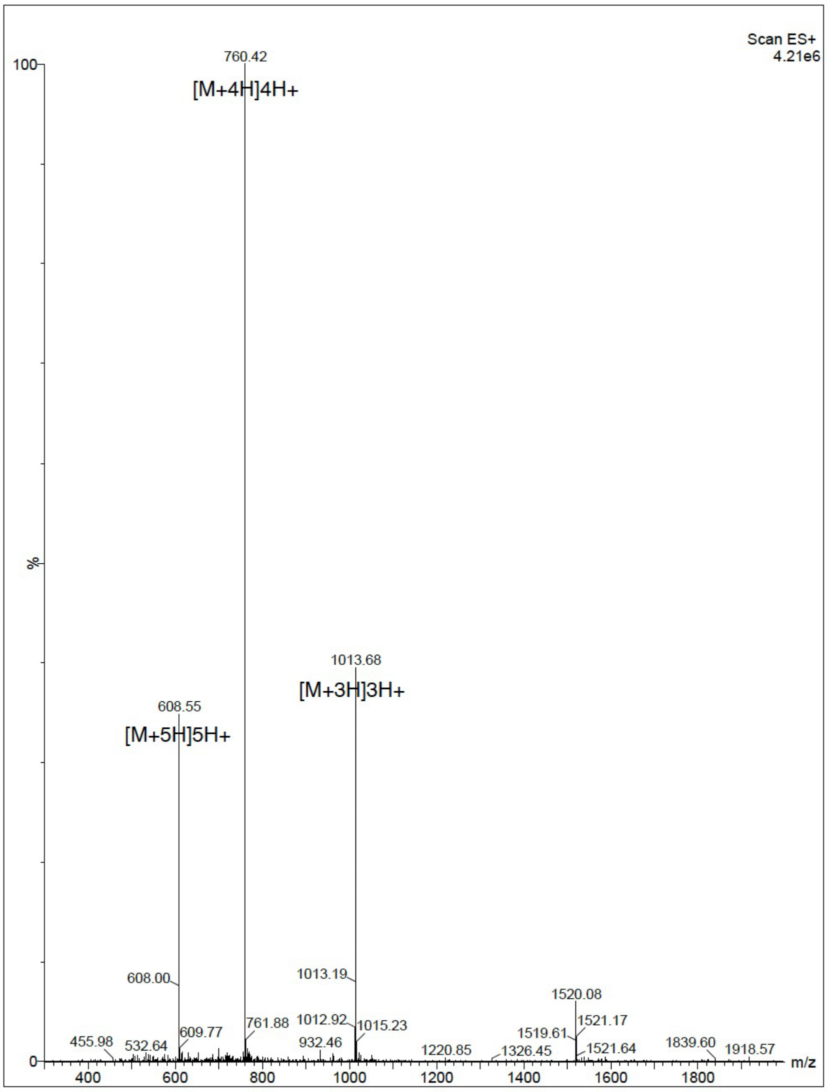


**Figure S33:** Validation of synthesized peptide R06. **a**, HPLC chromatography. **b**, Mass spectrometry.

a

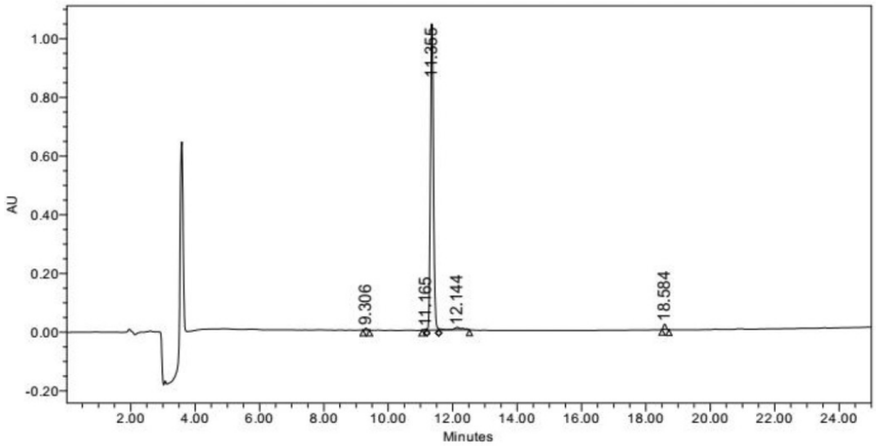


b

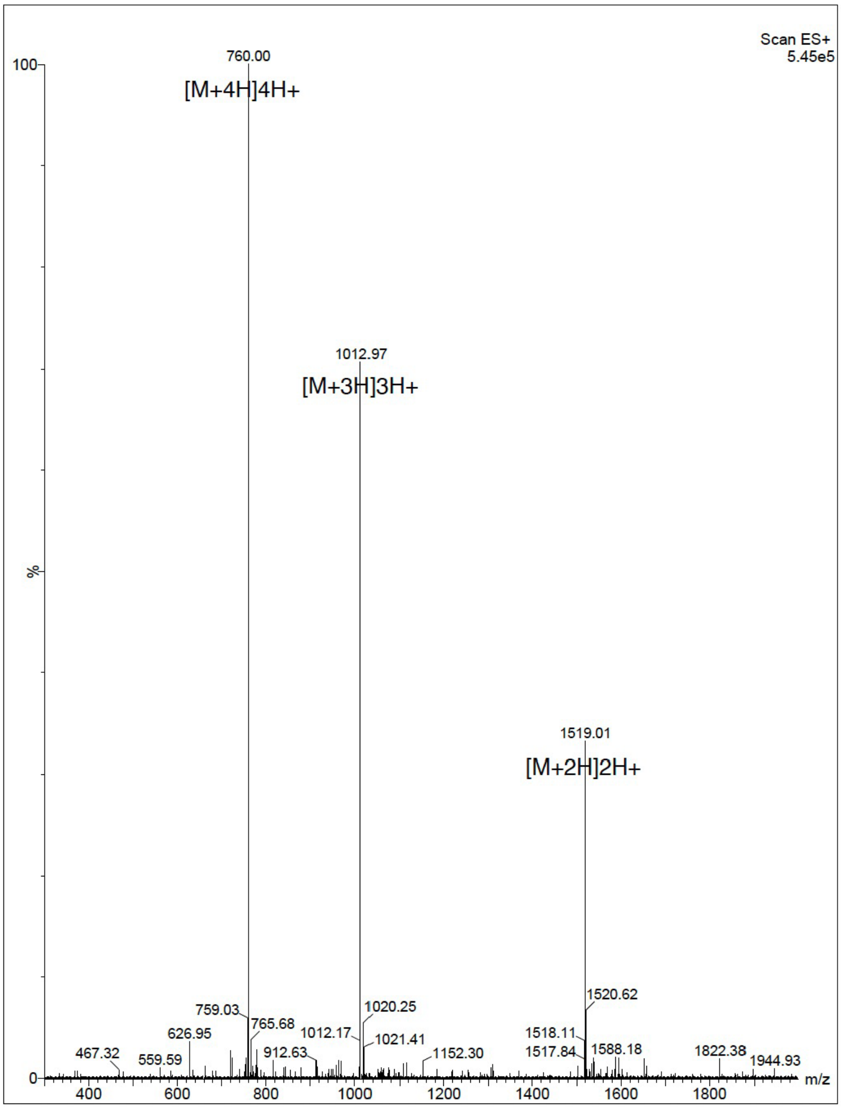


**Figure S34:** Validation of synthesized peptide R07. **a**, HPLC chromatography. **b**, Mass spectrometry.

**a**

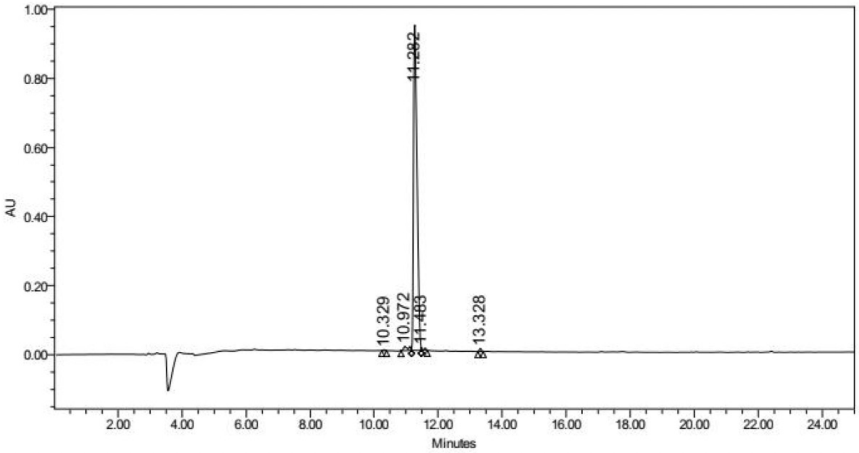


**b**

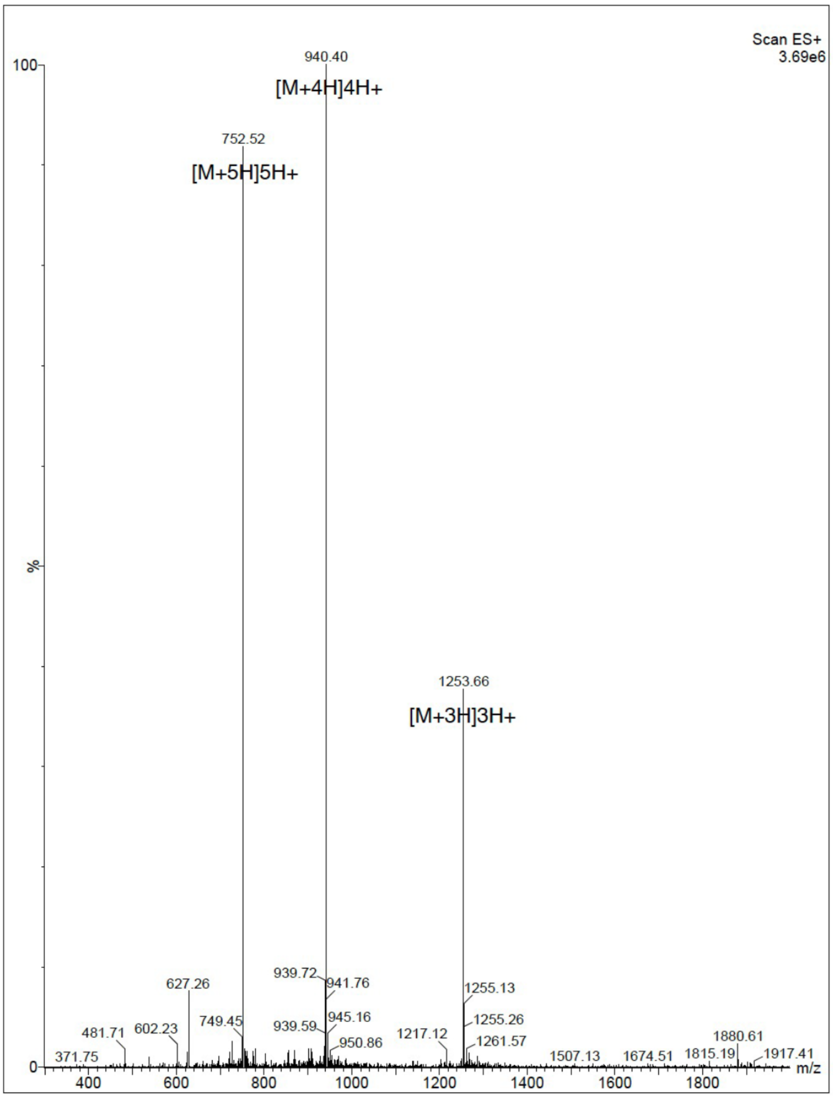


**Figure S35:** Validation of synthesized peptide R08. **a**, HPLC chromatography. **b**, Mass spectrometry.

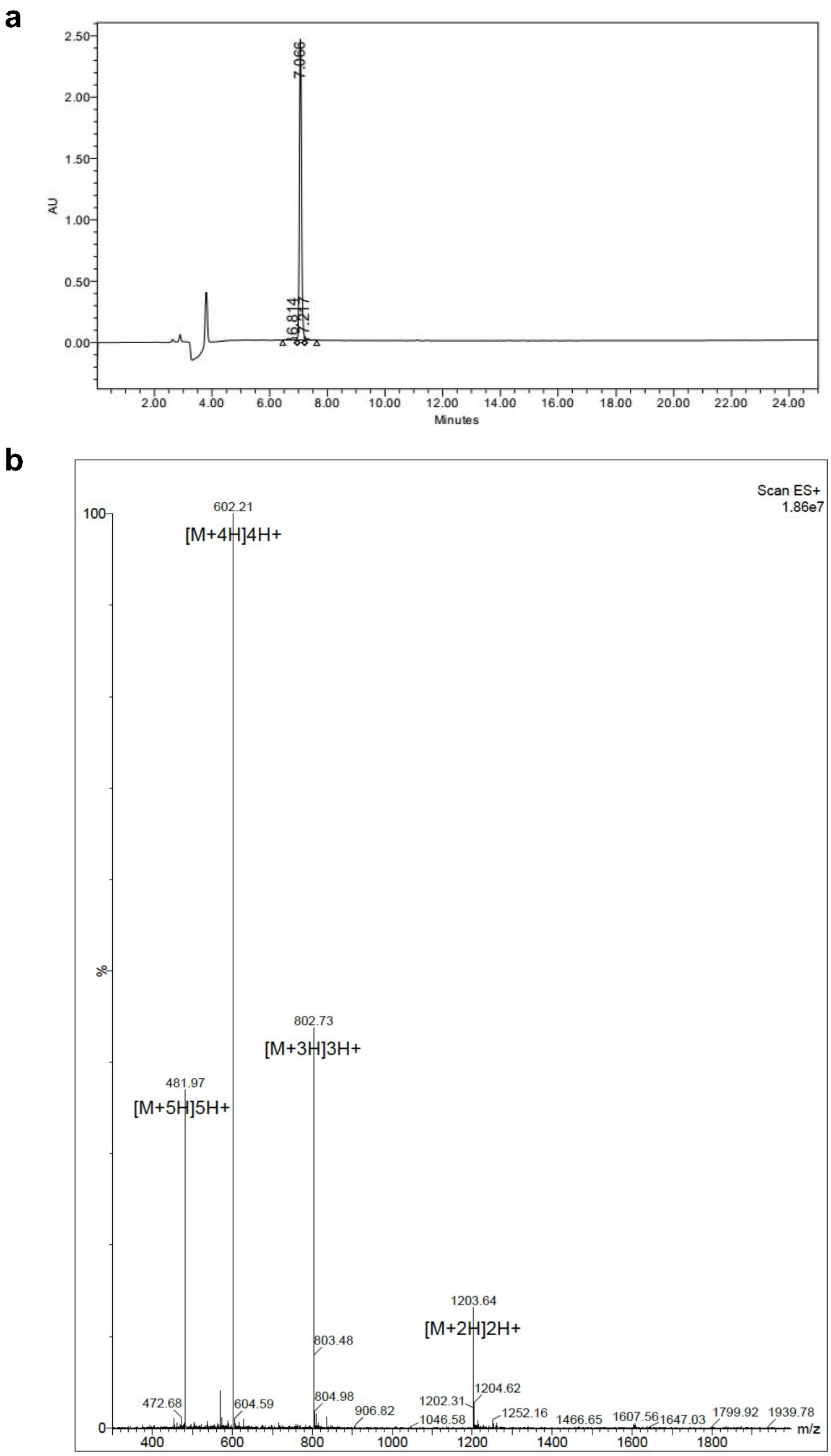


**Figure S36:** Validation of synthesized peptide R09. **a**, HPLC chromatography. **b**, Mass spectrometry.

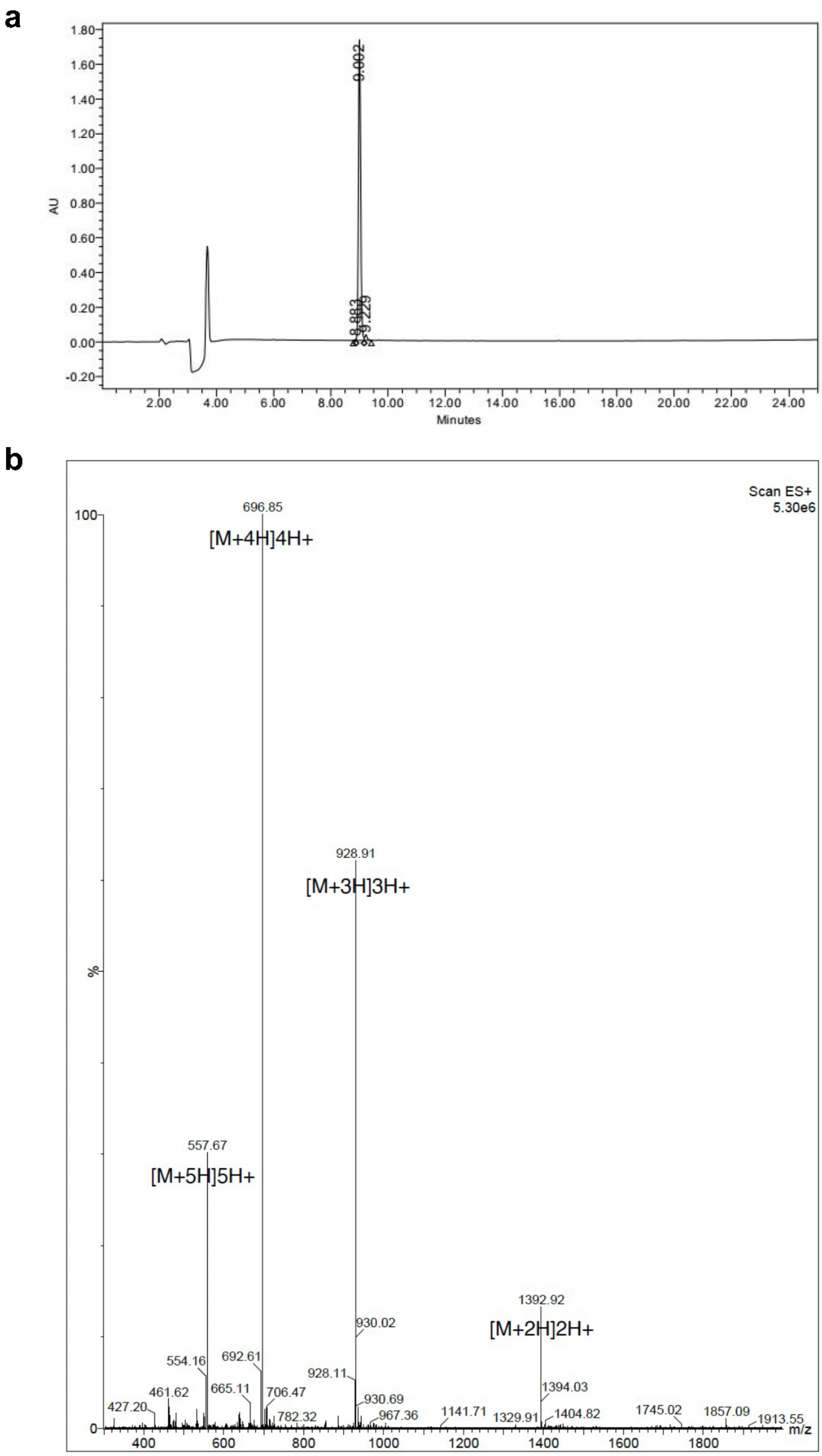


**Figure S37:** Validation of synthesized peptide R10. **a**, HPLC chromatography. **b**, Mass spectrometry.

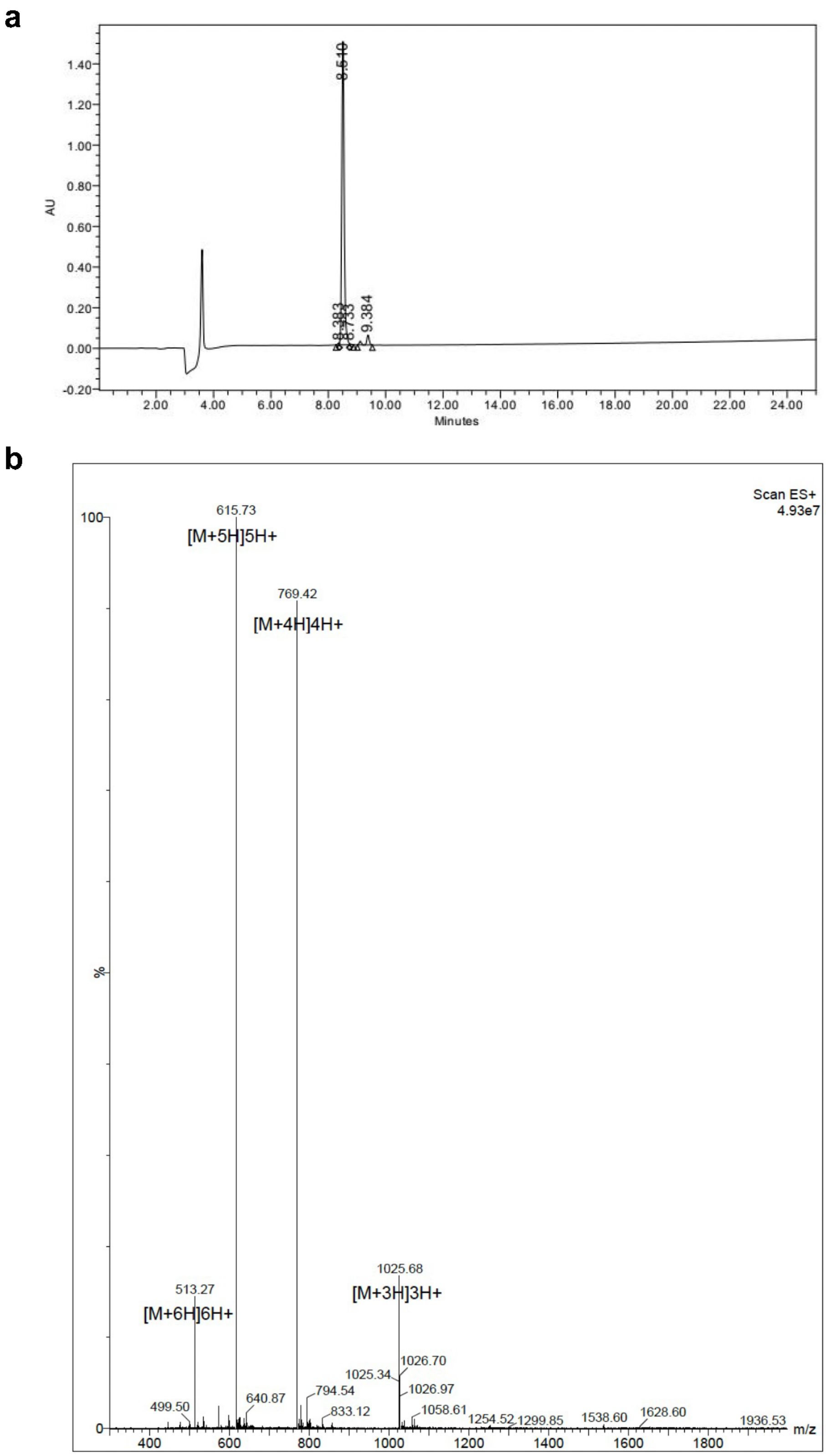


**Figure S38:** Validation of synthesized peptide A01ori. **a**, HPLC chromatography. **b**, Mass spectrometry.

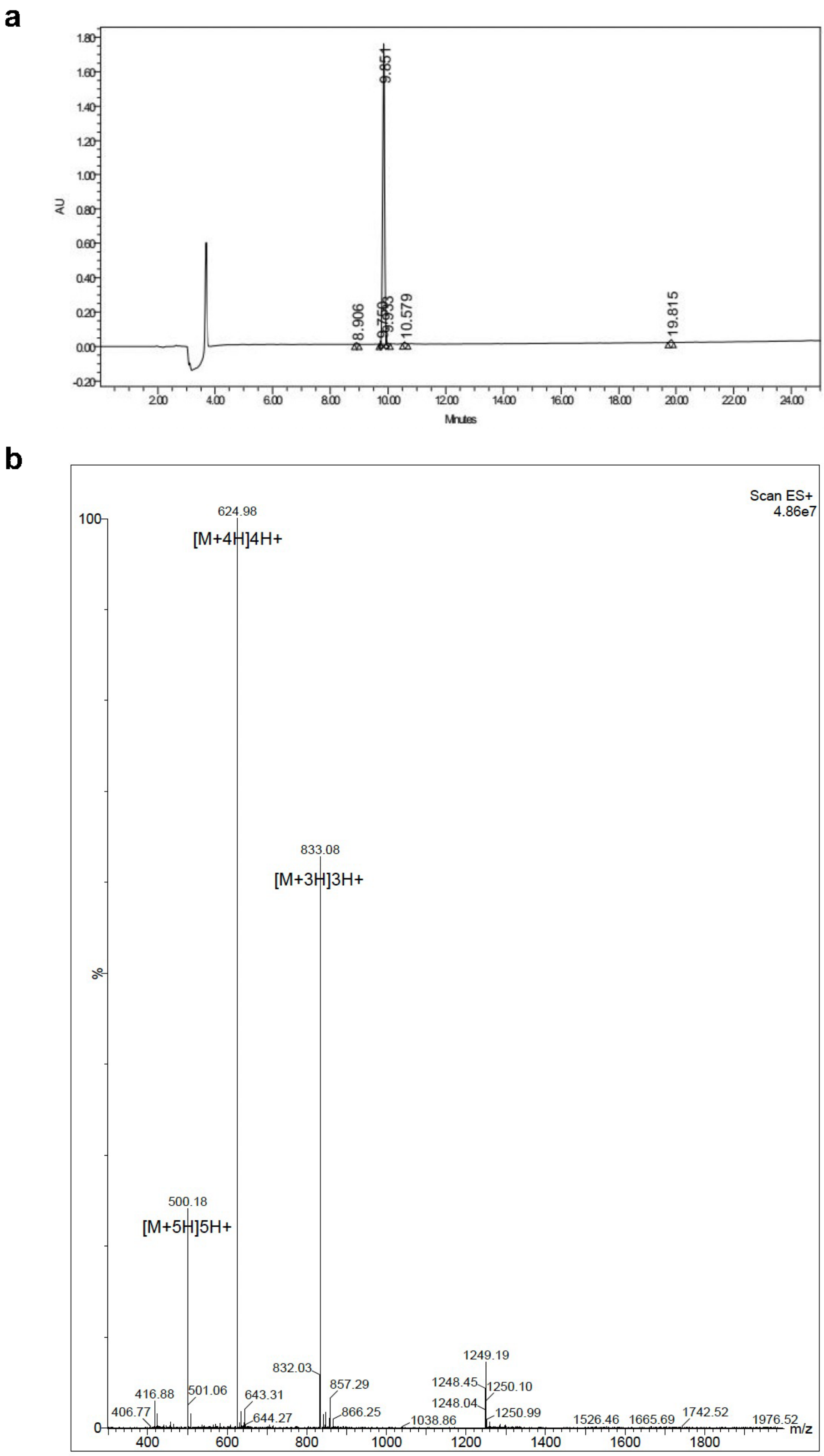


**Figure S39:** Validation of synthesized peptide A02ori. **a**, HPLC chromatography. **b**, Mass spectrometry.

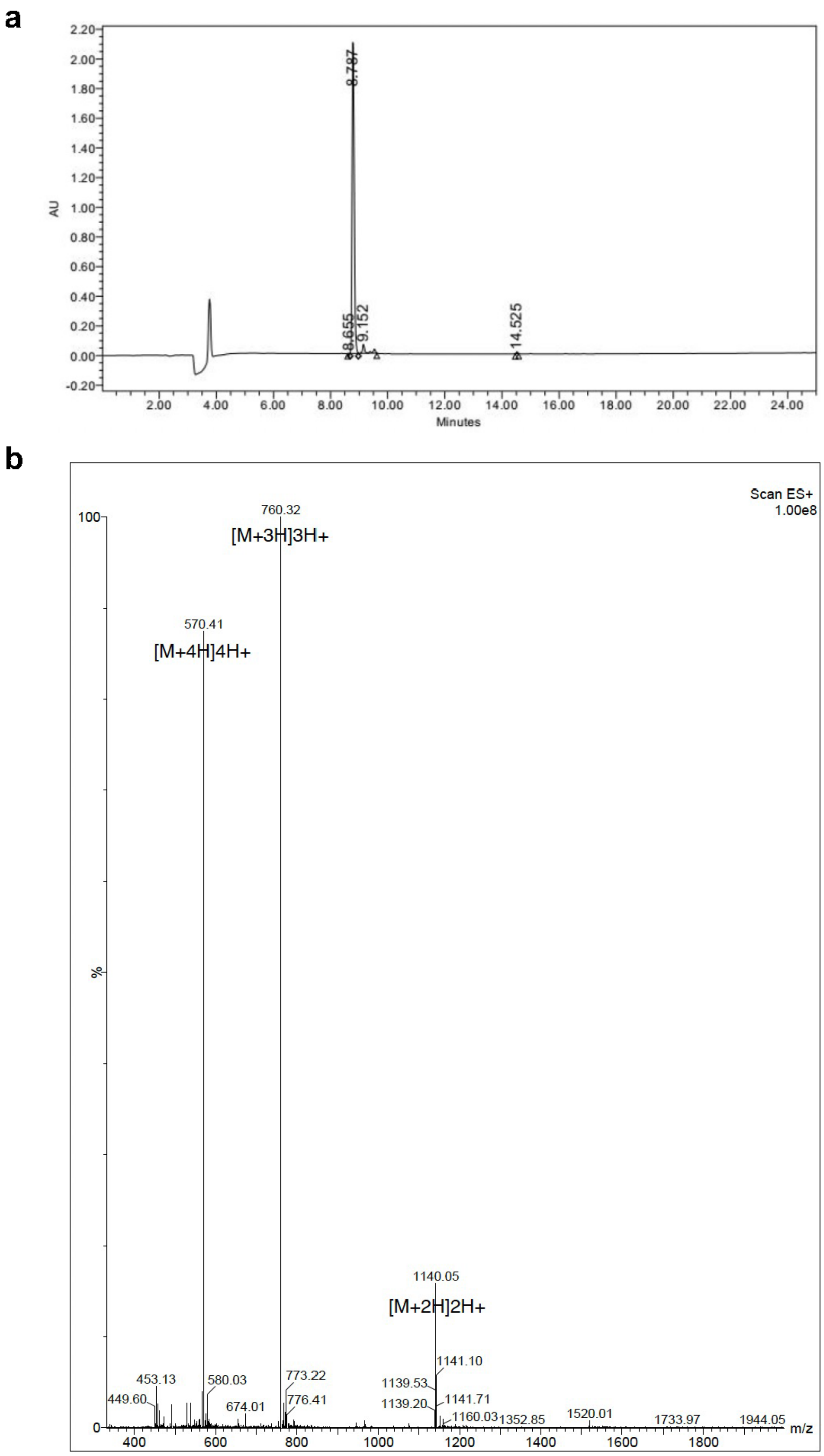


**Figure S40:** Validation of synthesized peptide A04ori. **a**, HPLC chromatography. **b**, Mass spectrometry.

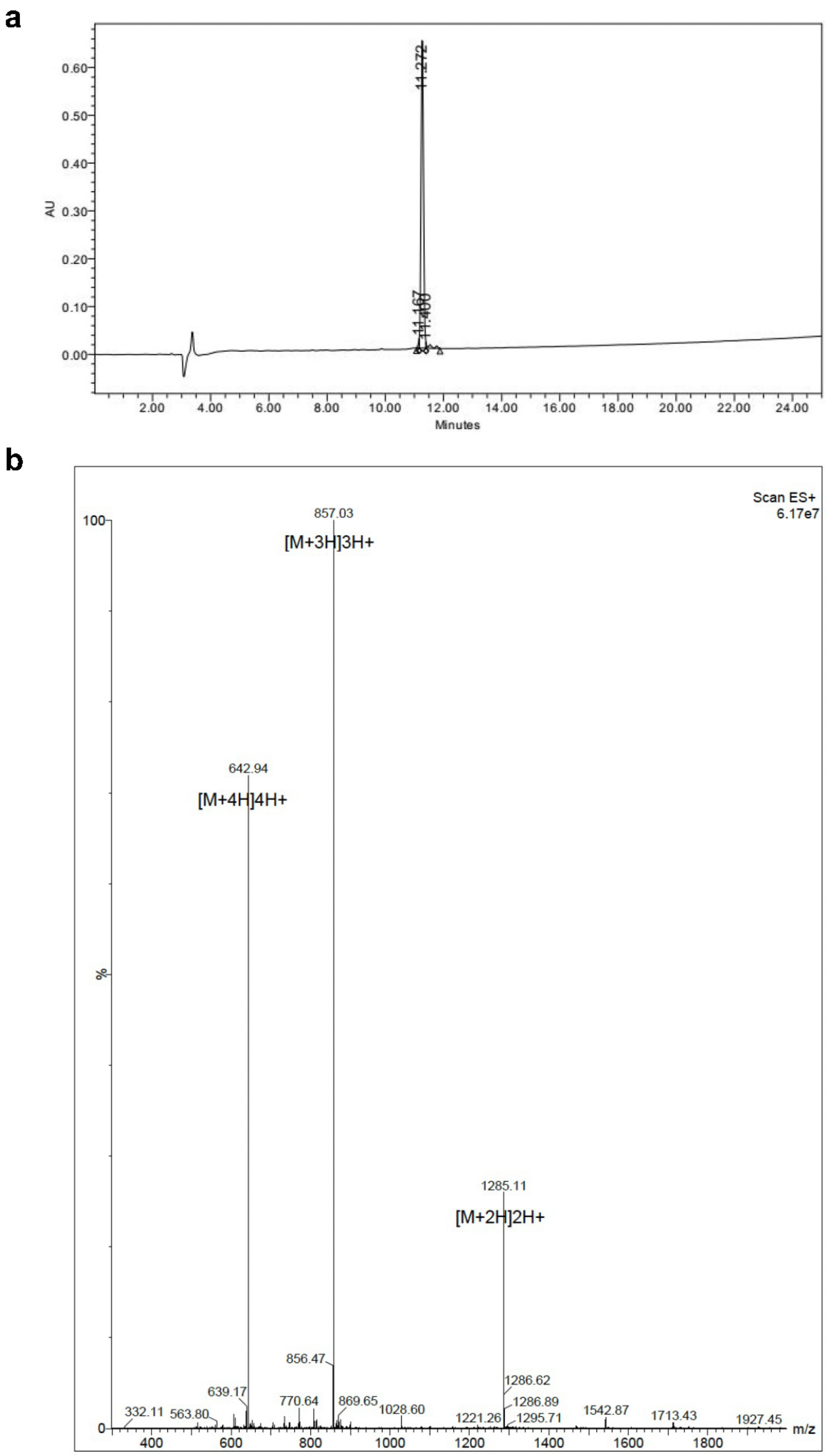


**Figure S41:** Validation of synthesized peptide A05ori. **a**, HPLC chromatography. **b**, Mass spectrometry.

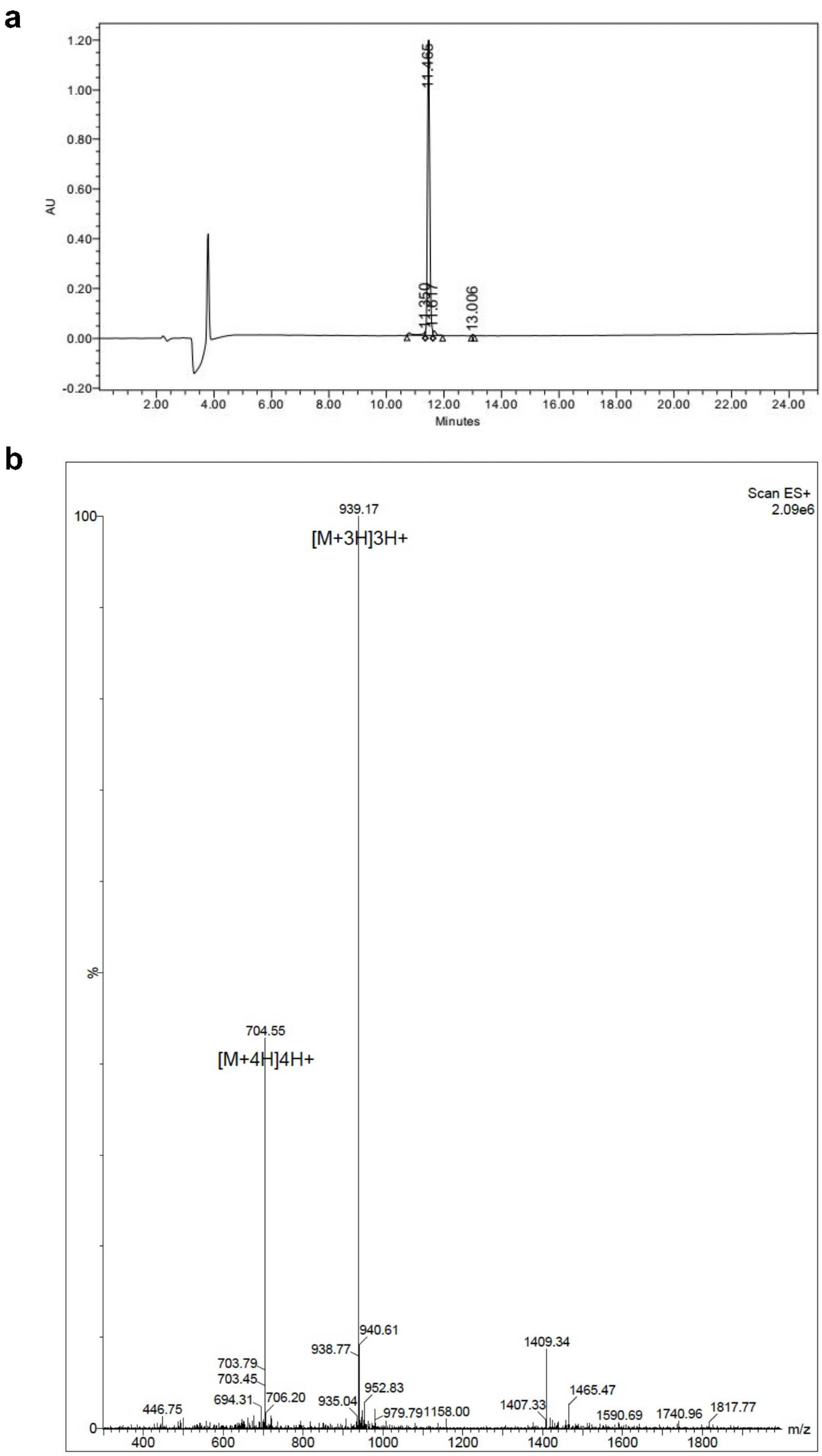


**Figure S42:** Validation of synthesized peptide A08ori. **a**, HPLC chromatography. **b**, Mass spectrometry.

a

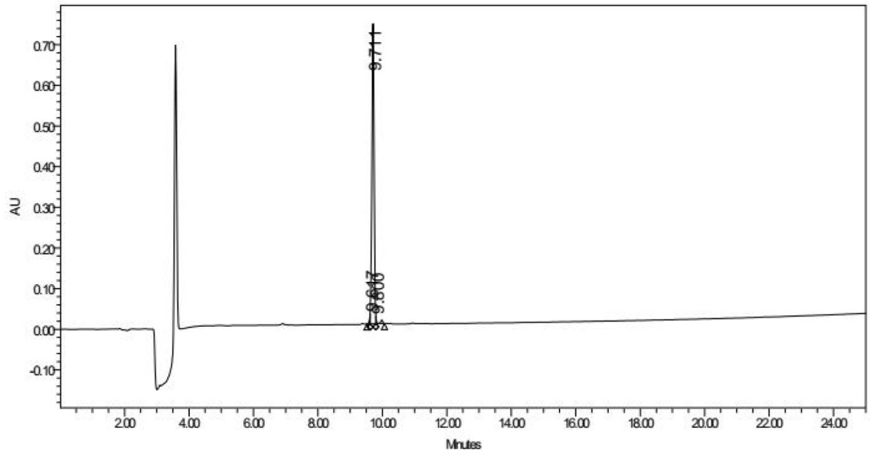


b

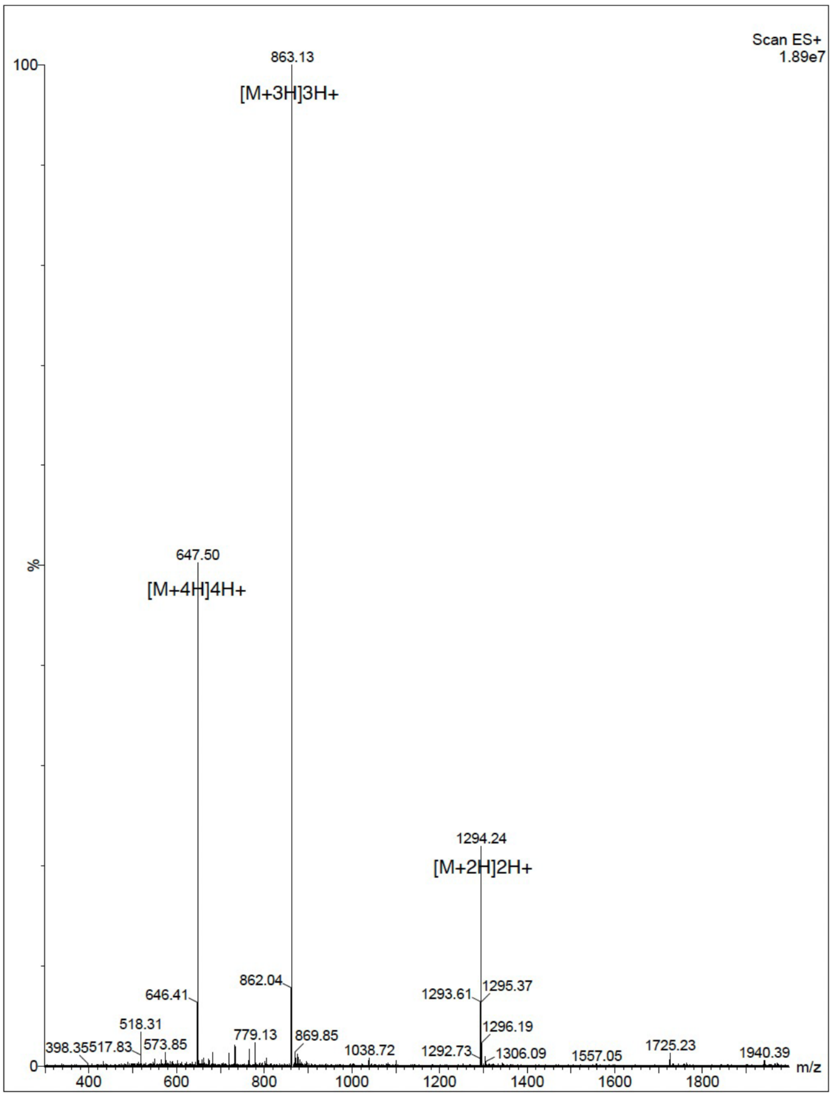


**Figure S43:** Validation of synthesized peptide A09ori. **a**, HPLC chromatography. **b**, Mass spectrometry.

**a**

**b**

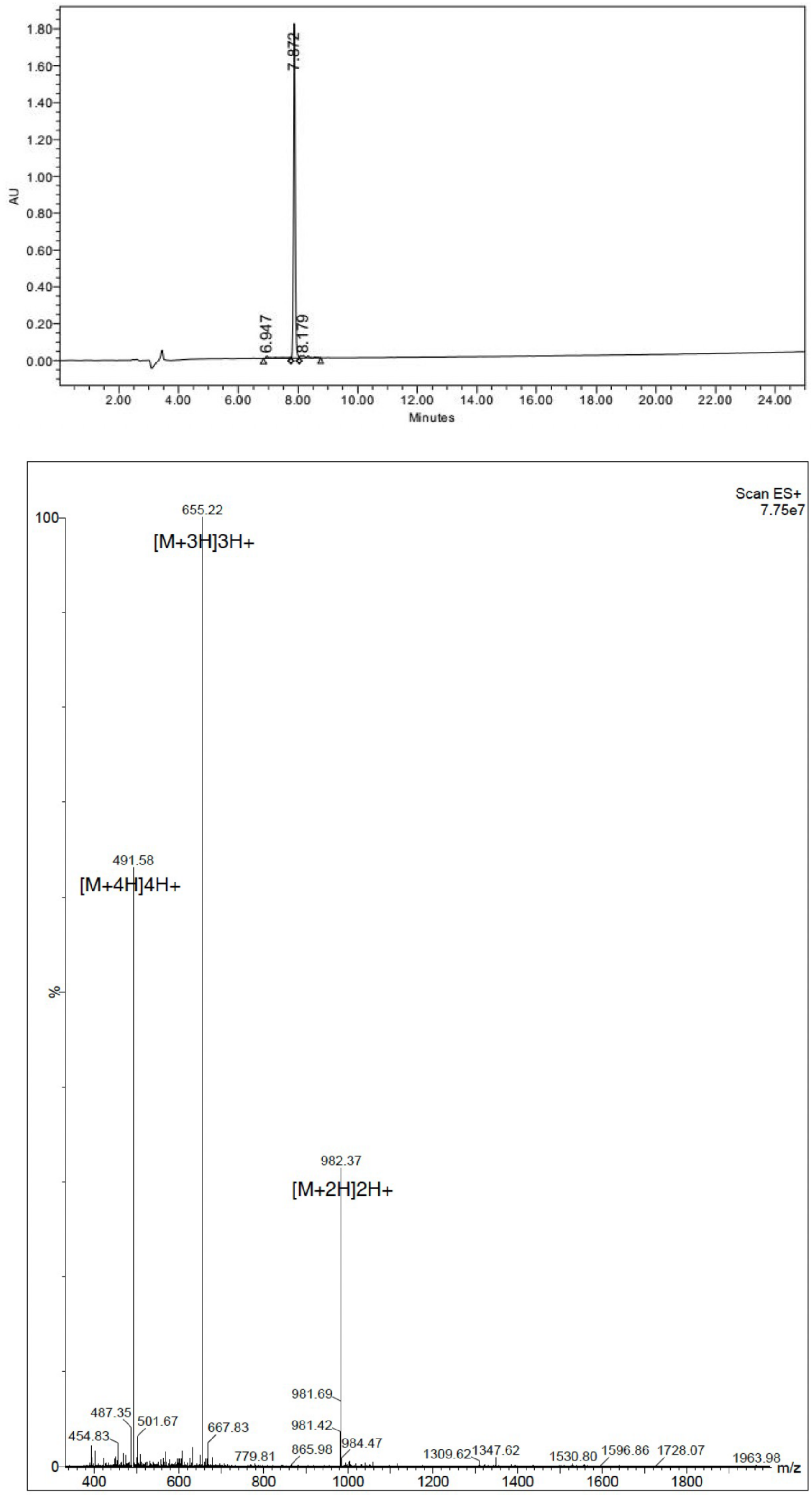


**Figure S44:** Validation of synthesized peptide A10ori. **a**, HPLC chromatography. **b**, Mass spectrometry.

a

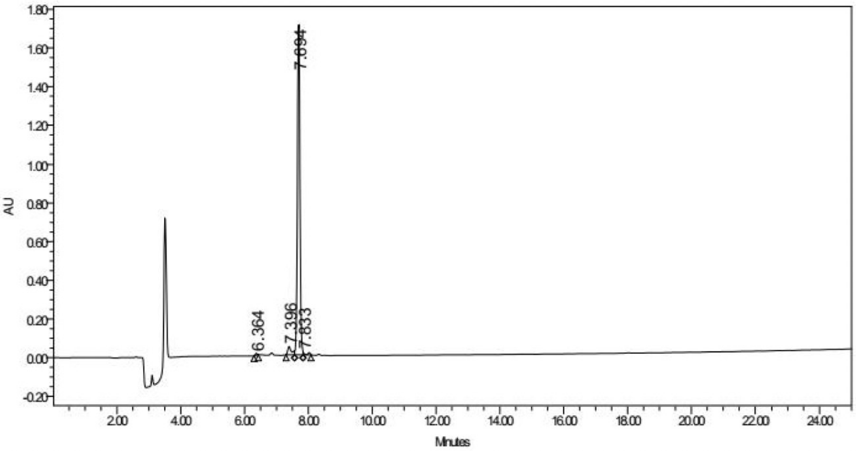


b

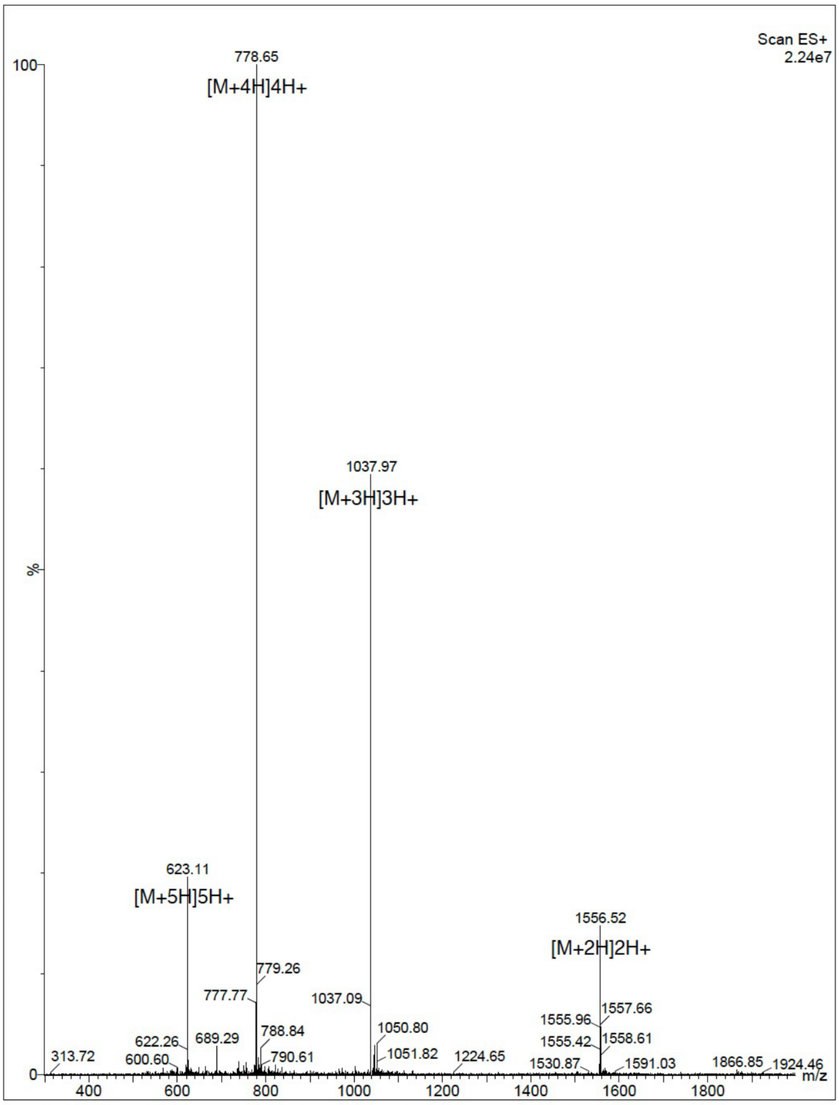


**Figure S45:** Validation of synthesized peptide 2-0. **a**, HPLC chromatography. **b**, Mass spectrometry.

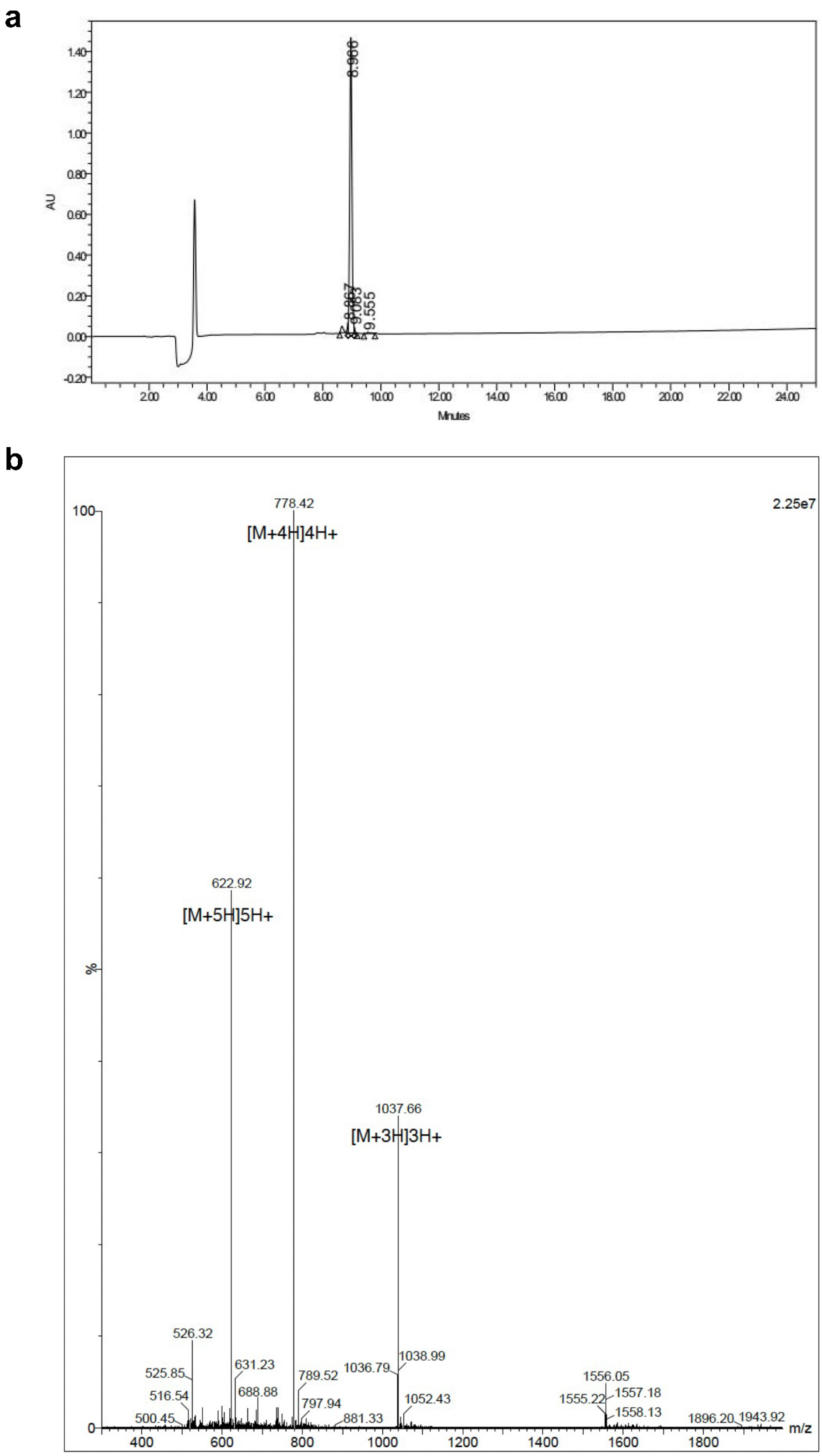


**Figure S46:** Validation of synthesized peptide 2-1. **a**, HPLC chromatography. **b**, Mass spectrometry.

a

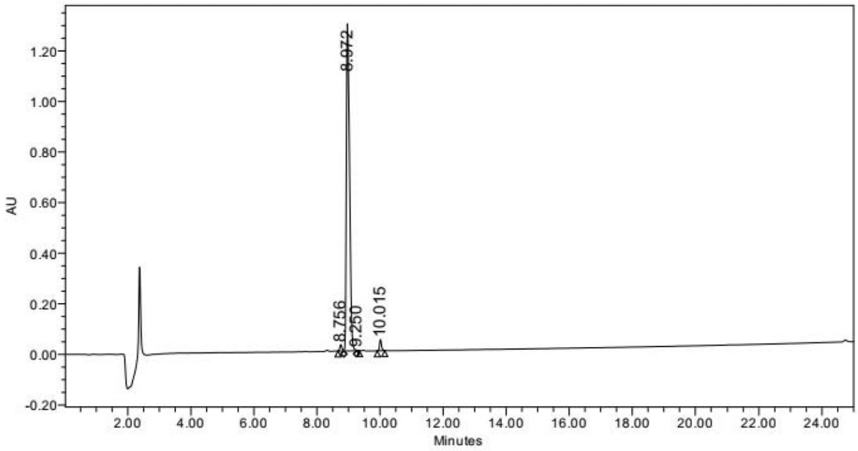


b

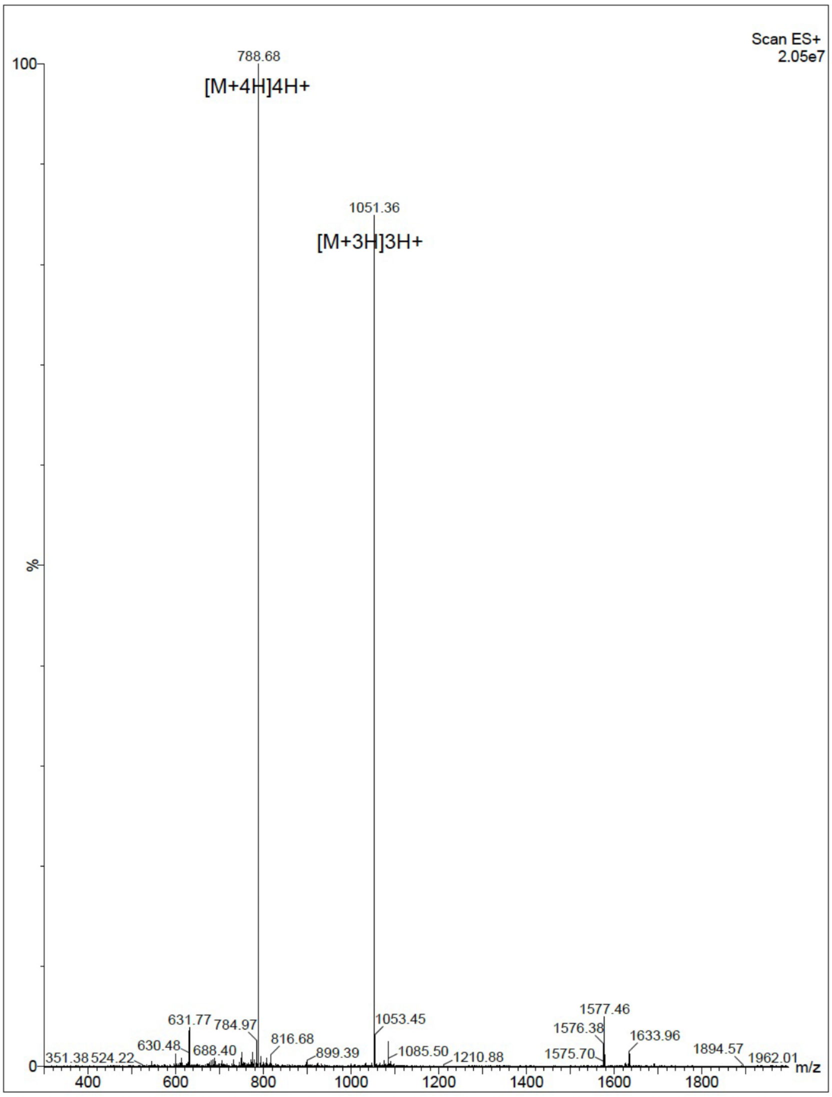


**Figure S47:** Validation of synthesized peptide 2-2. **a**, HPLC chromatography. **b**, Mass spectrometry.

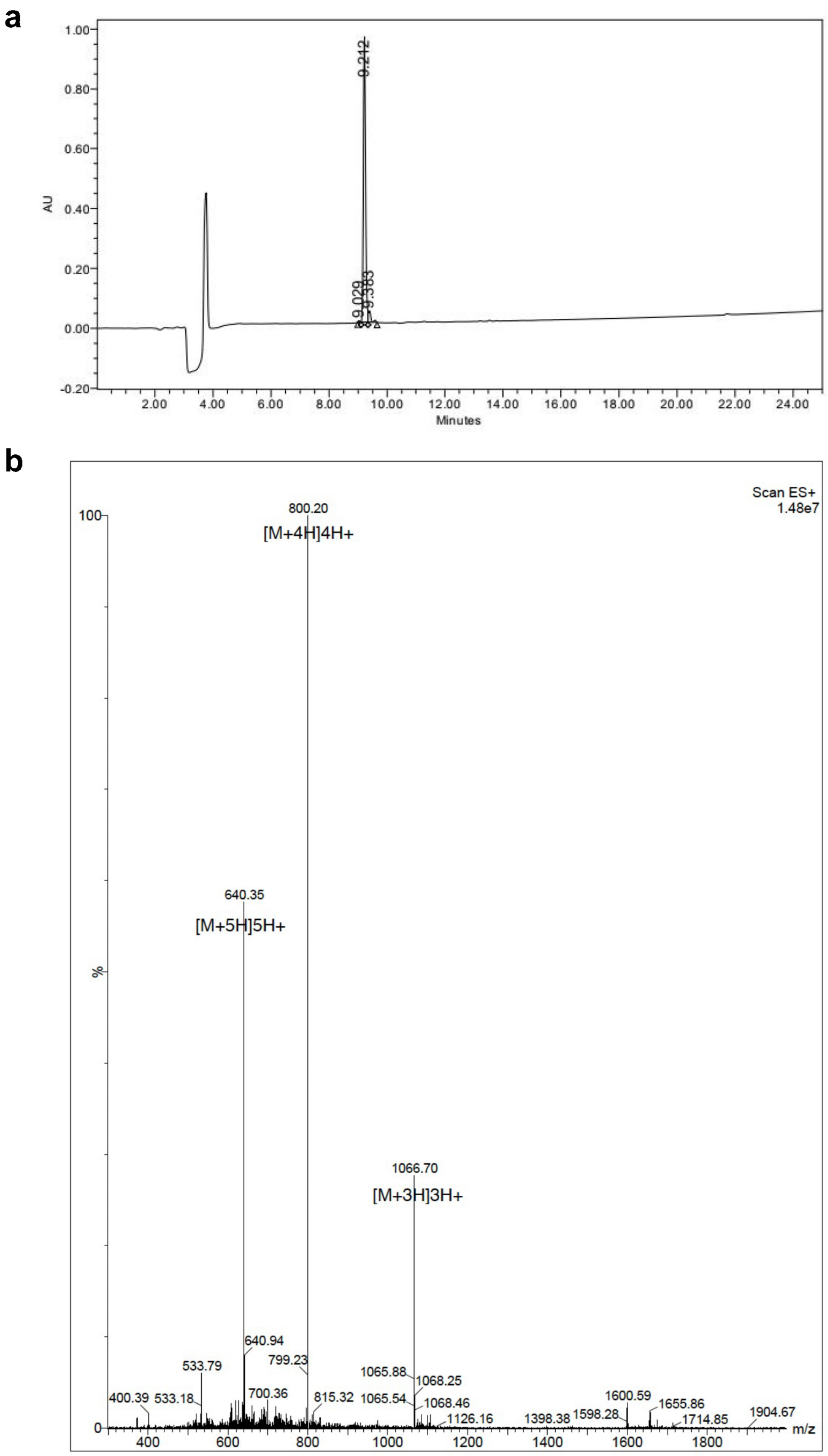


**Figure S48:** Validation of synthesized peptide 2-4. **a**, HPLC chromatography. **b**, Mass spectrometry.

**a**

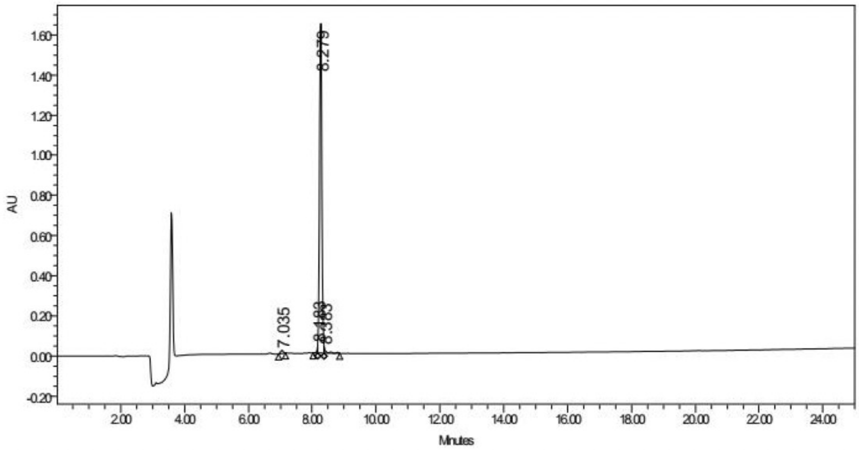


**b**

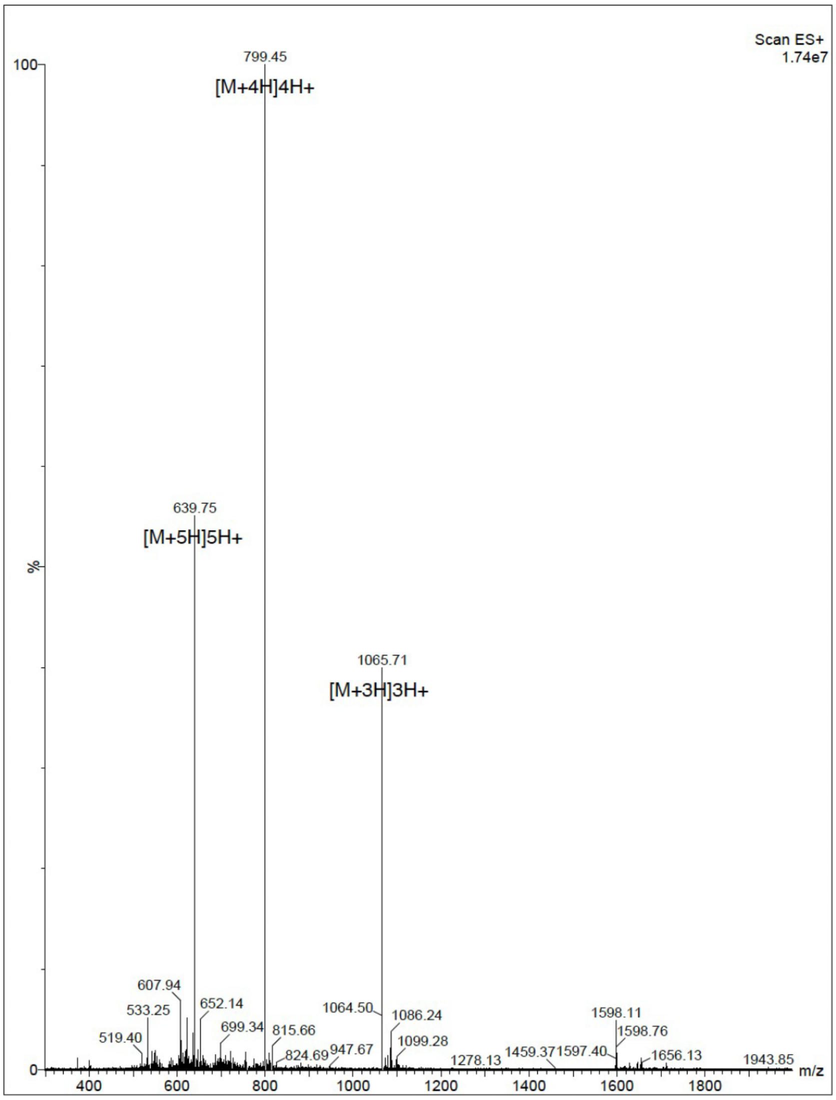


**Figure S49:** Validation of synthesized peptide 2-5. **a**, HPLC chromatography. **b**, Mass spectrometry.

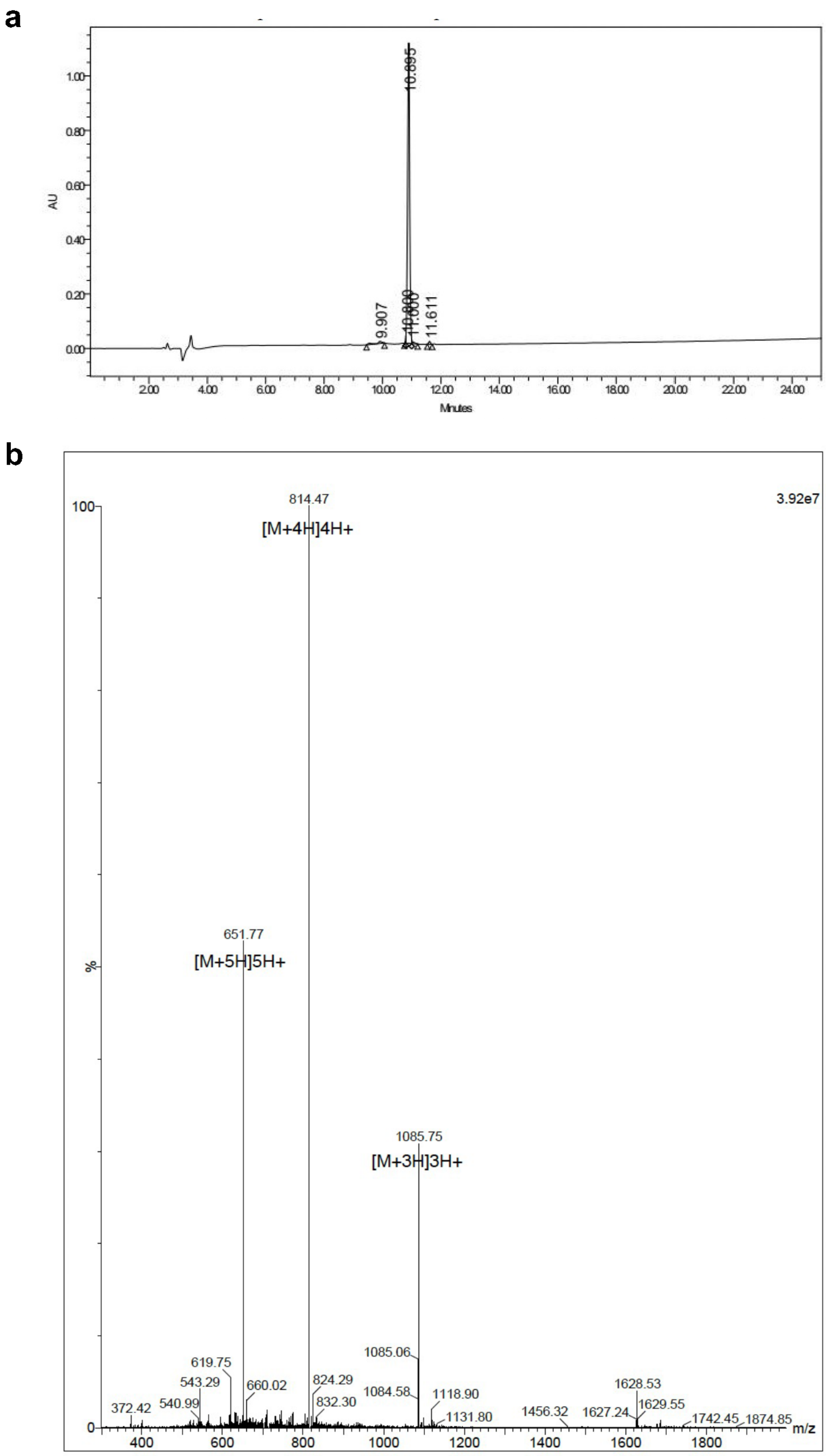


**Figure S50:** Validation of synthesized peptide 2-6. **a**, HPLC chromatography. **b**, Mass spectrometry.

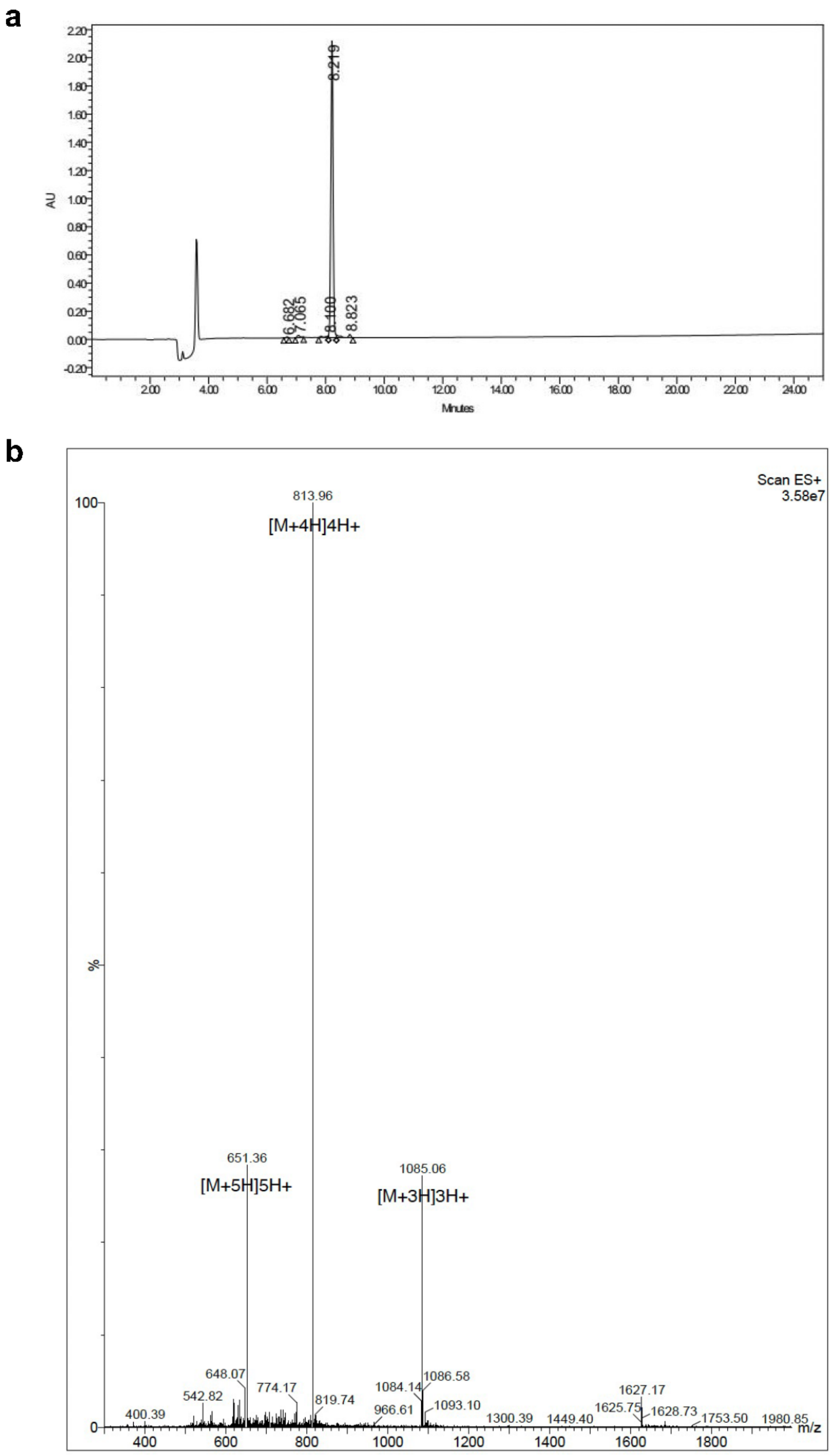


**Figure S51:** Validation of synthesized peptide 2-7. **a**, HPLC chromatography. **b**, Mass spectrometry.
